# Prenatal Environmental Stressors Impair Postnatal Microglia Function and Adult Behavior in Males

**DOI:** 10.1101/2020.10.15.336669

**Authors:** Carina L. Block, Oznur Eroglu, Stephen D. Mague, Chaichontat Sriworarat, Cameron Blount, Karen E. Malacon, Kathleen A. Beben, Nkemdilim Ndubuizu, Austin Talbot, Neil M. Gallagher, Young Chan Jo, Timothy Nyangacha, David E. Carlson, Kafui Dzirasa, Cagla Eroglu, Staci D. Bilbo

**Author notes:** Co-correspondence to: Staci Bilbo,; Cagla Eroglu,; Kafui Dzirasa.

## Abstract

Gestational exposure to environmental toxins and socioeconomic stressors are epidemiologically linked to neurodevelopmental disorders with strong male-bias, such as autism. We modeled these prenatal risk factors in mice, by co-exposing pregnant dams to an environmental pollutant and limited-resource stress, which robustly activated the maternal immune system. Only male offspring displayed long-lasting behavioral abnormalities and alterations in the activity of brain networks encoding social interactions. Cellularly, prenatal stressors diminished microglial function within the anterior cingulate cortex, a central node of the social coding network, in males during early postnatal development. Genetic ablation of microglia during the same critical period mimicked the impact of prenatal stressors on a male-specific behavior, indicating that environmental stressors alter neural circuit formation in males via impairing microglia function during development.

## Main Text

The incidences of neurodevelopmental disorders (NDD) (e.g. autism spectrum disorders, attention deficit disorder, learning disabilities) have been steadily increasing in recent decades suggesting a role for non-genetic environmental factors (*1, 2*). Furthermore, sex is a significant risk factor for these disorders, which have a strong male-bias (*3*).

Air pollutant exposure during pregnancy or the first year of life is one of the most consistent environmental risk factors for NDDs (*4-8*). However, the associations of single environmental agents, including air pollution, with NDDs have been relatively weak, and thus causality has been difficult to determine. Non-chemical stressors, such as limited resources or social support of the mother, can increase the vulnerability of the fetus to toxic exposures which could explain why certain populations are disproportionately affected (*9, 10*). In fact, neighborhood quality has recently been identified as a significant modifier of air pollution risk (*11*), suggesting environmental and social stressors synergize to increase vulnerability. But how these exposures alter fetal brain development and affect offspring behavior is largely unknown.

Inflammatory events during pregnancy such as maternal infection with bacteria or viruses have been shown to lead to maternal immune activation (MIA), which is linked to NDDs in offspring (*12-14*). Moreover, recent large-scale transcriptome-wide studies in post mortem brains of individuals diagnosed with an NDD have identified expression modules with enrichment of genes involved in neuroinflammatory function, with a particular dysregulation of microglial genes (*15-19*). Microglia are the primary immunocompetent cells of the brain and are exquisitely sensitive to perturbations of homeostasis, and thus may be poised as a primary responder to environmental insults. Microglia are also important regulators of activity-dependent synaptic remodeling during development (*20-23*), in which they prune inappropriate/weak synapses while sparing appropriate/strong connections. Importantly, aforementioned transcriptome studies have found that immune changes co-occur with gene enrichment modules affecting synaptic function. This suggests the possibility that neuroimmune changes during development could lead to aberrant synapse development due to altered microglial function.

Could the strong male-bias in NDDs, such as autism spectrum disorder (ASD), be linked to the differential response of fetuses to MIA? Notably, a recent analysis found that MIA was more common in male children with ASD compared to female children, suggesting that a sex difference in the response to maternal inflammation may be one mechanism which underlies increased male vulnerability (*24*). Furthermore, we and others have found strong sex differences in microglial development, maturation, and function, including an increased relative expression of microglial genes in male brains compared to females. Interestingly, the microglial genes that are enriched in male brains are also implicated in autism (*25, 26*). Together these data point to a mechanism by which sexually dimorphic microglial responses to prenatal stressors could lead to aberrant brain development primarily in males.

Here, we demonstrate that a combination of air pollution and maternal stress exposures during pregnancy activate the maternal immune system of mouse dams leading to altered synaptic and microglial development, persistent changes in brain circuit function, and long-lasting alterations in social and communication behavior primarily in male offspring.

### Prenatal exposure to air pollution and maternal stress induces maternal immune activation

To model a combination of chemical and social stressor exposures in mice, we exposed pregnant dams to intermittent chronic diesel exhaust particle (DEP) instillations to mimic air pollution. DEP is a primary toxic component of air pollution and a potent neuroinflammatory stimulus (*27-29*). Then, we applied a maternal stressor of resource deprivation during the last trimester of pregnancy by limiting the bedding and nesting material (i.e. DEP+MS condition) (Fig. 1A) (*30, 31*). Control dams received instillations of the vehicle solution (i.e. TBST) and were housed in standard cages with full nesting material (CON).

**Fig. 1.**
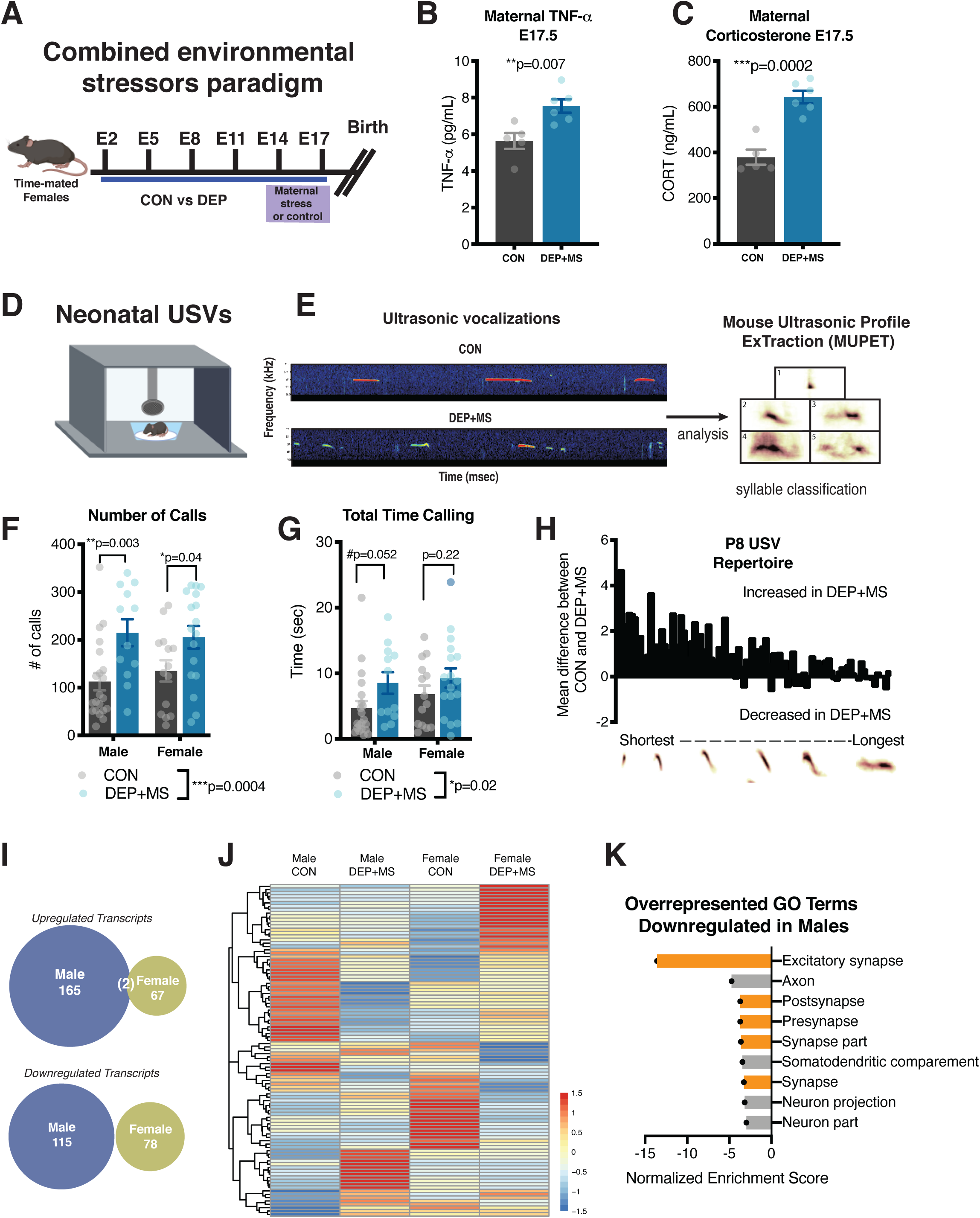
Combined prenatal stressors induce maternal immune activation and result in behavioral and transcriptome changes in offspring. (**A**) Schematic diagram of the experimental method. (**B-C**) Serum concentrations of TNFα and corticosterone 2 hours after CON or DEP instillation into pregnant dams at E17.5 (n= 5-6 mice/condition). (**D**) Neonatal USVs were collected by briefly separating the pup from the dam. (**E**) Representative USV spectrograms, USVs were analyzed using MUPET. (F-G) DEP+MS offspring emit more USVs and spend more time calling than CON mice (n=14-19 mice/condition/sex). (**H**) MUPET repertoire units organized from shortest to longest, DEP+MS offspring emit more calls that are preferentially shorter. (**I**) Number of DEG upregulated and downregulated in male and female PFC compared to CON (n=4 animals/condition/sex, male and female littermate pairs). (**J**) Heat map of DEG relative to control of the matched sex and show remarkably different patterns between male and female response to DEP+MS. (**K**) Synaptic genes are downregulated in male offspring. Unpaired t-tests (B,C), Two-way ANOVA with Holm-Sidak’s post hoc tests (F-G), Threshold free comparison of DEGs (I), Statistical overrepresentation analysis, Fisher’s Exact with FDR <0.05 (K). \*\*\**P*<0.001, \*\**P* < 0.01; \**P* < 0.05. Means ± SEM.

The combined environmental stressors, hereafter called DEP+MS, led to a strong induction of inflammatory cytokine tumor necrosis factor alpha (TNF-α) in the serum of exposed pregnant dams compared to CON at gestational/embryonic day E17.5 (Fig. 1B). A growing pre-clinical literature links MIA-induced cytokine expression in the pregnant dam to adverse outcomes in offspring. One of the other widely-adopted models of MIA, the maternal viral infection model (poly I:C), also increases TNF-α in dams (*32, 33*). In common with the poly I:C model, we also observed an increase in inflammatory cytokines IL-6 and IL-17a (Fig. S1A), but no change in anti-inflammatory IL-10 (Fig. S1B). Taken together, these results show that DEP+MS treatment leads to robust MIA in pregnant dams.

To further investigate and confirm the effects of maternal stress on DEP+MS dams, we measured the concentrations of maternal corticosterone (CORT), a stress hormone, in serum from gestational day E17.5 dams and found an increase in DEP+MS dams compared to CON dams (Fig. 1C), similar to that reported during postnatal nest deprivation (*31*). Importantly, CORT levels of CON dams were similar to baseline CORT levels of pregnant dams not receiving any treatment (Fig. S1C). These data show that our method of instillation alone is not sufficient to induce additional stress on pregnant dams. We did not observe any differences in pregnancy weight gain due to environmental stressors (Fig. S2B). Furthermore, we have previously found that prenatal DEP+MS does not alter maternal care, suggesting that alterations in DEP+MS pups are driven by MIA and not by fractured maternal care (*30*). Collectively, these findings show that DEP instillations which are combined with nest restriction during the last trimester of pregnancy leads to a combination of MIA and increased stress in these dams.

### Maternal immune activation via prenatal exposures affects neonatal offspring behavior and gene expression

To investigate the impact of combined prenatal stressors on offspring, we measured outcomes in the newborn pups. DEP+MS treatment did not alter litter size or sex composition (Fig. S2C). However, both male and female neonatal DEP+MS offspring had a significant decrease in weight compared to CON pups (Fig. S2D).

Neonatal mice, when briefly separated from the dam can emit ultrasonic vocalizations (USV). These USVs reflect an innate form of communication by the pups that elicits maternal care (*34, 35*). By recording the acoustic properties of USVs in neonatal offspring and analyzing the alterations in the pattern of calls (Fig. 1D-E), we tested whether DEP+MS alters neonatal communication. In wildtype C57BL/6J mice, peak USV production occurs at P8 (Fig. S3A), thus, to probe for developmental changes we recorded USVs from P7-P9. We found that pups from DEP+MS dams, emit more calls at P8 (Fig. 1F), mimicking phenotypes reported in other MIA models and a genetic mouse model of autism (*32, 36, 37*). Due to an *a priori* hypothesis regarding a potential sex difference, we performed post hoc analyses between sexes. Both males and females had a significantly increased number of calls (Fig. 1F), which was also accompanied by an increase in the total time calling (Fig. 1G). This increased number of calls in DEP+MS offspring was also evident at P7 but was no longer significantly different by P9 (Fig. S3B). These data suggest that prenatal DEP+MS is not inducing a developmental shift/delay in peak USV production, but instead leading to an abnormal calling behavior during the typical developmental window.

The properties and acoustic structure of USVs change across development and are context specific (*38, 39*). To further analyze these properties, we used Mouse Ultrasonic Profile ExTraction (MUPET) software, which applies machine learning to identify and cluster distinct repertoire/syllable units (*40*). Using MUPET across the 10,310 calls collected at P8, we extracted 80 distinct repertoire units from neonatal USVs (Fig. 1E, Fig. S3E). This analysis revealed that DEP+MS offspring emit more USVs across the whole repertoire of syllables with a preferential increase in shorter/less complex calls (Fig. 1H, Fig. S3E). Taken together, these results show that prenatal exposure to combined stressors lead to communication abnormalities in neonatal pups.

To gain molecular insights into the observed behavioral changes, we analyzed gene expression in the prefrontal cortices of P8 DEP+MS and CON male and female pups. We selected the prefrontal cortex because it is a brain region that is dysfunctional in many neurodevelopmental disorders and plays a critical role in regulating social and emotional behaviors (*41, 42*). Out of the 18,320 genes expressed, we identified 280 differentially expressed genes in DEP+MS males compared to CON males (Fig. 1I, 165 upregulated and 115 downregulated). In littermate females, only half as many genes (145) were differentially expressed compared to CON (Fig. 1I). Remarkably, there were only 2 differentially expressed genes that overlapped between males and females (Lcn2 and Dusp4, upregulated). Other than these two genes, the groups of differentially expressed genes were distinct in males and females (Fig. 1J). Thus, prenatal environmental insults result in distinct gene expression changes in male and female pup brains, suggesting sex-specific molecular effects of this manipulation on the offspring.

Even though only one of the top 10 upregulated genes (Lcn2) overlapped between male and female offspring datasets, there were several microglial enriched genes in the top 10 differentially expressed genes for both sexes (Fig. S4A-B). Furthermore, when we performed gene set enrichment analysis (GSEA), hallmark immune pathways were significantly enriched in both sexes (Fig. S5A-B).

On the other hand, there were no overlaps in hallmark downregulated pathways between sexes (Fig. S5C-D). The GSEA gene ontology for the cellular compartment revealed a male-specific downregulation of genes involved in synaptic structure and function in DEP+MS offspring (Fig. 1K). Taken together these RNA-Seq analyses show that, despite the fact that both male and female P8 mice display similar USV changes due to prenatal stressors, there are sexual dimorphisms in molecular and cellular responses. Here we captured changes in the expression of genes related to synapses (only males) and immune response (both sexes). Interestingly, brain gene expression studies from subjects with neurodevelopmental disorders show a similar down regulation in synaptic function genes and upregulation in immune response genes (*16-18*).

### Prenatal air pollution and maternal stress induce male-specific changes in adult socioemotional networks

Several neurodevelopmental disorders, such as ASD and ADHD, affect more males than females (*3*). Here we found behavioral changes in both sexes as neonates; however, the stark sex differences we found in PFC gene expression might be relevant to the increased susceptibility of males to develop sustained behavioral deficits due to prenatal insults (Fig. 1). Notably, previously, using the same DEP+MS model, we found a male-specific cognitive deficit in adulthood in a fear-conditioning task (*30*). Given these previous findings and combined with the downregulation of synaptic genes only in males, we next asked whether offspring have lasting changes in brain functional connectivity in a sex-specific manner.

In mouse models of neurodevelopmental disorders, a battery of behavioral tasks is utilized to measure social and emotional deficits (*43*). However, it is well known that many mouse models, including those with genetic construct validity, fail to exhibit measurable deficits in one or more of these commonly used assays (*44, 45*). Since altered brain network activity has been shown to be a sensitive measure of social deficits in ASD and preclinical models (*46-51*), we next investigated whether DEP+MS alters the network activity that underlies appetitive social behavior. To do so, we implanted 54 total CON and DEP+MS mice of both sexes with electrodes targeting eight brain regions (Fig. S6). We then recorded electrical oscillations concurrently from cortical and subcortical regions as mice performed an exploration task widely used to quantify appetitive social behavior (Fig. 2A). In this behavioral assay, experimental mice freely explore an arena that is divided into two chambers. One of these chambers contains a caged mouse (age- and sex-matched) of C3H background (highly social strain, non-aggressive), while the other chamber contains a caged inanimate object (Fig. 2A). During the test, the location of the experimental mice is tracked with video software, and the time spent in the proximity (4.98 cm) of either cage is used to calculate a preference ratio (interaction time_social_-interaction time_object_/ total interaction time). A ratio above 0 indicates a preference for the social stimulus versus object. This assay was repeated for each mouse for 10 sessions with at least one day off between sessions. A novel social and object stimulus were presented each day of testing, pseudorandomized to each side of the arena. In total, for each mouse, we collected 100 minutes of concurrent behavioral and electrical recordings. Overall mice displayed a preference for the social stimulus (preference score above 0; Fig. 2B); however, surprisingly, prenatal DEP+MS exposure resulted in a significant increase in the social preference scores of adult male mice compared to CON males (Fig. 2B). No difference in social preference was observed in female mice.

**Fig. 2.**
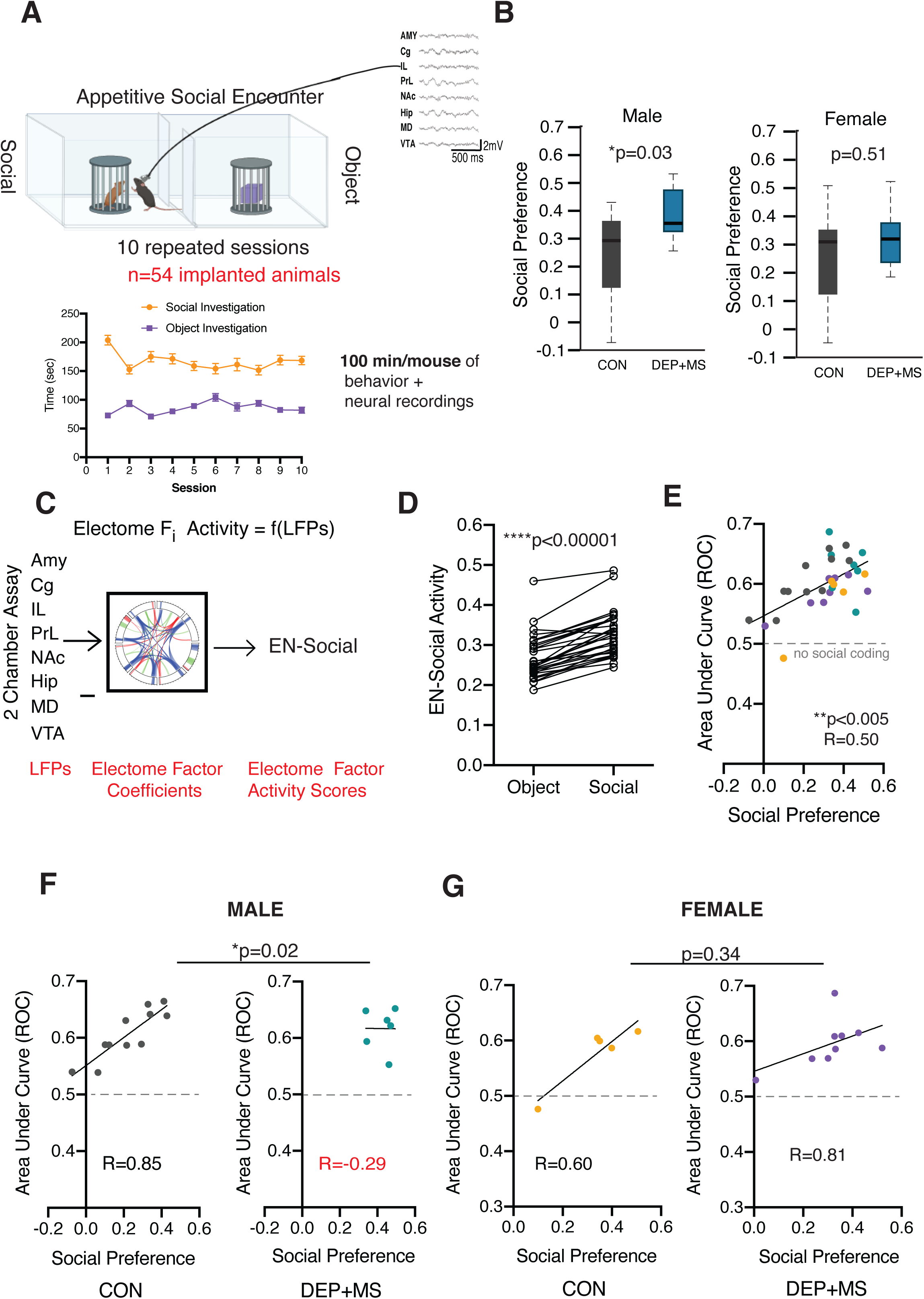
Electome network fails to signal social behavior in male offspring prenatally exposed to combined stressors. (**A**) CON and DEP+MS offspring were implanted with electrodes and underwent concurrent neural recording and in an appetitive social encounter task (top), mice were tested in 10 repeated session to collect 100 minutes of concurrent neural and behavioral data per mouse (bottom) (n=13-14 mice/condition/sex). (**B**) Prenatal DEP+MS exposure increased social preference in male (left), but not female mice (right). (**C**) LFP activity obtained during repeated two-chamber social interaction sessions was projected into the learned Electome factor coefficients. (**D**) Mice showed higher EN-Social activity during social compared to object interactions (n=31 animals). (**E**) The decoding accuracy of EN-Social activity signaled social preference across the population of implanted mice (n=31 animals). (**F-G**) Prenatal DEP+MS exposure disrupted the relationship between EN-Social activity and appetitive social behavior (social preference) in male, but not female mice (n=5-11 mice/condition/sex). Rank-sum test (B), Sign-rank test (D), Spearman’s correlation (**E**), Analysis of covariance with Box-Cox transformation, Spearman’s correlation (**F**,**G**). \*\*\*\**P*<0.0001, \**P* < 0.05. Means ± SEM.

Does this behavioral change in DEP+MS males reflect altered social processing or simply an increase in normal social appetitive drive? To address these questions, we utilized a model that was generated by Mague *et al*., 2020 (*52*) based on data collected from a untreated group of C57BL/6J WT mice performing the identical social preference task described above (hereafter referred to as EN-social, Fig. S7). This EN-social network was discovered using a machine-learning approach that utilizes a discriminative cross spectral factor analysis based in non-negative matrix factorization (dCSFA-NMF, (*53*)). In brief, this method integrates local field potential (LFP) activity from multiple brain regions with concurrent behavior (Fig. 2C). The model features utilize LFP power, which is a neural correlate of cellular population activity and synaptic activity within brain regions, and LFP synchrony is a neural correlate of brain circuit function between brain regions, quantified by how two LFPs across frequencies correlate over a millisecond timescale. The model also uses spectral Granger causality (*54*), which is used here as an estimate of information transfer within a circuit. Each of these features is resolved from 1-56 Hz. Critically, the activity of EN-Social predicts an animal’s investigation of the social stimulus and correlates with an individual animal’s social preference, reflecting the rewarding nature of social encounters on a mouse-by-mouse basis (*52*). Notably, the correlation of the EN-social network activity with social preference was found to be disrupted in a genetic mouse model of autism, despite a lack of change in social preference behavior (*52*).

Using this social brain state as a framework, we overlaid the brain activity of our 54 implanted mice onto this EN-social network. Overall, our experimental mice (both DEP+MS and CON) displayed higher EN-Social network activity when they were interacting with the social stimulus versus the object (Fig. 2D), providing further confirmation that this proposed network is encoding a social brain state. Furthermore, there was a direct correlation between the social preference scores of mice and the extent to which EN-Social activity encoded the social interaction (Fig. 2E), demonstrating that this network accurately reflected the rewarding component of social behavior across the new groups of mice. However, when we performed within-sex comparisons between the DEP+MS and CON groups, we found that this brain activity-behavior relationship was disrupted in the male DEP+MS mice (Fig. 2F), but not female mice exposed to DEP+MS or the CON mice of both sexes (Fig. 2F-G). In other words, in CON males and females and DEP+MS females, higher social preference directly correlated with increased activation of the EN-social network in response to social encounters; however, this correlation was abolished in DEP+MS male mice. These findings are strikingly similar to those found in a genetic model of ASD (ANK2 knockout mice) as mentioned above; in which the EN-social network activity-social preference relationship was also disrupted (*52*).

Taken together, these findings reveal that prenatal DEP+MS exposure causes MIA and leads to long lasting deficits in socioemotional encoding and altered behavior only in males. Importantly, the incidence of neurodevelopmental disorders is higher in males than females, and a recent study highlighted that a history of MIA is significantly higher in mothers of male children diagnosed with ASD compared to females (*24*). Our DEP+MS model captures this male-specific vulnerability to MIA, thus providing an important model to study the cellular and molecular mechanisms.

### Prenatal environmental toxins impair postnatal thalamocortical synapse development and microglial pruning in the anterior cingulate cortex

The anterior cingulate cortex (ACC), a critical node of EN-Social (Fig. S7), is functionally linked to communication outcomes (*55, 56*). Furthermore, genetic manipulation of synaptic function in the ACC is sufficient to drive social deficits in behavior (*57*). Therefore, since adult DEP+MS mice exhibited altered EN-Social function and social behavioral changes, we hypothesized that prenatal DEP+MS exposure alters circuit formation with the ACC during a critical window of synaptic development. In particular, we focused on the early postnatal period when USVs are produced (P6-P10) because we found that DEP+MS exposure disrupts USVs.

The ACC receives excitatory synaptic inputs from several cortical and subcortical areas including the thalamus. Thalamocortical synapses (TC), which are formed from thalamic axonal inputs onto the cortical dendrites, can be identified by the juxtaposition of vesicular glutamate transporter-2 (VGlut2) positive presynaptic terminals and PSD-95 positive postsynaptic densities (Fig. 3A-B) (*58*). Importantly, thalamocortical pathways are critical for relaying subcortical sensory information to the cortex, and hypoconnectivity of these pathways has been reported in ASD and is thought to underlie sensory processing issues in patients (*59-61*). Moreover, in our transcriptome analyses, we found a DEP+MS male-specific downregulation of excitatory synapse genes at P8, a time point corresponding to heightened TC synaptogenesis. Therefore, we wondered whether TC synapse development is affected in these animals.

**Fig. 3.**
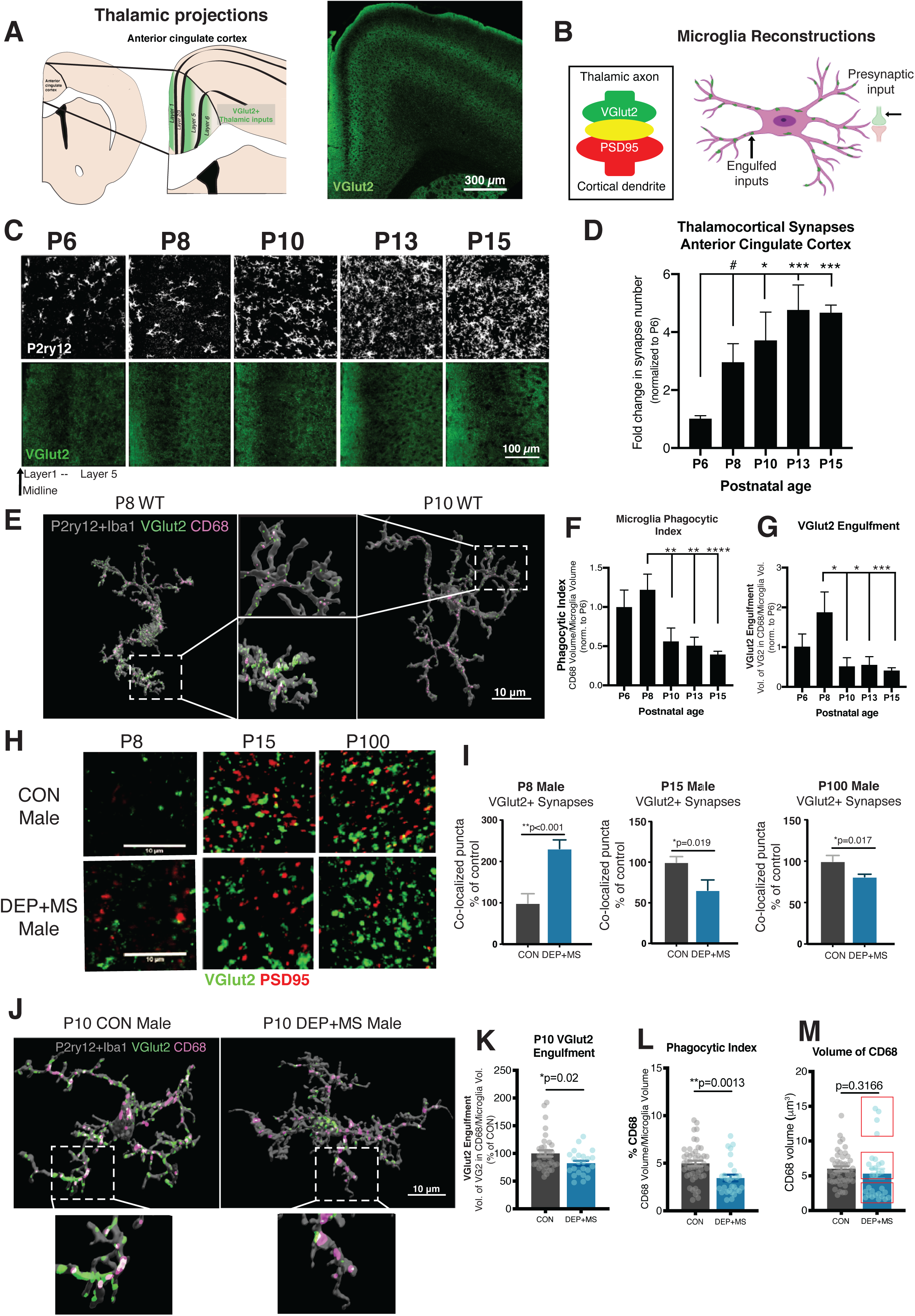
The second week of postnatal development is a critical window for ACC development. (**A**) Thalamic inputs into the ACC are pseudo laminated and can be labeled with VGlut2. (**B**) Synapses can be quantified by the apparent co-localization of VGlut2 and PSD95 (left), 3-D cell reconstructions allow for the visualization of internalized inputs inside microglia (right). (**C**) Representative images of VGlut2 and P2ry12 (TC input and microglia) in the ACC of WT mice from P6-P15. (**D**) Quantification of VGlut2 synapses in layer 1 of the ACC (n=3 replicates/mouse, 4 mice/age (2 males, 2 females), 300 images analyzed, data normalized to P6). (**E**) Representative 3-D reconstructions of microglia from WT P8 and P10 male mice. Lysosomes (CD68) and TC inputs (VGlut2) can be visualized inside microglia and quantified. (**F-G**) Quantification of CD68 and VGlut2 volume internalized in microglia, (n=6-7 cells/mice, 2 male mice/age, 75 microglia cells reconstructed, all values normalized to cell volume and engulfment normalized to P6). (**H**) Representative images of TC synapses (co-localized VGlut2 and PSD95) in the ACC of P8, P15 and P100 CON and DEP+MS male mice. (**I**) TC synapse numbers are increased at P8 and decreased at P15 and P100 in DEP+MS male mice compared to CON males (n=3 replicates/mouse, 3 mice/condition, 270 images analyzed). (**J**) Representative surface rendered microglia from the ACC of P10 CON and DEP+MS male offspring labeled with P2ry12+Iba1, VGlut2 and CD68. (**K-L**) DEP+MS male microglia engulf fewer VGlut2 inputs and have significantly less lysosomal content (phagocytic index). (**M**) No significant difference in mean CD68 volume, but CD68 volume distribution is altered in DEP+MS male microglia, red boxes. (n = 3–4 replicates/mouse, 3 mice/condition, 110 total images analyzed). One-way ANOVA with Holm Sidak’s post hoc tests (D, F, G), Nested t-test (I, K-M). \*\*\*\**P*<0.0001, \*\*\**P*<0.001, \*\**P* < 0.01; \**P* < 0.05. Means ± SEM.

In early postnatal brain development, an exuberant period of synaptogenesis is closely followed by and overlaps with a period of synaptic pruning, where weak or unnecessary synapses are eliminated (*21*). Disruption of either of these processes can profoundly affect both circuit formation and function. One mechanism of synaptic pruning and circuit refinement occurs via the activity-dependent engulfment of synaptic material by microglia (*21, 22*). Microglia selectively phagocytose (or trogocytose) presynaptic structures, which are degraded through trafficking to lysosomal compartments (Fig. 3B) (*62*). Increased microglial reactivity has been reported in several brain regions in ASD patients, although causality has been difficult to determine (*63, 64*). In our transcriptome analyses, we identified an enrichment of microglial genes and an upregulation of pathways involved in immune function, alongside a downregulation of synaptic genes in males (Fig. S4A-B; Fig. S5A-B; Fig 2K), suggesting a link between the two.

Although synapse formation and elimination have been described in other brain regions including the visual thalamus and barrel cortex (*21, 65*), the exact timing of TC synapse formation and elimination in the ACC are unknown. To first characterize the normal pattern of synaptic development in the ACC, we quantified synapse density and microglial engulfment during postnatal ages (P6-P15) in a naïve group of WT mice (Fig. 3B-C). During this time frame, we found a five-fold increase in TC synapse density (Fig. 3D). Furthermore, we found that the ACC becomes increasingly more organized and pseudo-laminated (Fig. 3C; Fig. S8A). These data reveal that, between P6 and P15, concurrent synapse formation and elimination are taking place in the ACC, leading to an increase in structural organization.

In our assessment of normal development in WT mice, we found that microglia undergo a period of rapid development, dramatically increasing in density and coverage between P8 and P10 (Fig. 3C). To assess whether this period coincides with peak alterations in synapse elimination, we performed IHC using a cocktail of two pan microglia markers (Iba1, Ionized calcium-binding adaptor protein-1; P2yr12, purinergic receptor), and antibodies for microglia lysosomes (CD68) and TC inputs (VGlut2). Synaptic engulfment can be quantified by 3-D reconstructions of microglia to visualize internalized VGlut2 and lysosomal compartments within the cell (Fig. 3B, right; Fig. 3E). We found that lysosomal content/phagocytic activity (CD68 in microglia) was highest at P8 and was significantly diminished beginning at P10 (Fig. 3F). Furthermore, quantification of VGlut2+ inputs within microglia which were also co-localized with CD68+ lysosomes, revealed that TC synapse engulfment also peaks at P8 and is largely completed by P10 (Fig. 3G). These data show that the period between P8 and P10 represents a critical window of microglia engulfment of TC synapses. Therefore, microglial dysfunction at this timepoint in development can have lasting consequences for brain connectivity.

Next, to determine whether TC synaptic structures were altered in the ACC of DEP+MS mice, we quantified the number of TC synapses in male and female offspring at P8, during the period of synaptogenesis when communication behavior is highly disrupted, at P15 when TC synapses reach their peak density, and in adulthood (>P60) when synapse density is relatively stable (Fig. 3H-I, Fig. S8C). At P8, we found a significant increase in the number of TC synapses in DEP+MS males, but not females (Fig. 3H-I, Fig. S8C). At P15, the timepoint of peak synapse abundance in WT mice, both male and female DEP+MS offspring had a significant reduction in TC synapse number compared to CON. However, this decrease in TC synapse number only persisted into adulthood in male offspring (Fig. 3H-I, Fig. S8C). Importantly, the density and distribution of neurons (NeuN+), astrocytes (Sox9+Olig2-) and oligodendrocytes(Sox9-Olig2+) were unchanged in DEP+MS offspring (Fig. S9, S10A-C). Taken together, these data show that DEP+MS males have an overgrowth of TC synapses at P8; however, this initial overgrowth is rapidly lost by P15 and results in a reduction in TC connections in the ACC, a phenotype that persists into adulthood only in males. Interestingly, in individuals with ASD, TC pathways are found to be hypo-connected, which may underlie sensory processing issues (*59*).

To determine whether rapid atrophy of TC synapses in males can be attributed to enhanced and prolonged microglial engulfment during this period (P8-P15), we next investigated whether DEP+MS male microglia had alterations in TC synapse engulfment at P10, when peak engulfment is completed in normal development. Surprisingly, we found that DEP+MS microglia were engulfing significantly less synapses (Fig. 3J-K). In line with these data, phagocytic activity was also significantly diminished in microglia from DEP+MS males (Fig. 3L). Intriguingly this change was not due to a significant reduction in volume of CD68 (Fig. 3M); but instead we found that there was a significant difference in the distribution of CD68 volume, with subsets of high and low CD68 expressing cells (S11A-B), suggesting that there may be additional functional changes in male DEP+MS microglia. In sum, microglia from male DEP+MS offspring engulf fewer TC synapses at P10, are less phagocytic and have alterations in the distribution of CD68. Taken together, our results indicate two unexpected phenomena. First, contrary to our initial hypothesis, microglia from DEP+MS males have diminished phagocytic function overall and secondly, this diminished function is only strongly affecting a subset of microglia.

### Prenatal exposure to environmental toxins leads to an increase in functional heterogeneity of male microglia

In DEP+MS males, we found diminished TC input engulfment by microglia at P10, suggesting that the reduction in TC synapse density by P15 cannot be attributed to enhanced microglial engulfment of synapses. Next, we wondered whether the atrophy of TC inputs could be attributed to an increase in microglia cell density. To investigate this possibility, we quantified microglia cell density in the ACC at P8, P15 and P25 in CON and DEP+MS offspring. To do so, we performed immunohistochemistry using antibodies against P2ry12 and Iba1 and independently labeled these antigens by using separate fluorophores. Microglia were identified by the presence of either P2ry12 and/or Iba1 signal, which also co-localized with the nuclear marker DAPI (Fig. S12A). There were no significant differences in the total density of microglial cells between CON and DEP+MS male offspring across all ages (Fig. S12B), showing that changes in microglia numbers are not likely to underlie alterations in synaptic development. Intriguingly, while the majority of microglia express high levels of both Iba1 and P2ry12, we identified a subset of cells which express high levels of one marker and not the other (Fig. 4A-B).

**Fig. 4.**
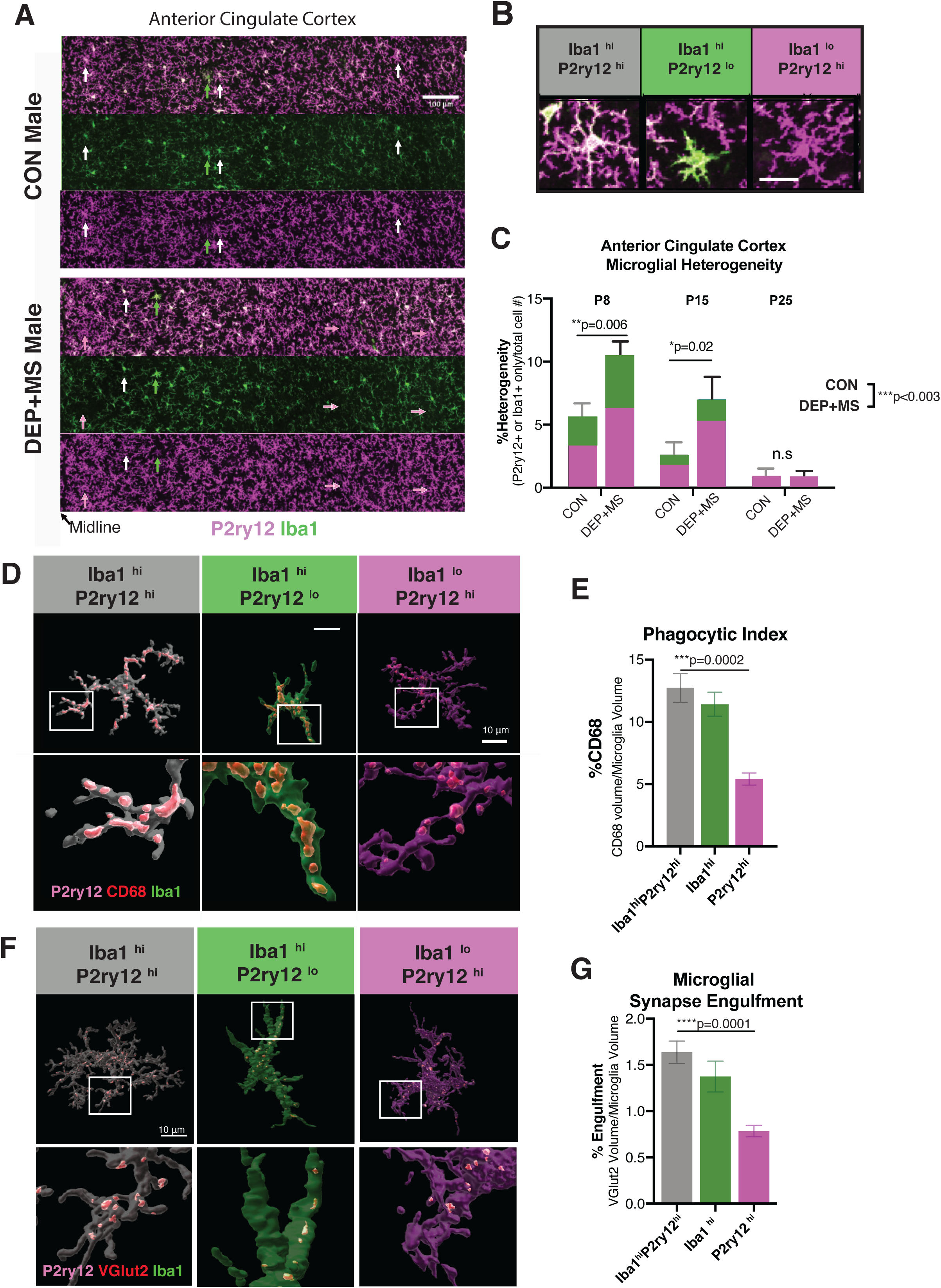
Microglial development and synaptic engulfment are altered in male offspring prenatally exposed to combined stressors. (**A**) Representative images of microglia labeled with P2ry12 and Iba1 in the ACC of P15 male offspring, arrows highlight microglia differentially labeled by P2ry12 and Iba1 (white-both high, green-Iba1 high, magenta-P2ry12 high). (**B**) Representative images of microglia with heterogeneous levels of expression of Iba1 and P2ry12 in the developing cortex. (**C**) Quantification of microglial heterogeneity across development in male offspring in the ACC, (n=3 replicates/mouse, 3-4 mice/condition/age, >6,000 cells counted). (**D**) Representative Imaris reconstructions of microglia at P8 with lysosomal content (CD68) in microglial subtypes, images acquired at 30x. (**E**) P2ry12^hi^ microglia are less phagocytic than Iba1^hi^P2ry12^hi^microglia cells (n=5-10 cells/mouse/cell subtype, n=3 animals/condition, total of 120 cells analyzed). (**F**) Representative Imaris reconstructions of microglia at P8 with internalized VGlut2 in microglial subtypes, images acquired at 60x. (**G**) P2ry12^hi^ microglia engulf fewer synapses compared to Iba1^hi^P2ry12^hi^ cells (n= 5-12 cells/mouse/cell subtype, n=3 mice/condition, total of 168 reconstructed cells). Two-way ANOVA condition x age, with Sidak’s post hoc test (C), Nested One-way ANOVA with Holm-Sidak’s post hoc test (E, G). \*\*\*\**P*<0.0001, \*\*\**P*<0.001, \*\**P* < 0.01; \**P* < 0.05. Means ± SEM.

P2ry12 and Iba1 each have important roles in microglia function and are known to be expressed at varying levels within microglia reflecting different cellular states. For example, P2ry12, which is a G protein-coupled purinergic receptor, is necessary for ADP/ATP-mediated chemotaxis and microglial process extension to sites of brain injury (*66, 67*). Furthermore, pharmacological block or deletion of P2ry12 during development leads to reduced critical period plasticity (*68*). Moreover, immune activation severely diminishes P2ry12 expression in microglia (*69, 70*). On the other hand, Iba1, an ionized calcium-binding adaptor protein, is known to modulate actin reorganization and facilitate cell migration as well as phagocytosis by microglia (*71*). Iba1 has also been known to increase with some inflammatory insults (*71-73*).

In both CON and DEP+MS male offspring, we observed three types of microglia with respect to their differential expression of Iba1 and P2ry12. The majority of microglia highly expressed both Iba1 and P2ry12 (Fig. 4B, Iba1^hi^P2ry12^hi^, left). On the other hand, some microglia expressed high levels of Iba1, but low levels of P2ry12 (Iba1^hi^P2ry12^lo^). These cells were often found in layers 2/3 and had a strikingly different morphology compared to Iba1^hi^P2ry12^hi^ (Fig. 4A and Fig. 4B, middle). We also found cells which were expressing high levels of P2ry12 and low levels of Iba1. These Iba1^lo^P2ry12^hi^ cells were often found in deeper layers, and their morphology was indistinguishable from the Iba1^hi^P2ry12^hi^ microglia (Fig. 4A and Fig. 4B, right). Although these different types of microglia were more common in specific layers, they were often neighbored by the predominant microglia subtype; Iba1^hi^P2ry12^hi^.

To determine if prenatal DEP+MS exposure modifies the relative abundance of these microglial subtypes, we quantified the percentage Iba1^hi^P2ry12^lo^ or Iba1^lo^P2ry12^hi^ microglia, which we termed here as microglial heterogeneity. Early in development (P8-15) microglial heterogeneity was higher in both CON and DEP+MS offspring compared to a later developmental time point, P25 (Fig. 4C). This observation suggests that the presence of these microglia subtypes does not reflect a pathological brain state, but rather part of a normal developmental process. However, microglial heterogeneity was strikingly enhanced in DEP+MS male offspring ACCs compared to CON, both at P8 and P15 (Fig. 4C). Recent single-cell analyses of microglia across development have revealed that these cells are molecularly highly heterogenous during very early postnatal ages (*74-77*). Our data indicate that subtypes of microglia with varied expression of two important microglial proteins P2ry12 and Iba1 are present during early ACC development and that prenatal DEP+MS insult increases the relative abundance of heterogeneity.

At P8 when microglial heterogeneity is high, microglia are also actively pruning VGlut2 synapses (Fig 3G). Therefore, we next tested if the three subtypes of microglia we identified (Fig 4B) differ in their phagocytic function and their ability to engulf VGlut2 synapses. To do so, we labelled the ACC with antibodies against Iba1, P2ry12 and CD68 and, using Imaris, we reconstructed a total of 120 Iba1^hi^P2ry12^hi^, Iba1^hi^P2ry12^lo^ and Iba1^lo^P2ry12^hi^ microglia (Fig. S13A-C). Next, we quantified the lysosomal content (CD68 volume/microglia volume) within these distinct microglial subtypes as a proxy for the phagocytic activity of these cells. There were no significant differences in the phagocytic activity of Iba1^hi^P2ry12^lo^ cells compared to the Iba1^hi^P2ry12^lo^. However, Iba1^lo^P2ry12^hi^ cells had significantly lower (∼50% less) CD68 content compared to the prevalent Iba1^hi^P2ry12^hi^ microglia type (Fig. 4D-E). This pattern of reduced CD68 content was present in both CON and DEP+MS microglia (Fig. S14A) but did not differ significantly between groups (Fig. S14B-C). These results indicate that Iba1^lo^P2ry12^hi^ cells have severely diminished phagocytic activity compared to the other two subtypes. These data suggest that diminished phagocytic activity could alter microglia’s ability to eliminate synaptic inputs.

To investigate whether the reduction in CD68 was linked to interactions with TC synapses, we labelled the ACC with antibodies that recognize Iba1, P2ry12 and VGlut2 and using Imaris, reconstructed a total of 168 Iba1^hi^P2ry12^hi^, Iba1^hi^P2ry12^lo^ and Iba1^lo^P2ry12^hi^ microglia (Fig. 4F). Next, we quantified volume of VGlut2+ TC inputs within these distinct microglial subtypes (VGlut2 volume/ cell volume). Compared to the Iba1^hi^P2ry12^hi^ microglia, Iba1^hi^P2ry12^lo^ cells had a small (∼20%) but trending decrease in engulfed TC inputs. On the other hand, Iba1^lo^P2ry12^hi^ microglia engulfed significantly fewer (∼50%) TC inputs compared to Iba1^hi^P2ry12^hi^ cells (Fig. 4F-G). Collectively, these results show that Iba1^lo^P2ry12^hi^ cells have diminished lysosomal content and engulf fewer TC synapses. Importantly, these functional differences between the three microglial subtypes are present in both CON and DEP+MS male offspring brains (Fig S15A), showing that prenatal insults do not affect the per cell functional responses. Instead the specific subsets of cells (Iba1^lo^P2ry12^hi^) are more abundant in DEP+MS offspring. Intriguingly, when we compared engulfment of microglia subsets, we found no group differences in synaptic engulfment between CON and DEP+MS male microglia at precisely P8 (Fig S15 C). Thus, the net impairment in synapse elimination that was observed at P10 (Fig 3K) is driven by the cumulative increase in less phagocytic microglia types. In summary, these results indicate that the atrophy of TC synapses in DEP+MS male brains is not due to an exuberant pruning function of microglia. Instead we found evidence of a loss of normal microglial function in DEP+MS male ACCs, reflected by increased heterogeneity and a net reduction in the ability of these cells to phagocytose synapses.

### Elimination of microglia during a critical postnatal period causes enduring communication deficits similar to those found in DEP+MS males

In males, prenatal DEP+MS insult leads to a transient diminishment of microglia function during an early postnatal period (Fig. 4), and also results in enduring behavioral alterations (*30*) (Fig. 2B). Could the transient loss-of-microglia-function during early development (P8) contribute to the enduring behavioral effects we observe in DEP+MS males? To ask this question, we first studied an adult male-specific communication behavior in CON and DEP+MS offspring. In this assay, males with sexual experience emit USVs when introduced to sexually receptive females (i.e. female in estrus or proestrus). These USVs, also known as the “courtship songs”, are distinct from neonatal USVs, in that they are more complex and critical for proper mating behavior and drive female mate choices (*78-80*). We performed our prenatal treatment on dams and raised offspring into adulthood. Mating songs from adult CON and DEP+MS males were recorded in this courtship assay. Female mice rarely vocalize during direct male-female interactions; thus USVs collected in this paradigm can be attributed to experimental male animals (*80*) (Fig. 5A-B).We found that male DEP+MS offspring exposed to a female in estrus are able to respond to a socially-relevant stimulus and similar to CON males emit a courtship song. We found no significant differences in the number of calls emitted between groups, but when we analyzed the acoustic properties of the calls, the individual calls were significantly shorter, thus resulting in a significant reduction in the time spent vocalizing (Fig. 5C-D; S16A). We found no significant differences in the frequency (kHz) of the emitted calls (Fig. S16B). Importantly, female mice, when presented with a choice, prefer courtship songs that are longer and more complex (*78*), suggesting DEP+MS males produce less competitive songs.

**Fig. 5.**
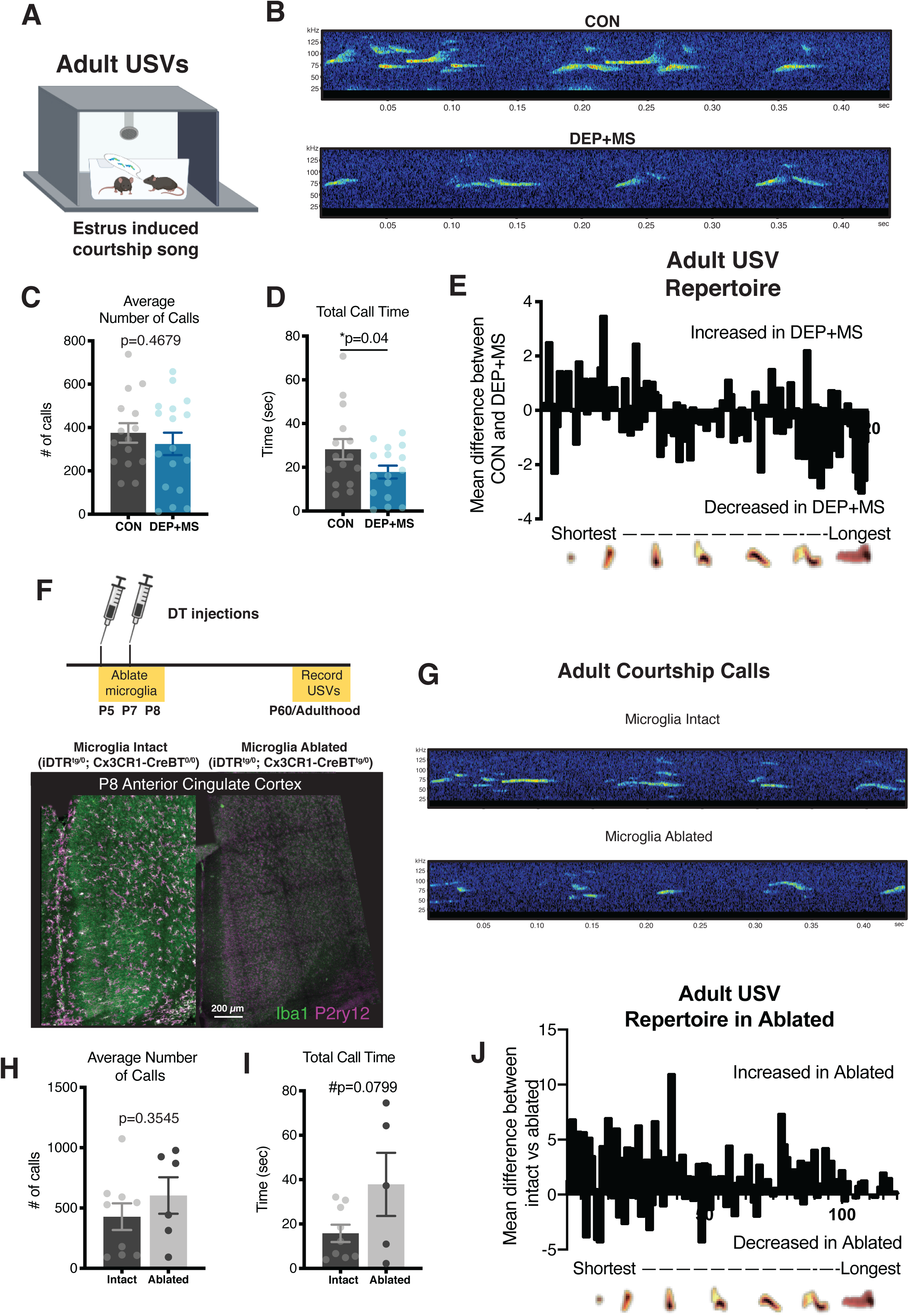
Neonatal microglial ablation mimics adult communication deficits in male offspring prenatally exposed to combined stressors. (**A**) USVs were collected from adult male mice by introducing a sexually receptive female. (**B**) Representative spectrograms of USVs from CON vs DEP+MS adult males. (**C-D**) No significant difference in number of calls, but adult male offspring spend less time calling than CON and emit USVs that are shorter in length (n=15-17 mice/condition). (**E**) MUPET repertoire units organized from shortest to longest, pattern of calls is shifted towards less complex in adult DEP+MS male offspring. (**F-G**) Cx3cr1-CreBT-inducible DTR expression, renders microglia sensitive to diphtheria toxin, for elimination. 80 ng of diphtheria toxin was administered to iDTR^tg/0^; Cx3cr1-CreBT^0/0^ and iDTR^tg/0^; Cx3cr1-CreBT^tg/0^ mice at P5 and P7, effectively eliminating 98% of microglia at P8 in mice expressing cre. (**G**) USVs were collected in adulthood. Representative spectrograms from males with neonatal ablation of microglia or microglia intact. (**H-I**) Neonatal ablation of microglia does not alter call number, but there is a trend towards an increase in total time calling (n=8-11 mice/genotype). (**J**) MUPET repertoire units organized from shortest to longest, microglia elimination shifts pattern of vocalization towards less complex calls. Unpaired t-test (C, D, H, I), Unpaired t-test (G, H) \**P* < 0.05. Means ± SEM.

To further investigate song complexity and to identify distinct repertoire units, we performed MUPET analyses on the ∼33,000 calls collected and identified 120 distinct repertoire units in both CON and DEP+MS males (Fig. S16C). These analyses revealed that DEP+MS male offspring have a preferential increase in calls that are short and less complex and a reduction in longer and more complex call types (Fig. 5E), once again suggesting DEP+MS male courtship songs are less competitive. Furthermore, this observed reduction in call complexity in adult DEP+MS males is similar to changes in call complexity observed in neonatal DEP+MS offspring that also preferentially emit calls that are shorter and less complex (Fig. 1H). Collectively these data reveal that changes in vocalization persist into adulthood in DEP+MS offspring, allowing us to test the hypothesis that a loss-of-function in microglia during the critical window of TC synapse development will produce a similar behavioral outcome.

To investigate whether microglia loss-of-function plays a causative role in the observed behavioral and brain dysfunction, we targeted and eliminated microglia during the early postnatal period of synapse development in the ACC. To do so, we used a genetic depletion strategy, which enables Cre-mediated expression of the diphtheria toxin receptor (DTR) specifically in CX3CR1+ cells by crossing commercially available iDTR mice to Cx3cr1-CreBT mice (Fig. S17A) (*81, 82*). Because mice do not express the DTR, they are naturally resistant to diphtheria toxin (DT); however, upon the Cre-induced expression of DTR, cells will undergo apoptotic cell death when DT binds to the receptor (Fig. S17A-B) (*83*). Furthermore, DT crosses the blood-brain-barrier, thus allowing for the ablation of CNS cells, such as CX3CR1+ microglia (*83*).

While pharmacological depletion strategies, such as CSF1R inhibitor molecules, are commonly used for depleting microglia, depletion via CSF1R inhibitor can take up to two weeks and requires daily oral administration which is disruptive for neonatal animals and their mothers (*84*). Our transgenic elimination strategy allows for a rapid depletion with minimal disruptions to pups; furthermore previous studies have demonstrated that genetic elimination results in microglial repopulation from a CNS-resident internal pool without contribution from peripheral or infiltrating cells (*85*). In our previous analyses we found that only a subset of microglia have a normal loss of function. While we recognize that elimination of all microglia is far harsher, this manipulation enables us to casually link microglia loss of function to the behavioral outcomes we have observed.

Thus, to investigate whether microglia loss-of-function early in development could modify adult communication behavior, we depleted microglia at early postnatal ages during the period we identified where rapid TC synapse formation and elimination are occurring (∼P8). To do so we crossed Cx3cr1-CreBT^tg/0^ dams with iDTR^tg/tg^ males to generate offspring negative and positive for Cre, all carrying a single copy of the iDTR transgene. All neonatal offspring (negative and positive) for Cre (Cre ^0/0^, iDTR^tg/0^, microglia intact; Cre ^tg/0^, iDTR^tg/0^, microglia ablated) received DT injections at P5 and P7, which eliminated ∼98% of microglia cells by P8 in mice expressing the Cre transgene (Fig. 5F, Fig S17A-B). By P30 microglia have fully repopulated the brain; however, expression of homeostatic marker P2ry12 remained low in the cortex, thus driving a type of heterogeneity not typically observed at this age (Fig. S18A).

Male offspring underwent microglia ablation (or intact control) during the early postnatal period and were raised into adulthood where they were tested in an estrus-induced courtship assay (Fig. 5G). Similar to our findings in DEP+MS males, we found no significant differences in the number of USVs emitted in our intact vs ablated condition (Fig. 5H-I). However, when we performed MUPET analyses of song complexity, we found a preferential increase in calls that are shorter and less complex (Fig. 5J). While not identical, these findings largely mimic the observed differences in CON vs DEP+MS male songs. Thus, microglia depletion during the critical window of synapse elimination (around P8) has lasting effects on behavior that are similar to prenatal DEP+MS treatment. These data reveal that microglia loss of function either due to MIA during pregnancy or depletion during early postnatal development, profoundly alters communication behavior in male offspring.

## Discussion

Immune dysfunction in pregnant mothers is increasingly implicated in the pathogenesis of NDDs and is strongly linked to male offspring-specific behavioral outcomes (*24*). Here we show that prenatal co-exposure to two highly prevalent environmental factors, air pollution and maternal stress, is sufficient to induce maternal immune activation and significantly increase stress hormones in pregnant mice. We found that both male and female offspring born to these dams had altered USVs as neonates. Surprisingly however, gene expression changes in the prefrontal cortices of neonatal mice were sexually dimorphic, and behavioral alterations only persisted into adulthood in male mice. These data indicate that prenatal environmental insults result in a distinct response in developing male brains compared to females.

Similarly, we found that functional activation of the social brain network was disrupted only in adult DEP+MS male offspring. Specifically, the relationship between social investigation and activation of this circuit was no longer behaviorally relevant in DEP+MS males. A possible interpretation of this difference is that during development synaptic circuits that encode social interactions form differently in male mice which were exposed to DEP/MS as fetuses. In agreement with this possibility, in the ACC, a critical brain region within this network, in DEP+MS male mice we found an early overabundance of TC synapses by the end of the first postnatal week; however, by P15, this phenotype was reversed suggesting that altered synaptic development leads to persistent brain miswiring in social circuits. One mechanism by which these synapses might be lost is due to excessive synapse elimination by microglia cells, which were also implicated in our sequencing analysis. Increased microglial activation has been described in several neurological disorders, thus overactivation of microglia is thought to severely impact brain health. However, investigation of microglial engulfment revealed that the loss of TC synapses we observed cannot be readily explained by excess synapse elimination by microglia. Instead, microglia from DEP+MS males were less phagocytic and engulfed fewer synapses. Moreover, we discovered ACC microglia have a developmentally regulated antigenic and functional heterogeneity. This heterogeneity was strongly enhanced in males prenatally exposed to DEP+MS, leading to the overabundance of one specific subtype of microglia with severely diminished phagocytic activity that eliminated fewer TC synapses. Taken together our findings indicate that over-pruning of TC synapses by microglia does not underlie the structural and functional changes we observed in DEP/MS male ACCs. Our results however, are in line with a loss-of-function phenotype with microglia, consistent with previous findings that if microglial phagocytosis levels are not finely tuned to clearance requirements within a given brain region, this can result in aberrant brain development and altered behavior (*86*). Future studies investigating the origins and etiology of the different microglial subtypes in developing brains may provide important information about how microglial dysfunction may impact the pathogenesis of NDDs.

In disorders such as autism, early overgrowth of synaptic connectivity is often followed by an atrophy, but the mechanism of the atrophy remains unknown. Here we see a similar phenotype, but we know that the loss of TC synapses cannot be explained by exuberant synapse elimination by microglia; thus how these synapses are lost remains unclear. Synapses can be removed via multiple mechanisms including astrocyte-mediated elimination (*87*). Astrocytes are macroglial cells which mediate synapse formation, functional maturation, and elimination. In particular, a number of studies revealed important roles for astrocytes in controlling TC synapse formation and maturation (*88-90*). Furthermore, neuroimmune insults are known to trigger different reactive profiles in astrocytes which may also be happening in DEP+MS male brains, potentially underlying synapse loss and circuit dysfunction (*91*). Thus future investigations are needed to determine if astrocytes or other brain cell types are also involved in the dysfunctional synaptic development that we observed in DEP+MS male brains. Similarly, future studies investigating the cellular and molecular mechanisms underlying male and female responses to MIA are needed to further our understanding of why male brains are more vulnerable and/or female brains are protected (*24*).

In conclusion, our combined stress model has allowed us to rigorously investigate the mechanisms underlying abnormal brain development in response to these pervasive environmental factors. Our findings clearly indicate that environmental pollutants can synergize with social stress in pregnant mothers and cause MIA, which we found to have specific long-term effects on the development and function of male brains. This is particularly concerning, now more than ever, because ongoing climate change caused by increased economic activity, and a reduction in environmental enforcements have led to a rapid worsening air quality in recent years. Heightened air pollution is likely to differentially synergize with the social stressors causing further disparities in the well-being of future generations. Therefore, our findings provide an important first step towards revealing the non-genetic causes for NDDs so that preventative and therapeutic approaches can be developed.

## Acknowledgments

We thank Dr. Ian Gilmour, Environmental Protection Agency, for the DEP. We thank Michael Muehlbauer from the Duke Molecular Physiology Institute Metabolomics Research Group for aiding in running the U-Plex MSD. We thank Duke Genomic Analysis and Bioinformatics Shared Resource for aid in sequencing and data analysis. We thank the Duke Light Microscopy Core Facility for access to microscopes and software for image analysis. We thank Michael D. Gunn for the BAC transgenic mice. Biorender was used to generate elements of main figures 1,2,3 and 5 and supplemental figures 2 and 17.

## Author contributions

C.L.B., S.D.M., K.D., C.E. and S.D.B. conceived and planned experiments. C.L.B., O.E., S.D.M., C.S., K.E.M., C.B., D.H., K.A.B., N.N., Y.C.J., T.N. carried out the experiments and data collection. C.L.B., O.E., S.D.M., C.S., C.B., D.H., K.A.B., N.N., A.T., D.C., N.M.G. analyzed the data. C.L.B., K.D., C.E., and S.D.B. wrote the manuscript, with feedback from S.D.M. See supplementary text for detailed author contributions.

## Data and materials availability

All next-generation sequencing data can be viewed at NCBI GEO. Code used for u-net image restoration and cell counts is deposited at https://github.com/ErogluLab/CellCounts. Code for puncta analysis is deposited at: https://github.com/physion/puncta-analyzer.

## Supplementary Materials

Materials and Methods

Supplemental Text

Figures S1-S18

References (92-103)

