## Supplementary Material for "Prenatal Environmental Stressors Impair Postnatal Microglia Function and Adult Behavior in Males"

**This PDF file includes:**

Materials and Methods
Supplementary Text
Figs. S1 to S19
References 92-103

#### Materials and Methods

##### Animals

All experiments were performed in accordance with the guidelines of the Division of Laboratory Animal Resources from Duke University School of Medicine and Institutional Animal Care and Use Committee. For experiments in wildtype animals, we obtained adult male and female C57BL/6J mice from Jackson Laboratories (Bar Harbor, ME) and maintained an internal colony of breeding animals for all experiments. For microglia ablation studies, *Cx3cr1-CreBT* (MW126GSat) mice were generated and provided by Michael D. Gunn and backcrossed over 6 generations to C57BL/6J (Bar Harbor, ME) and crossed to Rosa26-DTR (Stock#: 007900, C57BL/6-Gt(ROSA) 26Sortm1(HBEGF)Awai/J) mice which were purchased from Jackson Laboratories (Bar Harbor, ME).

##### Prenatal stressors

*Diesel exhaust particle exposure.* Performed as previously described (*30),* briefly, adult females were time-mated and checked twice daily for the presence of a vaginal plug, considered to be E0. Females were paired in individually ventilated cages with specialized bedding (AlphaDri; Shepherd Specialty Papers, Milford, NJ) and *ad libitum* access to food (PicoLab Mouse Diet 5058; Lab-Diet, Philadelphia, PA) and filtered water. Females were treated with diesel exhaust particles (DEP) delivered via oropharyngeal aspiration. Beginning on E2, females were lightly anesthetized with 2% isoflurane and treated with either 50 µg of DEP suspended in 50 µL of PBS, 0.05% Tween-20, or vehicle alone (CON). Females received a total of six doses every 3 days from E2-E17.

*Maternal stress/ nest restriction.* To induce maternal stress, we utilized our adaptation of a previously described nest restriction model applied to the postnatal period (*30, 31*). Beginning at 5pm on E13, control females receiving vehicle treatment were singly housed in a clean cage with full size nestlet (CON), and females exposed to DEP were housed in clean cages with a thin layer of bedding under an elevated fine-gauge aluminum mesh platform (mesh dimensions, 0.4 cm × 0.9 cm; McNichols Co., Tampa, FL) and provided with two-thirds of one square of felt-like nesting material (~1.9 g; MS group). This design results in two groups of dams: control dams (CON) and dams exposed to combined environmental stressors (DEP+MS). On E18.5 prior to the birth of pups, all dams were placed into a clean cage with a full size nestlet and were treated identically. All pups were born into standard caging environment and remained with the mother until tissue collection time point or weaning age ~P24, at which time mice were group housed with same-sex littermates at a maximum 5 animals per cage.

##### Microglial ablation

*Cx3cr1-CreBT* $^{tg/0}$ females were mated to iDTR $^{tg/tg}$ males. After birth, all offspring received 80 ng i.p. injection of diphtheria toxin at P5 and P7, effectively eliminating >95% of microglia in the ACC. Pups were raised into adulthood and USVs were collected from males ~P60.

##### Maternal Immune Activation

To determine the immune activation of pregnant dams, on E17.5, 2-3 hours post treatment, a 5.5 mm lancet was used to pierce the submandibular vein and cheek blood was collected into a sterile Eppendorf tube. To separate serum from red blood cells, blood was centrifuged twice at 16,000 ×g at 4°C for 10 mins. Separated serum was collected into a clean Eppendorf tube and stored at -80°C until analysis.

*Maternal Cytokine Analysis.* For serum analysis, a multiplex electrochemiluminescence immunoassay kit (U-Plex Proinflammatory Panel, Mouse) was purchased from Meso Scale Discovery (Rockvillle, MD) and used according to manufacturer instructions to measure serum cytokine concentrations (pg/mL) of IFN-γ, IL-1β, IL-4, IL-5, IL-6, IL-10, IL-17A, IL-12/IL-23p40 MCP-1, and TNF-α. To prevent antibody cross-reactivity IL-17A and IL-12/IL-23p40 were coated in 1 plate, and the remaining antibodies were coated onto a separate plate. Analysis of blinded samples was conducted by Duke Molecular Physiology Institute Metabolomics laboratory. Samples were run in duplicates, plates were read with a Sector Imager 2400 (Meso Scale Discovery), and data were analyzed using the Discovery Workbench 4.0 software (Meso Scale Discovery). Any values below the LLOD were assigned 0 pg/mL, all samples were within the detection range for IFN-γ, IL-5, IL-6, IL-10, IL-17A, IL-12/IL-23p40, MCP-1 and TNF-α. For IL-1β and IL-4, more than half of the samples fell below the LLOD and were excluded from further analyses.

*Maternal CORT measurement*. Corticosterone serum levels were measured using a commercially available ELISA kit (K014-H; Arbor Assays, Ann Arbor, MI). To determine whether method of instillation induced additional stress, control serum was rapidly collected from a non-pregnant female and three WT pregnant females without prenatal treatment. The optical density measurements (Bio-Tek Instruments) from the microplate reader were uploaded to https://www.myassays.com/arbor-assays-corticosterone-enzyme-immunoassay-kit.assay to calculate corticosterone concentration for each sample.

**Behavioral procedures**

We assessed behavioral outcomes as a result of prenatal stressors in a cohort of neonatal (P7-P9) male and female offspring (*n* = 14-17 animals/sex from 4 litters per treatment group). A separate cohort of male offspring were utilized to assess outcomes in adulthood (n=16-20 animals/group from 4 litters per treatment group). A separate cohort of mice undisturbed during development, were generated to assess neurophysiological ends points with concurrent behavior (n=13-27 animals/sex per treatment group).

**Neonatal Ultrasonic vocalizations**

To determine ultrasonic vocalization (USV) developmental timeline in C57BL/6J animals, USVs were collected from P4-P9. For experimental animals, USVs were collected on postnatal days 7-9. Pups were briefly separated from dams and placed into a sound attenuating chamber for 3 minutes, USVs were recorded using an externally polarized condenser microphone with a frequency range of 10-200 kHz attached to the Avisoft-Ultrasound Gate recording software (Avisoft Bioacoustics, Berlin, Germany). Pup weight and toe clip identification was performed immediately after USV collection. WAV files for each pup were converted to spectrograms and analyzed with automated whistle tracking parameters by the Avisoft SASLab Pro software (Avisoft Bioacoutics) and manually validated for accuracy. For call complexity analysis, WAV files were analyzed using Mouse Ultrasonic Profile ExTraction Tool (MUPET) in MATLAB, which is an unsupervised machine-learning-based algorithm that analyzes vocalization parameters, classifies syllables into distinct repertoires and compares vocalization pattern between test groups (*40*). Repertoire units were sorted by length using a Python script.

**Adult Ultrasonic vocalizations**

Prior to behavioral testing in adulthood, all animals were handled five times. Because males are the primary source of USVs in male-female encounters, adult USVs were only collected from male offspring using an estrous-induced courtship paradigm (*78*). In this paradigm, male mice, after gaining sexual experience, are exposed to sexually receptive WT stimulus animals for 5 minutes for 3 days. To identify sexually receptive females, stimulus animals were vaginally swabbed, and cell morphology was assessed to identify females in estrous or proestrous (*92*). Females identified to be in estrous or proestrus were utilized as stimulus animals. USVs were recorded for 5 minutes using an externally polarized condenser microphone with a frequency range of 10-200 kHz attached to the Avisoft-Ultrasound Gate recording software (Avisoft Bioacoustics, Berlin, Germany). WAV files were analyzed using MUPET. To filter noise, calls below 35 kHz were excluded, a noise-reduction value of 8.8 was utilized with a minimum syllable duration of 2.0 msec.

**Electrode implantation surgery**

Mice were anesthetized with 1.5% isoflurane, placed in a stereotaxic device, and metal ground screws were secured above the cerebellum and anterior cranium. The recording bundles designed to target basolateral and central amygdala (AMY), medial dorsal thalamus (MD), nucleus accumbens core and shell (NAc), VTA, medial prefrontal cortex (mPFC), and VHip were centered based on stereotaxic coordinates measured from bregma (Amy: -1.4mm AP, 2.9 mm ML, -3.85 mm DV from the dura; MD: -1.58mm AP, 0.3 mm ML, -2.88 mm DV from the dura; VTA: -3.5mm AP, ±0.25 mm ML, -4.25 mm DV from the dura; VHip: -3.3mm AP, 3.0mm ML, -3.75mm DV from the dura; mPFC: 1.62mm AP, ±0.25mm ML, 2.25mm DV from the dura; NAc: 1.3mm AP, 2.25mm ML, -4.1 mm DV from the dura, implanted at an angle of 22.1°). We targeted prelimbic cortex, infralimbic cortex using the PFC bundle by building a 0.5mm and 1.1mm DV stagger into our electrode bundle microwires, and animals were implanted bilaterally in mPFC and VTA; all other bundles were implanted on the left side. The NAc bundle included a 0.6mm DV stagger such that wires were distributed across NAc core and shell. We targeted BLA and CeA by building a 0.5mm ML stagger and 0.3mm DV stagger into our AMY electrode bundle. In order to mitigate pain and inflammation related to the procedure, all animals received carprofen (5 mg/kg, s.c.) injections once prior to surgery and then once every 24 hours for three days following electrode implantation.

**Histological Confirmation**

Histological analysis of implantation sites was performed at the conclusion of experiments to confirm recording sites used for neurophysiological analysis. Animals were perfused with 4% paraformaldehyde and brains were harvested and stored for 24 hours in PFA. Brains were cryoprotected with sucrose and frozen in OCT compound and stored at -80C. Brains were sliced at 35µm and stained using NeuroTrace fluorescent Nissl Stain (N21480, ThermoFisher Scientific, Waltham, MA). Floating sections were washed 3 times in PBST (0.1%). Sections were incubated in PBS with Nissl antibody (1:300) for 10 mins at room temperature and washed once in PBST (0.1%) and twice in PBS with azide (0.01% $NaN_3$), after which the entire brain was mounted. Images were obtained using a Nikon Eclipse fluorescence microscope at 4× and 10× magnifications. Only animals in which all eight implantation sites were confirmed were included in the analysis. Multiple animals were removed due to tissue destruction during histological analysis in which implantation could not be confirmed.

**Neurophysiological data acquisition**

Mice were connected to a headstage (Blackrock Microsystems, UT, USA) without anesthesia, and placed in each behavioral arena. Neuronal activity was sampled at 30kHz using the Cerebus acquisition system (Blackrock Microsystems Inc., UT). Local field potentials (LFPs) were bandpass filtered at 0.5–250Hz and stored at 1000Hz. Neurophysiological recordings were referenced to a ground wire connected to both ground screws.

**Two-chamber social Interaction test**

Social preference was measured using a two-chamber assay in which animals explored a novel object or a novel mouse. The apparatus was a rectangular arena (61cm × 42.5cm × 22cm) constructed from clear plexiglass with a clear plexiglass wall dividing the arena into two equal chambers with an opening in the middle allowing free access between both chambers. The floor of the arena was constructed using a one-way mirror that allowed for video recording from beneath in order to avoid obstruction from electrophysiological recording equipment. Plastic, circular holding cages (8.3cm diameter and 12cm tall) were centered in each of the two chambers and were used to house either a novel object or sex- and age-matched C3H target mouse. The arena was evenly lit with indirect white light (~125 lux). Test mice were handled and habituated to the social preference chambers and empty holding cages for a least three days prior to testing. Subsequently, mice underwent ten separate social preference test sessions, with at least one day off in between sessions, in which the test mice were allowed to freely explore the arena for ten minutes; the holding cages contained either a novel object or novel C3H target mouse. The side of the chamber holding the object/mouse was determined pseudorandomly, such that the object/mouse would not be placed in the same chamber on more than two consecutive sessions in order to prevent side biases and to distinguish target-specific effects from location-specific effects. Plastic toys and glass objects were used as novel objects with the object being between 3-5cm in all directions. Video data was tracked using Bonsai Visual Reactive Programming software and the time spent in the proximity (4.98cm) of either holding cage was used to determine social preference scores.

The social preference for each session was defined as:

$$\frac{\overline{InteractionTime_S} - \overline{InteractionTime_O}}{\overline{InteractionTime_S} + \overline{InteractionTime_O}}$$

where $Interactions_S$ is the total time spent proximal to the other mouse, and $InteractionTime_O$ is the total time spent proximal to the object.

**LFP preprocessing to remove signal artifact**

Rather than manually screening data, we used an automated heuristic strategy to remove recording segments with non-physiological signals. First, we estimated the envelope of the signal in each channel using the magnitude of the Hilbert transform. For any 1-second window where the envelope exceeds above a pre-selected low threshold, the entire segment is removed if the envelope exceeds a second, high threshold at any point within that window. The two thresholds were determined independently for each brain region. The high threshold was selected to be 5 times the median absolute deviation of the envelope value for that region. Five median absolute deviations were chosen as the high threshold because it is roughly equivalent to 3 standard deviations from the mean for normally distributed data, but is robust to outliers in the data. The low threshold was empirically chosen to be 3.33% of the high threshold. If more than half the window was removed for a channel, we removed the rest of that window for that channel as well. In addition, any windows where the standard deviation of the channel is less than 0.01 were also removed. Using this approach, 13±3.5% of the data per mouse were excluded from this analysis. This conservative strategy optimized the potential of our learning model to discover a network that was uniquely related to appetitive social emotional brain states.

**Feature Estimation**

The LFPs were averaged across electrodes within each brain-region to yield a more robust estimate of the LFP for each region. Each LFP recording was divided into 1 second windows with a univariate time series associated with each region. Feature extraction was performed with MATLAB (The MathWorks, Inc., Natick, MA). The three features of interest were frequency-based power within each region, frequency-based coherence between each pair of regions, and frequency-based Granger causality between each region.

For estimating power, we used the pwelch function in MATLAB, which averages multiple periodograms estimated using different segments of the window to obtain a denoised power spectrum. A sliding Fourier transform with a Hamming window was applied to the average LFPs (default pwelch settings) and the power was estimated at 1Hz intervals. Estimating the frequency-based coherence was done using magnitude-squared coherence, defined as

$$C_{AB}(f) = \frac{|Psd_{AB}(f)|^2}{Psd_{AA}(f)Psd_{BB}(f)},$$

which normalizes the cross-spectral estimates by the power spectra in each region, yielding a value between 0 and 1. This was done in MATLAB using the function mscohere with default settings, also at a resolution of 1 Hz.

The Granger causality is a measure of causal information flow between two signals (*53*). While the original definition did not decompose this flow by frequency, work by (Barnett and Seth, 2014) developed the theory and toolbox to do this, known as the *Multivariate Granger Causality* (MVGC) MATLAB toolbox (*93*). We used standard procedure as defined in the method, the non-stationary data went through a highpass Butterworth filter with a stopband at 1Hz and a passband starting at 4Hz. Granger causality values for each window were calculated using a 20-order AR model via the *GCCA_tsdata_to_smvgc* function of the MVGC toolbox. Once again, these causality values were estimated at the same frequency intervals as the power and coherence.

The Granger features themselves are not additive, a major drawback with most factor models. Rather than using the features directly, we used the exponential of all causality values, which can be interpreted as a ratio of total power to the unexplained power. That is,

$$\exp\left(\mathrm{f}_{\mathrm{Y}\to X}(\lambda)\right) = \frac{|\mathrm{S}_{\mathrm{XX}}(\lambda)|}{\left|\mathrm{S}_{\mathrm{XX}}(\lambda) - \mathrm{H}_{\mathrm{XY}}(\lambda)\Sigma_{\mathrm{Y|X}}\mathrm{H}_{\mathrm{XY}}(\lambda)^{*}\right|}$$

where $f_{Y\to X}(\lambda)$ represents Granger causality at frequency $\lambda$ from region *Y* to region *X*, $S_{XX}(\lambda)$ represent the spectral power in region *X* at frequency $\lambda$, and $H_{XY}(\lambda)\Sigma_{Y|X}H_{XY}(\lambda)^{*}$ represents the component of that power that is predicted by region *Y*. These values can be occasionally very large due to estimation error, and was capped at 10 to prevent undue influence from single observations.

**Discriminative cross spectral factor analysis non-negative matrix factorization (dCSFA-NMF)**
We used a non-negative matrix factorization to synthesize these estimated features into a network-based model of neural dynamics. This is termed Supervised Cross-Spectral Factor Analysis – Nonnegative Matrix Factorization (CSFA-NMF). This model is fully described elsewhere (*53*), and the code to implement these models is publicly available at https://github.com/carlson-lab/encodedSupervision. To provide a succinct description of the methodology, CSFA-NMF assumes each window of data to be an independent stationary observation. Relevant dynamics and behavior occur at the timescale of windows rather than individual LFP measurements. In this work we chose a 1 second window as a compromise between fast-changing dynamics in behavior and the extra stability in feature estimation provided by longer windows. Prior work has shown that shorter windows decrease predictive accuracy, and 5 second windows would not be fast enough to capture the rapidly changing behavioral dynamics needed for these experiments.

Each window of data consists of the estimated power, coherence and exponential granger features totaling P distinct observations per window. These observations were vectorized. We use $n$ to denote a window within the $N$ total windows. We describe the preprocessed data as $X_n \in \mathbb{R}_+^P$ (the $P$-dimensional non-negative domain) and the observed behavioral label as $y_n \in \{0,1\}$, where the binary indicates a social or non-social behavioral label. The objective function learned by this model is

$$\min_{W,d,\phi} \sum_{n=1}^{N} \|x_n - Wf(x_n;\phi)\|_2^2 + \lambda\|y_n - d^T f(x_n;\phi)\|_2^2,$$

where K is the number of different electomes. Each electome is described by a column in $W \in \mathbb{R}_+^{P\times K}$ (e.g., $W = [w_1, \dots, w_K]$) that describes the multi-region spectral power and coherence relationships. The electome factor scores are given by the multi-output function $f(x_n;\phi)\colon \mathbb{R}_+^p \to \mathbb{R}_+^K$, and the relationship between the electome factor scores and the behavioral labels is given by $d \in \mathbb{R}^p$. $\lambda$ balances the relative importance of prediction relative to reconstruction. $d$ was defined to have a single non-zero element in order to limit the predictive capacity to a single latent network. This is a formulation of an NMF model that performs approximate inference using supervised autoencoders and requires the user to choose a parametrization for $f(x_n; \phi)$. In our method, this is simply set to an affine function following by a non-linearity, $f(x_n; \phi) = softplus(Ax + b)$, where the parameters of the function are $\phi = \{A, b\}$ and the softplus means an element-wise operation of the operation $\mathrm{softplus}(a) = \log(1 + \exp(a))$, which maps a real number to the non-negative space. While other rectifying functions are possible (such as the popular Rectified Linear Unit (ReLU)), we chose the softplus to prevent vanishing gradients in the parameter estimation.

This model is able to be learned through stochastic gradient descent and was implemented in TensorFlow 1.09 using the ADAM algorithm for learning. In addition to the benefits of increasing predictive ability, replacing explicit network score estimates with a predictive function allows for quicker inferences with stochastic rather than batch training. Furthermore, once a predictive function is learned we can calculate the electome scores on new data simply by calling the function $f(x_n; \phi)$. This contrasts with other methods which typically require a potentially difficult optimization problem to estimate each new electome score. This allows for future application requiring real-time estimation.

**Hyper-parameter Selection**
This analysis requires us to choose several parameters, notably the number of electomes *K*, the supervision strength $\lambda$, the relative importance of the features, and the parameterization of the mapping function $f(x_n; \phi)$. For the mapping function we chose an affine transform with a softplus activation to avoid overfitting and to prevent vanishing gradients respectively. Our analysis has two goals, to predict behavior in new animals well and to describe the brain dynamics accurately. These two goals are measured by the reconstruction error of the features and the by the mean Area Under the Curve (mean AUC) on validation mice respectively. Choosing the supervision strength was chosen to be a value found to work well in previous analyses. The number of networks *K* was chosen using an elbow analysis using an unsupervised NMF model, where we chose *K* to be the number of networks where minimal gains in reconstructive loss were observed.  This model's parameters were learned elsewhere (*53*).  The electome scores on the mice in this paper are estimated by putting the extracted features through the previously trained function $f(x_n; \phi)$. Thus, since the mice in this paper were *not* used for hyperparameter selection or training, and thus represent a true estimate of the accuracy and reconstructive ability of the model when applied to the this novel population.

**RNA-Seq analysis of transcriptome**
*Tissue and sample preparation*. Tissue samples were harvested from a cohort of behaviorally naïve P8 pups. Animals were anesthetized with avertin and perfused with saline (n=4

animals/group/sex). The brain was immediately extracted, and the prefrontal cortex was dissected before being flash frozen in liquid nitrogen and stored at -80°C until RNA-extraction.

*RNA-extraction*. Frozen samples were homogenized in 1000 µl TRIzol Reagent (15596026, Thermo Fisher Scientific, Waltham, MA) and vortexed at 2000 rpm for 5 min. 200 µl of chloroform (Sigma-Aldrich, C2432, St. Louis, MO) was added to each tube and vortexed for additional 2 min, samples were allowed to phase separate before being centrifuged at 11,900 rpm for 15 min at 4°C, after which the top clear aqueous phase was separated into a fresh tube. 500 µl of Isopropanol (Thermo Fisher Scientific, NY) was added and samples were vortex at 2000 rpm for 1 min and incubated at room temperature for an additional 10 minutes and then centrifuged for 10 mins. Supernatant was discarded and RNA pellet was washed two times with 1 ml of ice cold 75% ethanol, air dried and resuspended in 40 µl of RNase-free water.

*Library prep and sequencing*. All RNA samples were coded numerically. Sequencing was performed blind to sample identity by Sequencing and Genomic Technologies Shared Resource Duke Center for Genomic and Computational Biology. Extracted total RNA quality and concentration was assessed on Fragment Analyzer (Agilent Technologies) and Qubit 2.0 (ThermoFisher Scientific), respectively. RNA-seq libraries were prepared using the commercially available KAPA Stranded mRNA-Seq Kit (Roche). In brief, mRNA transcripts are first captured using magnetic oligo-dT beads, fragmented using heat and magnesium, and reverse transcribed using random priming. During the 2$^{nd}$ strand synthesis, the cDNA:RNA hybrid is converted into to double-stranded cDNA (dscDNA) and dUTP incorporated into the 2$^{nd}$ cDNA strand, effectively marking the second strand. Illumina sequencing adapters are then ligated to the dscDNA fragments and amplified to produce the final RNA-seq library. The strand marked with dUTP is not amplified, allowing strand-specificity sequencing. Libraries were indexed using a dual indexing approach allowing for all the libraries to be pooled and sequenced on the same sequencing run. Before pooling and sequencing, fragment length distribution for each library was first assessed on a Fragment Analyzer (Agilent Technologies). Libraries were also quantified using Qubit. Molarity of each library was calculated based on qubit concentration and average library size. All libraries were then pooled in equimolar ratio and sequenced. Sequencing was done on an Illumina NovaSeq 6000 sequencer. The pooled libraries were sequenced on a S-Prime flow cell at 50bp paired-end. Once generated, sequence data was demultiplexed and Fastq files generated using bcl2fastq v2.20.0.422 file converter from Illumina.

*Transcriptome data analysis methods*. RNA-seq data was processed by the Genomic Analysis and Bioinformatics Shared Resource, Duke Center for Genomics and Computational Biology using the TrimGalore toolkit (http://www.bioinformatics.babraham.ac.uk/projects/trim_galore) which employs Cutadapt *(94*) to trim low-quality bases and Illumina sequencing adapters from the 3' end of the reads. Only reads that were 20nt or longer after trimming were kept for further analysis. Reads were mapped to the GRCm38v73 version of the mouse genome and transcriptome (*95*) using the STAR RNA-seq alignment tool (*96*). Reads were kept for subsequent analysis if they mapped to a single genomic location. Gene counts were compiled using the HTSeq tool (http://www-huber.embl.de/users/anders/HTSeq/). Only genes that had at least 10 reads in any given library were used in subsequent analysis. Normalization and differential expression were carried out using the DESeq2 (*97*) Bioconductor (*98*) package with the R statistical programming environment (www.r-project.org). We controlled for plate in each model that we ran. The false discovery rate was calculated to control for multiple hypothesis testing. Gene set enrichment analysis (*99*) was performed to identify pathways associated with altered gene expression for each of the comparisons, PANTHER (http://www.pantherdb.org/) was used to perform a statistical overrepresentation test (*100*).

**Immunohistochemistry**

Mice used for IHC were anesthetized with 200 mg/kg tribromoethanol (avertin) and perfused with Tris-Buffered Saline (TBS, 25 mM Tris-base, 135 mM NaCl, 3 mM KCl, pH 7.6) supplemented with 7.5 mM heparin, followed by 4% PFA in TBS. Brains were extracted and post-fixed in 4% PFA in TBS overnight at 4C. After fixation, brains were washed 3 times with TBS and transferred to a 30% sucrose/TBS solution for cryoprotection. Brains were frozen and embedded into a solution containing 2 parts 30% sucrose and 1 part OCT (Tissue Tek, Sakura, Torrance, CA), and stored at -80°C. For synaptic staining brains were sections at 20 μm thickness, for cell counting and microglia reconstructions brains were sectioned at 40 μm, tissue sections were stored floating in a 1:1 mixture of TBS/glycerol at -20°C.

*Synaptic staining*. Free-floating sections were washed 3 times for 10 minutes with TBS with 0.2% Triton X-100 (Roche, Indianapolis, IN) and blocked in 5% Normal Goat Serum (NGS; Jackson ImmunoResearch, West Grove, PA) with 0.2% Triton X-100 in TBS for 1 hour at room temperature. Primary antibodies (see table below) were diluted in 5% NGS in TBS with 0.2% Triton X-100. Sections were incubated overnight at 4°C with primary antibodies and washed three times for 10 minutes with TBS the following morning. Secondary Alexa-fluorophore conjugated antibodies (Invitrogen, Carlsbad, CA) were diluted (1:200) in 5% NGS in TBS with 0.2% Triton X-100, and sections were incubated with secondary antibodies for 2 hours at room temperature, protected from light. After incubation, sections were washed three times for 15 minutes in TBS and mounted with VECTASHIELD with DAPI (Vector Laboratories, Burlingame, CA). Images were acquired on an Olympus FluoView 3000 confocal laser-scanning microscopes.
*Acquisition and analysis of synaptic staining*. Staining, image acquisition, and analysis were performed as in Ippolito and Eroglu (2010) (*101*) with adjustments. Synaptic staining was performed in two male/female littermate pairs at P6, P8, P10, P13 and P15 in WT C57BL6/J offspring to determine the normal developmental pattern. Synaptic staining was performed at P8, P15 and P100 in male and female offspring for CON and DEP+MS conditions. Image acquisition was performed in layer 1 (L1) of the ACC from P8, P15 and adult CON and DEP+MS animals. We chose to conduct our analyses in L1, because this layer contains sparse neuronal cell bodies, and receives dense axonal inputs from both thalamic and neighboring regions. 5.1 mm-thick confocal images (optical section depth 0.33 μm, 15 sections/scan) were acquired at 60× magnification plus 1.4× optical zoom using the Olympus FluoView 3000 confocal microscope or Zeiss 880.

*Synapse quantification.* Maximum projections of 3 consecutive optical sections were generated using ImageJ. The Puncta Analyzer Plugin (available at: https://github.com/physion/puncta-analyzer) for ImageJ was used to count the number of colocalized synaptic puncta. The individual analyzing the images was always blinded to the experimental conditions. At least 5 maximum projections per brain, from 3 brain sections per animal, were analyzed using a nested t-test.

*Cell staining*. Free-floating sections were washed 3 times for 10 minutes with TBS with 0.5% Triton X-100 (Roche, Indianapolis, IN) and blocked in 5% Normal Goat Serum (NGS; Jackson ImmunoResearch, West Grove, PA) with 0.5% Triton X-100 in TBS for 1 hour at room temperature. Primary antibodies (see table below) were diluted in 5% NGS in TBS with 0.5% Triton X-100. Sections were incubated overnight at 4°C with primary antibodies and washed three times for 10 minutes with TBS the following day. Secondary Alexa-fluorophore conjugated antibodies (Invitrogen, Carlsbad, CA) were diluted (1:500) in 5% NGS in TBS with 0.5% Triton X-100, and sections were incubated with secondary antibodies for 2 hours at room temperature, protected from light. During the last five minutes of secondary incubation DAPI was added to achieve a dilution of 1:40,000 (ThermoFisher D1306). After incubation, sections were washed three times for 15 minutes in TBS and mounted with an in-house mounting media (20mM Tris pH8.0, 90% Glycerol, 0.5% N-propyl gallate). Images were acquired on an Olympus Fluoview 3000 confocal laser-scanning microscope.

*Microglia staining for cell counts.* Free-floating sections were washed 3 times for 10 minutes with TBS with 0.5% Triton X-100 (Roche, Indianapolis, IN) and blocked in either 5% Normal Donkey Serum (NDS; Jackson ImmunoResearch, West Grove, PA) with 0.5% Triton X-100 in TBS for 1 hour at room temperature. Primary antibodies (see table below) were diluted in 5% NDS in TBS with 0.5% Triton X-100. Sections were incubated overnight at room temperature with primary antibodies and washed three times for 10 minutes with TBS the following day. Secondary Alexa-fluorophore conjugated antibodies (Invitrogen, Carlsbad, CA) were diluted (1:500) in 5% NDS in TBS with 0.5% Triton X-100, and sections were incubated with secondary antibodies for 2 hours at room temperature, protected from light. During the last five minutes of secondary incubation DAPI was added to achieve a dilution of 1:40,000 (ThermoFisher D1306). After incubation, sections were washed three times for 15 minutes in TBS and mounted with an in-house mounting media (20mM Tris pH8.0, 90% Glycerol, 0.5% N-propyl gallate). Images were acquired on an Olympus Fluoview 3000 confocal laser-scanning microscopes.

*Acquisition and analysis for cell counts of neurons, astrocytes and oligodendrocytes.* Cell counts for NeuN, Olig2 and Sox9 were performed in male and female CON and DEP+MS offspring at P8. Coronal brain sections (40 μm) containing the ACC triple labeled with NeuN, Olig2 and Sox9. Confocal *z*-stacks of the ACC were acquired using the 30× objective on an Olympus FluoView 3000 microscope. Tile scans of 10 μm *z*-stacks were acquired for the entire ACC using a 1.0 μm step size. To expedite imaging, a neural network was trained to denoise images from the resonant scanner (*102*). Briefly, high-resolution images were acquired using the galvanometer scanner. Gaussian noise was added to reduce the signal-to-noise ratio to levels expected from the resonant scanner. These degraded images were used to train a neural network. This neural network model was applied to images acquired using a resonant scanner with the same objective and confocal. Restored images were stitched using the grid/collection stitching feature in Fiji (1.52p). Images were max-projected, and an ROI of the ACC was applied, the ROI was pseudo-layered into bins of 160-micron lengths from the midline to layer 6 of the ACC. Incomplete layers were not counted. Automated cell counting was performed using the U-Net deep neural network (*103)*. Separate models were trained for each individual marker (NeuN, Olig2 and Sox9) and were manually verified for accuracy.

*Acquisition and analysis for cell counts of microglia.* Microglia cell counts of Iba1 and/or P2ry12 positive cells were performed in male and female CON and DEP+MS offspring at P8, P15 and P25. Coronal brain sections containing the ACC triple labeled with P2ry12, Iba1 and CD68. Confocal z-stacks of the ACC were acquired using the 30x silicone objective on an Olympus FluoView 3000 microscope. Tile scans of 10-micron z-stacks were acquired for the entire ACC using a 1.0-micron step size. Images were stitched using Olympus software, Images were max projected, and an ROI of the ACC was applied, the ROI was pseudo-layered into bins of 160-micron lengths from the midline to layer 6 of the ACC. Incomplete layers were not counted. Cell counts were manually performed in Fiji using cell counter feature. Cells were counted when positive for DAPI and a microglial cell marker (Iba1 and/or P2ry12). Total number of microglia cells includes cells positive for DAPI and 1 or 2 microglial markers. Heterogenous microglia cells are considered cells expressing only high levels of one microglia marker (Iba1$^{hi}$P2ry12$^{lo}$, Iba1$^{lo}$P2ry12$^{hi}$). Percent heterogeneity was quantified as number of singly high labeled microglia over total number of all microglia cells.

*Acquisition and analysis for Cd68 quantification in heterogeneous microglia.* Cd68 content was quantified in heterogenous microglia at P8 in CON and DEP+MS male offspring. 40 µm coronal brain sections containing the ACC were triple labeled with Iba1, P2ry12 and CD68. Confocal z-stacks of the ACC were acquired using the 30× objective on an Olympus FluoView 3000 microscope. Using the resonant scanner tile scans of 20 µm *z*-stacks were acquired for the entire ACC using a 0.35 µm step size. Images were stitched using the grid/collection stitching feature of Fiji (1.52p). To enable the expedited acquisition of large tile scanned images, a deep neural network (U-Net) was utilized to denoise images (*98*). A lab specific pipeline was generated using Python and is available from (repository). Imaris software 9.5.1 was used to create surface renderings of individual microglia cells labeled with either Iba1 or P2y12, incomplete or poorly labeled cells were excluded from analyses, Cd68 content within microglia surface makers was quantified. After surface renderings, cells were identified as single or double positive and Cd68 content was normalized to cell volume.

*Microglia engulfment staining.* Free-floating sections were washed 3 times for 10 minutes with TBS with 0.5% Triton X-100 (Roche, Indianapolis, IN) and blocked in either 5% Normal Goat Serum (NGS; Jackson ImmunoResearch, West Grove, PA, NDS; Jackson ImmunoResearch, West Grove, PA) with 0.5% Triton X-100 in TBS for 1 hour at room temperature. Primary antibodies (see table below) were diluted in 5% NGS TBS with 0.5% Triton X-100. Sections were incubated overnight at room temperature with primary antibodies and washed three times for 10 minutes with TBS the following morning. Secondary Alexa-fluorophore conjugated antibodies (Invitrogen, Carlsbad, CA) were diluted (1:500) in 5% NGS in TBS with 0.5% Triton X-100, and sections were incubated with secondary antibodies for 2 hours at room temperature, protected from light. During the last five minutes of secondary incubation DAPI was added to achieve a dilution of

1:40,000 (ThermoFisher D1306). After incubation, sections were washed three times for 15 minutes in TBS and mounted with an in-house mounting media (20mM Tris pH8.0, 90% Glycerol, 0.5% N-propyl gallate). Images were acquired on an Olympus Fluoview 3000 confocal laser-scanning microscopes.

*Acquisition and analysis of synaptic engulfment.* Synaptic engulfment was performed in two pairs of WT C57BL6/J male offspring at P6, P8, P10, P13 and P15 to determine the normal developmental pattern of synapse elimination in the ACC. Synapse engulfment was quantified at P8 and P10 in CON and DEP+MS male offspring. For WT characterization and P10 analysis, 40 um coronal brain sections containing the ACC were stained for Iba1 and P2ry12 on the same fluorophore, Cd68 and VGlut2. For P8 engulfment analyses coronal brain sections containing the ACC were stained for Iba1, P2ry12 and VGlut2 all on separate fluorophores, due to the limit of fluorophores and inadequate stability of the 450 fluorophores, CD68 was excluded from these analyses. Confocal z-stacks of the ACC were acquired using the 60× oil objective on an Olympus FluoView 3000 microscope (experimental groups) or Zeiss 880 (WT characterization), with 2.0× zoom. An entire microglia cell was capture with 0.35 um step size. Huygens Professional 19.10.0p3 64b was used to deconvolve images. Imaris 9.5.1 was used to create surface renderings of individual microglia cells, Cd68 within microglia surface and VGlut2 within microglia surface. Volume of phagocytes and engulfed synapses is normalized to cell volume.

IHC antibody table

| **Antibody name** | **Vendor** | **Dilution for IHC** |
|---|---|---|
| NeuroTrace fluorescent Nissl Stain | Thermofisher N21480 | 1:300 |
| Guinea pig anti-VGlut1 | Millipore Sigma Aldrich AB5905 | 1:2000 |
| Guinea pig anti-VGlut2 | Synaptic Systems 135404 | 1:2000 |
| Rabbit anti-PSD95 | Life Technologies 51-6900 | 1:350 |
| Rat anti-CD68 | BioLegend 137002 | 1:500 |
| Chicken anti-IBA1 | Synaptic Systems 234006 | 1:1000 |
| Rabbit anti-P2y12 | Anaspec AS-55043A | 1:2000 |
| Goat anti-Iba1 | Novus Biologicals NB 100-1028 | 1:500 |
| Rabbit anti-Sox9 | Millipore Sigma Aldrich AB5535 | 1:2000 |
| Mouse anti-Olig2 | Millipore Sigma Aldrich MABN50 | 1:250 |
| Mouse anti-NeuN | Millipore Sigma Aldrich MAB377 | 1:500 |
| Guinea pig anti-VGAT | Synaptic Systems 131004 | 1:1000 |
| Rabbit anti-Gephryin | Synaptic Systems 147002 | 1:1000 |
| DAPI | ThermoFisher D1306 | 1:10,000 |

*Statistics.* We analyzed all data using GraphPad Prism version 8.0 (San Diego, CA), MATLAB Version 2017a (Natick, MA) or TIBCO Statistica Software version 13.5.0.17 (Palo Alto, CA). Student's t-test was used to analyze data sets with two groups or Rank-sum test for nonparametric data. One-way ANOVAs were used to analyze data sets with more than two groups. Two-way ANOVAs were used to analyze data sets with two independent variables. Nested analyses were performed for sets of data using biological replicates. Levene's test for homogeneity of variance was used to determine differences in distribution. Spearman's correlation was used to test the relationship between two variables and analysis of covariance was used to test the regression pattern between two groups. All data are represented as mean +/- SEM. See tables below for detailed statistical measures for all analyses.

#### Supplementary Text

Statistics for main and supplemental figures:

Figure 1. Statistics

| | Comparison | Statistical test | *n* | *Sexes included* | *Variable* | Statistic | *p-value* |
|---|---|---|---|---|---|---|---|
| B | Maternal TNF-alpha: CON vs DEP+MS dams | Unpaired *t*-test | 5-6 dams/condition | Females | Condition | t(9)=3.404 | 0.0078 |
| C | Maternal CORT: CON vs DEP+MS dams | Unpaired *t*-test | 5-6 dams/condition | Females | Condition | t(9)=6.211 | 0.0002 |
| F | Number of calls (P8): Condition vs Sex | Two-way ANOVA | 13-19 animals/condition/sex | Males and Females | Condition | F (1, 60) = 15.08 | 0.0003 |
| | | | | | Sex | F (1, 60) = 0.05782 | 0.8108 |
| | | | | | Interaction | F (1, 60) = 0.2213 | 0.6398 |
| | CON Males: DEP+MS Males | Holm-Sidak's post hoc | 14-19 animals/condition | Males | Condition | t(60)= 3.11 | 0.0057 |
| | CON Females: DEP+MS Females | Holm-Sidak's post hoc | 15-16 animals/condition | Females | Condition | t(60)= 2.389 | 0.02 |
| G | Total time calling (P8): Condition vs Sex | Two-way ANOVA | 14-19 animals/condition/sex | Males and Females | Condition | F (1, 60) = 6.401 | 0.014 |
| | | | | | Sex | F (1, 60) = 0.2461 | 0.6216 |
| | | | | | Interaction | F (1, 60) = 0.2694 | 0.6057 |
| | CON Males: DEP+MS Males | Holm-Sidak's post hoc | 14-19 animals/condition | Males | Condition | t(60)= 2.178 | 0.0656 |
| | CON Females: DEP+MS Females | Holm-Sidak's post hoc | 15-16 animals/condition | Females | Condition | t(60)= 1.408 | 0.1643 |
| I | Upregulated/Downregulated | DESeq2 | 4 animals/condition/sex | Males and Females | Condition | no FDR | 0.05 |
| K | Overrepresented GO terms downregulated in males | Fisher's Exact with FDR <0.05 | n=4 animals/condition | Males | Condition | | |
| | Panther GO cellular component | | | | **NES** | **FDR** | **p-value** |
| | Excitatory synapse | | | Males | -13.58 | 1.14E-02 | 4.54E-05 |
| | Axon | | | Males | -4.73 | 3.21E-04 | 3.19E-07 |
| | Postsynapse | | | Males | -3.71 | 1.24E-02 | 5.53E-05 |
| | Presynapse | | | Males | -3.7 | 4.75E-02 | 4.24E-04 |
| | Synapse part | | | Males | -3.63 | 4.75E-04 | 1.18E-06 |
| | Somatodendritic comparement | | | Males | -3.45 | 3.01E-03 | 8.97E-06 |
| | Synapse | | | Males | -3.28 | 4.01E-04 | 7.96E-07 |
| | Neuron projection | | | Males | -3.18 | 4.78E-04 | 7.11E-07 |
| | Neuron part | | | Males | -2.98 | 2.61E-04 | 1.30E-07 |

Figure 2. Statistics

| | Comparison | Statistical test | n | Sexes include d | Variable | Statistic | p-value |
|---|---|---|---|---|---|---|---|
| B. Left | Social preference: CON vs DEP+MS Males | Rank-sum test | 13-14 animals/conditio n | Males | Condition | U=151 | 0.031 |
| B. Righ t | Social preference: CON vs DEP+MS Females | Rank-sum test | 13-14 animals/conditio n | Females | Condition | U=182 | 0.51 |
| D. | Social Network Activity: all animals | Sign-rank test | 32 mice | Males and Females | Stimulus | | 1.210×10 -6 |
| E. | Area Under Curve vs Social preference: all animals | Spearman Correlation | 32 mice | Males and Females | | | 0.005 |
| F. | Correlation (AUC vs Social preference):CON vs DEP+MS Males | Analysis of covariance | 7-11 animals/conditio n | Males | Condition | F(1,14)=6.9 2 | 0.02 |
| | CON Males | Spearman's Correlation post hoc | 11 mice | Males | | RHO = 0.85 | 0.002 |
| | DEP+MS Males | Spearman's Correlation post hoc | 7 mice | Males | | RHO =-0.29 | 0.56 |
| G. | Correlation (AUC vs Social preference) CON vs DEP+MS Females | Analysis of covariance | 5-8 animals/conditio n | Females | Condition | F(1,9)=0.93 | 0.36 |
| | CON Females | Spearman's Correlation post hoc | 5 mice | Females | | RHO=0.60 | |
| | DEP+MS Females | Spearman's Correlation post hoc | 8 mice | Females | | RHO=0.81 | |

#### Figure 3. Statistics

| | Comparison | Statistical test | n | Sexes included | Variable | Statistic | p-value |
|---|---|---|---|---|---|---|---|
| D | Thalamocortical synapses across development: age | One-way ANOVA | 4 animals/age; 3 sections/animal; 5 z-stacks/section | Males and Females | Age | F (4, 48) = 7.153 | 0.0001 |
| | P6:P8 | Holm-Sidak's multiple comparisons test | | sexes combine d | | t(48)= 2.352 | 0.0228 |
| | P6:P10 | Holm-Sidak's multiple comparisons test | | sexes combine d | | t(48)= 2.721 | 0.018 |
| | P6:P13 | Holm-Sidak's multiple comparisons test | | sexes combine d | | t(48)= 4.628 | 0.0001 |
| | P6:P15 | Holm-Sidak's multiple comparisons test | | sexes combine d | | t(48)= 4.513 | 0.0001 |
| F | Microglia phagocytic capacity across development: age | One-way ANOVA | 2 animals/age; 5-10 cells/animal | Males | Age | F (4, 63) = 7.128 | 0.0001 |
| | P8:P6 | Holm-Sidak's multiple comparisons test | 4:4 | Males | | t(63)=0.835 3 | 0.4067 |
| | P8:P10 | Holm-Sidak's multiple comparisons test | 4:4 | Males | | t(63)=3.47 | 0.0028 |
| | P8:P13 | Holm-Sidak's multiple comparisons test | 4:4 | Males | | t(63)=3.189 | 0.0044 |
| | P8:P15 | Holm-Sidak's multiple comparisons test | 4:4 | Males | | t(63)=4.987 | <0.000 1 |
| G | VGlut2 Engulfment across development: age | One-way ANOVA | 2 animals/age; 5-10 cells/animal | Males | Age | F (4, 63) = 4.749 | 0.0021 |
| | P8:P6 | Holm-Sidak's multiple comparisons test | | Males | | t(63)=1.517 | 0.1344 |
| | P8:P10 | Holm-Sidak's multiple comparisons test | | Males | | t(63)=3.315 | 0.0046 |
| | P8:P13 | Holm-Sidak's multiple comparisons test | | Males | | t(63)=2.74 | 0.0159 |
| | P8:P15 | Holm-Sidak's multiple comparisons test | | Males | | t(63)=4.096 | 0.0005 |
| I | VGlut2 Synapse (P8): CON vs DEP+MS Males | Nested t-test | 3 animals/condition; 3 sections/animal; 5 z-stacks/section | Males | Conditio n | t(28)=5.350 | <0.000 1 |
| | VGlut2 Synapse (P15): CON vs DEP+MS Males | Nested t-test | 3 animals/condition; 3 sections/animal; 5 z-stacks/section | Males | Conditio n | t(28)=2.655 | 0.0198 |
| | VGlut2 Synapse (P100): CON vs DEP+MS Males | Nested t-test | 3 animals/condition; 3 sections/animal; 5 z-stacks/section | Males | Conditio n | t(28)=2.695 | 0.0174 |
| K | Microglial Engulfment (P10): CON vs DEP+MS Males | Nested t-test | 3 animals/condition; 10-15 microglia/animal | Males | Conditio n | t(73)=2.285 | 0.0262 |
| L | Microglial Phagocytic activity (P10): CON vs DEP+MS Males | Nested t-test | 3 animals/condition; 10-15 microglia/animal | Males | Conditio n | t(73)=3.357 | 0.0013 |
| M | Microglial Volume (P10): CON vs DEP+MS Males | Nested t-test | 3 animals/condition; 10-15 microglia/animal | Males | Conditio n | t(73)=1.008 | 0.3166 |

Figure 4. Statistics

| | Comparison | Statistical test | n | Sexes included | Variable | Statistic | p-value |
|---|---|---|---|---|---|---|---|
| C. | Microglia Heterogeneity (P8, P15, P25): CON vs DEP+MS Male x age | Two-way ANOVA | n=3 animals/treatment/age ; 3 biological replicates/animal | Males | Condition | F (1, 50) = 14.87 | 0.0003 |
| | | | | | Age | F (2, 50) = 30.13 | <0.0001 |
| | | | | | Interaction | F (2, 50) = 4.296 | 0.019 |
| | P8: CON vs DEP+MS | Holm-Sidak's multiple comparisons test | | Males | Condition | t(50)=3.378 | 0.0043 |
| | P15: CON vs DEP+MS | Holm-Sidak's multiple comparisons test | | Males | Condition | t(50)=3.1243 | 0.0056 |
| | P25: CON vs DEP+MS | Holm-Sidak's multiple comparisons test | | Males | Condition | t(50)=0.0802 | 0.9364 |
| E. | Microglia Heterogeneity Phagocytic Index (P8): cell subtype | Nested one-way ANOVA | 5-7 cells/subtype/animal; 3 animals/condition | Males | Cell subtype | F(2, 19)=14.35 | 0.0002 |
| | Iba1 high vs. Both high | Iba1 high vs. Both high | | Males | Cell subtype | t(19)=0.9936 | 0.3329 |
| | P2ry12$^{hi}$ vs. Both high | P2ry12$^{hi}$ vs. Both high | | Males | Cell subtype | t(19)=5.054 | 0.0001 |
| G. | Microglial Heterogeneity Synapse engulfment (P8): cell subtype | Nested one-way ANOVA | 5-15 cells/subtype/animal; 3 animals/condition | Males | Cell subtype | F(2, 163)=13.49 | 0.0001 |
| | Iba1 high vs. Both high | Sidak's multiple comparisons test | | Males | Cell subtype | t(163)=1.512 | 0.2472 |
| | P2ry12$^{hi}$ vs. Both high | Sidak's multiple comparisons test | | Males | Cell subtype | t(163)=5.163 | <0.0001 |

Figure 5. Statistics

| | Comparison | Statistical test | n | Sexes include d | Variable | Statistic | p-value |
|---|---|---|---|---|---|---|---|
| C | Number of calls (<P60): CON vs DEP+MS Males | Unpaired t-test | n=15-17/animals/condition | Males | Condition | F(1, 8)=0.03957 | 0.8473 |
| D | Total call time (<P60) :CON vs DEP+MS Males | Unpaired t-test | n=15-17/animals/condition | Males | Condition | F(1, 8)=2.05 | 0.0416 |
| H | Number of calls (<P60): Microglia Intact vs ablated Males | Unpaired t-test | n=6-9 animals/genotype | Males | Genotype (ablation) | t(13)=0.9602 | 0.3545 |
| I | Total call time (<P60): Microglia Intact vs ablated Males | Unpaired t-test | n=6-9 animals/genotype | Males | Genotype (ablation) | t(13)=1.913 | 0.0799 |

#### Figure S1. Statistics

| | **Comparison** | **Statistical test** | ***n*** | ***Sexes include*** | ***Variable*** | **Statistic** | ***p*-value** |
|---|---|---|---|---|---|---|---|
| A | Pro-inflammatory Cytokine Analysis: CON vs DEP+MS dams | One-way ANOVA | 5-6 dams/condition | Females | Condition | t(9)=3.404 | 0.0078 |
| B | IL-10 dam serum: CON vs DEP+MS dams | Unpaired t-test | 5-6 dams/condition | Females | condition | t(9)=0.2781 | 0.7872 |
| C | Maternal CORT: CON vs DEP+MS dams vs untreated | One-way ANOVA | 4-6 dams/condition | Females | Condition | F (3, 12) = 27.24 | <0.0001 |
| | CON vs untreated pregnant dams | Holm-Sidak's post hoc | 4-5 dams/condition | Females | condition | t(12)=1.361 | 0.1987 |
| | CON vs DEP+MS dams | Holm-Sidak's post hoc | 5-6 dams/condition | Females | condition | t(12)=5.454 | 0.0004 |

#### Figure S2. Statistics

| | **Comparison** | **Statistical test** | ***n*** | ***Sexes included*** | ***Variable*** | **Statistic** | ***p*-value** |
|---|---|---|---|---|---|---|---|
| B | Maternal pregnancy weight gain: CON vs DEP+MS dams | unpaired t-test | 5-6 dams/condition | Females | condition | t(9)=0.2065 | 0.841 |
| D | Neonatal weight gain: CON vs DEP+MS pups | Two-way ANOVA | 31-33 pups/condition | Sexes combined | Condition | F (1, 64) = 14.84 | 0.0003 |
| | | | | | Age | F (4, 252) = 757.7 | <0.0001 |
| | | | | | Interaction | F (4, 252) = 2.798 | 0.0267 |
| | P7: CON vs DEP+MS | Sidak's multiple comparisons test | 31-33 pups/condition | sexes combined | condition | t(316)=2.582 | 0.0136 |
| | P8: CON vs DEP+MS | Sidak's multiple comparisons test | 31-33 pups/condition | sexes combined | condition | t(316)=2.722 | 0.0136 |
| | P9: CON vs DEP+MS | Sidak's multiple comparisons test | 31-33 pups/condition | sexes combined | condition | t(316)=3.606 | 0.0011 |
| | P10: CON vs DEP+MS | Sidak's multiple comparisons test | 31-33 pups/condition | sexes combined | condition | t(316)=3.849 | 0.0006 |
| | P15: CON vs DEP+MS | Sidak's multiple comparisons test | 31-33 pups/condition | sexes combined | condition | t(316)=4.577 | <0.0001 |

Figure S3. Statistics

| | Comparison | Statistical test | *n* | *Sexes included* | *Variable* | Statistic | *p*-value |
|---|---|---|---|---|---|---|---|
| A | WT USVs characterization across development | Two-way ANOVA | 4-10 pups/group/sex | Male and females | Age | F (5, 80) = 2.808 | 0.0218 |
| | | | | | Sex | F (1, 80) = 0.1014 | 0.751 |
| | | | | | Interaction | F (5, 80) = 0.2454 | 0.9414 |
| | P4: P5 | Holm-Sidak's post hoc | 9-16 pups/age | sexes combined | age | t(80)=1.29 | 0.3111 |
| | P4: P6 | Holm-Sidak's post hoc | 9-23 pups/age | sexes combined | age | t(80)=1.894 | 0.1743 |
| | P4: P7 | Holm-Sidak's post hoc | 9-22 pups/age | sexes combined | age | t(80)=2.345 | 0.0833 |
| | P4:P8 | Holm-Sidak's post hoc | 9-14 pups/age | sexes combined | age | t(80)=3.492 | 0.0039 |
| | P4:P9 | Holm-Sidak's post hoc | 8-9 pups/age | sexes combined | age | t(80)=1.385 | 0.3111 |
| B | Number of calls across development | Mixed-effects model (REML) | 33-35 pups/condition | sexes combined | condition | F (1, 34) = 21.52 | <0.0001 |
| | | | | | age | F (2, 68) = 4.373 | 0.0164 |
| | | | | | interaction | F (2, 53) = 1.931 | 0.155 |
| | P7: CON vs DEP+MS | Holm-Sidak's post hoc | 33-35 pups/condition | sexes combined | condition | t(87)=3.94 | 0.0005 |
| | P8: CON vs DEP+MS | Holm-Sidak's post hoc | 33-35 pups/condition | sexes combined | condition | t(87)=3.805 | 0.0005 |
| | P9: CON vs DEP+MS | Holm-Sidak's post hoc | 33-35 pups/condition | sexes combined | condition | t(87)=1.614 | 0.1101 |
| C | Total time calling across development | Mixed-effects model (REML) | 33-35 pups/condition | sexes combined | condition | F (1, 66) = 6.767 | 0.0114 |
| | | | | | age | F (2, 123) = 3.205 | 0.044 |
| | | | | | interaction | F (2, 123) = 0.9532 | 0.3883 |
| | P7: CON vs DEP+MS | Holm-Sidak's post hoc | 33-35 pups/condition | sexes combined | condition | t(189)=2.315 | 0.043 |
| | P8: CON vs DEP+MS | Holm-Sidak's post hoc | 33-35 pups/condition | sexes combined | condition | t(189)=2.466 | 0.043 |
| | P9: CON vs DEP+MS | Holm-Sidak's post hoc | 33-35 pups/condition | sexes combined | condition | t(189)=0.9523 | 0.3421 |
| D | Call frequency across development | Mixed-effects model (REML) | 33-35 pups/condition | sexes combined | condition | F (1, 63) = 0.1303 | 0.7194 |
| | | | | | age | F (2, 122) = 14.43 | <0.0001 |
| | | | | | interaction | F (2, 122) = 2.580 | 0.0799 |
| | P7: CON vs DEP+MS | Holm-Sidak's post hoc | 33-35 pups/condition | sexes combined | condition | t(185)=0.5541 | 0.8238 |
| | P8: CON vs DEP+MS | Holm-Sidak's post hoc | 33-35 pups/condition | sexes combined | condition | t(185)=0.1789 | 0.8583 |
| | P9: CON vs DEP+MS | Holm-Sidak's post hoc | 33-35 pups/condition | sexes combined | condition | t(185)=1.324 | 0.463 |

#### Figure S5. Statistics

| | comparison | NAME | n | sexes included | NES | FDR q-val | NOM p-val |
|---|---|---|---|---|---|---|---|
| A | GSEA Hallmark Pathways: Upregulated males | HALLMARK_EPITHELIAL_MESENCHYMAL_TRANSITION | 4 animals/condition | Males | 2.162075 | 0 | 0 |
| | | HALLMARK_MITOTIC_SPINDLE | | Males | 2.0154824 | 0 | 0 |
| | | HALLMARK_ANGIOGENESIS | | Males | 1.937755 | 6.78E-04 | 0 |
| | | HALLMARK_WNT_BETA_CATENIN_SIGNALING | | Males | 1.9177881 | 7.84E-04 | 0.0019305 |
| | | HALLMARK_G2M_CHECKPOINT | | Males | 1.904847 | 0.00104114 | 0 |
| | | HALLMARK_NOTCH_SIGNALING | | Males | 1.8360845 | 0.00166879 | 0.00182815 |
| | | HALLMARK_E2F_TARGETS | | Males | 1.5811505 | 0.02251368 | 0.00157978 |
| | | HALLMARK_UV_RESPONSE_DN | | Males | 1.5089926 | 0.0432611 | 0.00505051 |
| | | HALLMARK_INTERFERON_GAMMA_RESPONSE | | Males | 1.5072445 | 0.03893471 | 0.00490998 |
| B. | GSEA Hallmark Pathways: Upregulated females | HALLMARK_MYC_TARGETS_V1 | 4 animals/condition | Females | 2.6254957 | 0 | 0 |
| | | HALLMARK_OXIDATIVE_PHOSPHORYLATION | | Females | 2.3127167 | 0 | 0 |
| | | HALLMARK_DNA_REPAIR | | Females | 2.1645634 | 0 | 0 |
| | | HALLMARK_INTERFERON_ALPHA_RESPONSE | | Females | 1.8586267 | 0.00212617 | 0.00168634 |
| | | HALLMARK_E2F_TARGETS | | Females | 1.7030339 | 0.00610582 | 0 |
| | | HALLMARK_MYC_TARGETS_V2 | | Females | 1.60789 | 0.01213864 | 0.00905797 |
| | | HALLMARK_INTERFERON_GAMMA_RESPONSE | | Females | 1.4548359 | 0.04686008 | 0.01594896 |
| C. | GSEA Hallmark Pathways: Downregulated males | HALLMARK_OXIDATIVE_PHOSPHORYLATION | 4 animals/condition | Males | -2.1912339 | 0 | 0 |
| | | HALLMARK_MYC_TARGETS_V1 | | Males | -1.6196603 | 0.02915523 | 0 |
| | | HALLMARK_PEROXISOME | | Males | -1.5772246 | 0.0284296 | 0.00956938 |
| | | HALLMARK_HEME_METABOLISM | | Males | -1.5042268 | 0.04180964 | 0.00265957 |
| | | HALLMARK_FATTY_ACID_METABOLISM | | Males | -1.4710518 | 0.04401244 | 0.00476191 |
| D. | GSEA Hallmark Pathways: Downregulated females | HALLMARK_EPITHELIAL_MESENCHYMAL_TRANSITION | 4 animals/condition | Females | -2.452149 | 0 | 0 |
| | | HALLMARK_UV_RESPONSE_DN | | Females | -2.0402446 | 0 | 0 |
| | | HALLMARK_COAGULATION | | Females | -1.9143175 | 0.00204379 | 0 |

| | | | | | | | |
|---|---|---|---|---|---|---|---|
| | | HALLMARK_CHOLESTEROL_HOMEOSTASIS | | Females | -1.8883774 | 0.00187375 | 0 |
| | | HALLMARK_ANGIOGENESIS | | Females | -1.7585561 | 0.00561232 | 0 |

#### Figure S8. Statistics

| | **Comparison** | **Statistical test** | ***n*** | ***Sexes included*** | ***Variable*** | **Statistic** | ***p*-value** |
|---|---|---|---|---|---|---|---|
| C. Left | VGlut2 Synapse (P8): Females, Treatment | Nested t-test | 3 animals/condition; 3 sections/animal; 5 z-stacks/section | Females | condition | t(28)=1.696 | 0.0925 |
| C. Middle | VGlut2 Synapse (P15): Females, Treatment | Nested t-test | 3 animals/condition; 3 sections/animal; 5 z-stacks/section | Females | condition | t(28)=2.210 | 0.0297 |
| C. Right | VGlut2 Synapse (P100): Females, Treatment | Nested t-test | 3 animals/condition; 3 sections/animal; 5 z-stacks/section | Females | condition | t(28)=0.9387 | 0.3823 |

#### Figure S10. Statistics

| | **Comparison** | **Statistical test** | ***n*** | ***Sexes included*** | ***Variable*** | **Statistic** | ***p*-value** |
|---|---|---|---|---|---|---|---|
| A. Top | ACC NeuN Density: Condition x Sex | Two-way ANOVA | 3 animals/condition/sex; 3 sections/animal | Males and females | condition | F (1, 41) = 0.2346 | 0.6307 |
| | | | | | sex | F (1, 41) = 0.5149 | 0.4771 |
| | | | | | interaction | F (1, 41) = 0.001858 | 0.9658 |
| A. Second from top | ACC Olig2 Density: Condition x Sex | Two-way ANOVA | 3 animals/condition/sex; 3 sections/animal | Males and females | condition | F (1, 41) = 1.519 | 0.2248 |
| | | | | | sex | F (1, 41) = 2.656 | 0.1108 |
| | | | | | interaction | F (1, 41) = 0.7852 | 0.3807 |
| A. Third from top | ACC Sox9 Density: Condition x Sex | Two-way ANOVA | 3 animals/condition/sex; 3 sections/animal | Males and females | condition | F (1, 41) = 2.730 | 0.1061 |
| | | | | | sex | F (1, 41) = 0.01303 | 0.9097 |
| | | | | | interaction | F (1, 41) = 1.031 | 0.3159 |
| A. Bottom | ACC Olig2+/Sox9+ Density: Condition x Sex | Two-way ANOVA | 3 animals/condition/sex; 3 sections/animal | Males and females | condition | F (1, 41) = 3.122 | 0.0847 |
| | | | | | sex | F (1, 41) = 1.311 | 0.2893 |
| | | | | | interaction | F (1, 41) = 1.153 | 0.2588 |

#### Figure S11. Statistics

| | Comparison | Statistical test | *n* | *Sexes included* | *Variable* | Statistic | *p*-value |
|---|---|---|---|---|---|---|---|
| A-B | Microglia CD68 distribution: CON vs DEP+MS | Levene's test for homogeneity of variance | 3 animals/condition; 5-15 microglia/animal | Males | Condition | F(1, 73)=6.6787 | 0.0117 |
| | | | | Males | | | |

#### Figure S12. Statistics

| | Comparison | Statistical test | *n* | *Sexes included* | *Variable* | Statistic | *p*-value |
|---|---|---|---|---|---|---|---|
| B | Microglia density across development | Two-way ANOVA | 3 animals/condition; 3 sections/animal | Males | Condition | F (1, 66) = 0.2121 | 0.6466 |
| | | | | males | age | F (2, 66) = 57.30 | <0.0001 |
| | | | | males | interaction | F (2, 66) = 0.4212 | 0.658 |

#### Figure S14. Statistics

| | Comparison | Statistical test | *n* | *Sexes included* | *Variable* | Statistic | *p*-value |
|---|---|---|---|---|---|---|---|
| A. Left | CON Paired microglia heterogeneity Cd68 | Nested One-way ANOVA | 3 animals; 5-7 cells/animal/subtype; (n=59 cells total) | males | Cell subtype | F(2, 56)=9.357 | 0.0003 |
| | Both high vs Iba1 high | Holm-Sidak's post hoc | n=20-21 cells/subtype | males | cell subtype | t(56)=0.2087 | 0.8355 |
| | Both high vs P2ry12 high | Holm-Sidak's post hoc | n=18-20 cells/subtype | males | cell subtype | t(56)=3.863 | 0.0006 |
| A. Right | DEP+MS Paired microglia heterogeneity CD68 | Nested One-way ANOVA | 3 animals; 5-7 cells/animal/subtype; (n=61 cells total) | males | Cell subtype | F(2,19)=7.078 | 0.005 |
| | Both high vs Iba1 high | Holm-Sidak's post hoc | n=20 cells/subtype | males | cell subtype | t(19)=1.271 | 0.2191 |
| | Both high vs P2ry12 high | Holm-Sidak's post hoc | n=20-21 cells/subtype | males | cell subtype | t(19)=3.697 | 0.0031 |
| B. Left | Volume of microglia-Both high | unpaired-t test | 3 animals; 5-15 cells/animal; n=72 cells total | Males | Condition | t(4)=0.6075 | 0.5548 |
| B. Middle | Volume of microglia-Iba1 high | unpaired-t test | 3 animals; 5-10 cells/animal; n=44 cells total | Males | Condition | t(4)=0.08630 | 0.9936 |
| B. Right | Volume of microglia-P2ry12 high | unpaired-t test | 3 animals; 5-11 cells/animal; n=52 cells total | Males | Condition | t(4)=1.043 | 0.3174 |
| C. Left | TC engulfment- both high | unpaired-t test | 3 animals; 5-15 cells/animal; n=72 cells total | Males | Condition | t(4)=0.5811 | 0.5719 |
| C. Middle | TC engulfment- iba1 high | unpaired-t test | 3 animals; 5-10 cells/animal; n=44 cells total | Males | Condition | t(4)=1.801 | 0.0792 |
| C. Right | TC engulfment- p2ry12 high | unpaired-t test | 3 animals; 5-11 cells/animal; n=52 cells total | Males | Condition | t(4)=0.5888 | 0.4577 |

#### Figure S15 Statistics

| | Comparison | Statistical test | *n* | *Sexes included* | *Variable* | Statistic | *p*-value |
|---|---|---|---|---|---|---|---|
| A. Left | CON Paired microglia heterogeneity engulfment | Nested One-way ANOVA | 3 animals; 5-15 cells/animal/subtype; (n=89 cells total) | males | Cell subtype | F (2, 40) = 8.344 | 0.0009 |
| | Both high vs Iba1 high | Holm-Sidak's post hoc | n=19-39 cells | males | | t(40)=2.591 | 0.0133 |
| | Both high vs P2ry12 high | Holm-Sidak's post hoc | n=31-39 cells | males | | t(40)=3.906 | 0.0007 |
| A. Right | DEP+MS Paired microglia heterogeneity engulfment | Nested One-way ANOVA | 3 animals; 5-15 cells/animal/subtype; (n=76 cells total) | males | Cell subtype | F (2, 73) = 5.563 | 0.0056 |
| | Both high vs Iba1 high | Holm-Sidak's post hoc | n=24-32 cells | males | | t(29)=0.4195 | 0.6761 |
| | Both high vs P2ry12 high | Holm-Sidak's post hoc | n=20-32 cells | males | | t(29)=3.194 | 0.0041 |
| B. Left | Volume of microglia-Both high | unpaired-t test | 3 animals; 5-15 cells/animal; n=72 cells total | Males | Condition | t(4)=1.420 | 0.2285 |
| B. Middle | Volume of microglia-Iba1 high | unpaired-t test | 3 animals; 5-10 cells/animal; n=44 cells total | Males | Condition | t(4)=8.422 | 0.0011 |
| B. Right | Volume of microglia-P2ry12 high | unpaired-t test | 3 animals; 5-11 cells/animal; n=52 cells total | Males | Condition | t(4)=1.405 | 0.2328 |
| C. Left | TC engulfment- both high | unpaired-t test | 3 animals; 5-15 cells/animal; n=72 cells total | Males | Condition | t(4)=0.8274 | 0.4545 |
| C. Middle | TC engulfment- iba1 high | unpaired-t test | 3 animals; 5-10 cells/animal; n=44 cells total | Males | Condition | t(4)=1.581 | 0.189 |
| C. Right | TC engulfment- p2ry12 high | unpaired-t test | 3 animals; 5-11 cells/animal; n=52 cells total | Males | Condition | t(4)=1.872 | 0.1345 |

#### Figure S16. Statistics

| | Comparison | Statistical test | *n* | *Sexes included* | *Variable* | Statistic | *p*-value |
|---|---|---|---|---|---|---|---|
| A. | Syllable Length: Adult CON vs DEP+MS | unpaired t-test | 15-17 animals | males | condition | t(30)=2.753 | 0.0101 |
| B. | USV frequency: Adult CON vs DEP+MS | unpaired t-test | 15-17 animals | males | condition | t(30)=0.5150 | 0.6102 |

Author contributions:

| | |
|---|---|
| **Carina L. Block** | Jointly conceived of experiments with SDM, KD, CE and SDB; generated and collected all experimental mice for behavior, transcriptomic, histology and neurophysiological experiments with OE and KEM; jointly collected maternal serum with OE and KEM; jointly performed PFC dissections and extracted RNA with KEM; jointly performed histological confirmations for electrode implantation experiments with, SDM, OE, NN, TN, and CB; jointly collected and analyzed neonatal and adult USV behavior with OE and CS; supervised analysis of WT synapse density and engulfment performed by KAB; jointly performed P10 engulfment analysis with KAB; supervised implementation of u-net for image restoration and cell counts; jointly performed analysis of cell counts with CS; jointly performed IHC and imaging with OE for microglia analysis; quantified and analyzed microglia density, heterogeneity and Imaris reconstructions of CD68 and VGlut2 in heterogeneous microglia; jointly performed MUPET analysis of neonatal and adult USVs with CS; performed ablation experiments and characterization; jointly performed genotyping with OE; jointly collected USVs in ablated animals with KEM; jointly collected behavior and tissue with LJC. Jointly wrote and edited the paper with SDB, KD, CE, SDM. |
| **Oznur Eroglu** | Jointly generated and collected experimental animals with CLB and KEM; jointly collected maternal serum with CLB and KEM; jointly performed histological confirmations for electrode implantation experiments with SDM, CLB, NN, TN, and CB; jointly collected and analyzed neonatal USV behavior with CLB and CS; jointly performed IHC and imaging with CLB for microglia and synapse analysis; jointly performed genotyping with CLB. |
| **Stephen D. Mague** | Jointly conceived two-chamber social interaction experiment; jointly built electrodes with CB and KD; jointly performed electrode implantations in DEP and Control mice for two-chamber social interaction experiment with CB; jointly analyzed behavioral data and performed neurophysiological data processing for two-chamber social interaction experiments with CB, NN, and NMG; jointly performed histological confirmations for electrode implantation experiments with CLB, OE, NN, TN, and CB; edited the paper. |
| **Chaichontat Sriworarat** | Jointly analyzed neonatal and adult USV behavior with CLB; implemented MUPET analysis of USVs; generated lab specific code |

| | |
|---|---|
| | for u-net image restoration and cell counts; jointly performed analysis of cell counts with CLB. |
| **Cameron Blount** | Built electrodes for mice two-chamber social interaction test and EPM experiments; jointly implanted mice for EPM experiment with SDM; jointly collected data in DEP+MS and CON mice for two-chamber social interaction test experiment with NN; jointly analyzed behavioral data for two-chamber social interaction experiments and performed neurophysiological data processing for all experiments presented in the paper with SDM, NN, and NMG; jointly performed histological confirmations for electrode implantation experiments with SDM, CLB, OE, NN, TN. |
| **Karen E. Malacon** | Jointly generated experimental mice with CLB and OE; jointly collected maternal serum with CLB and OE; jointly performed PFC dissections and extracted RNA with CLB; jointly collected USVs in ablated animals with CLB. |
| **Kathleen A. Beben** | Jointly performed analysis of WT synapse density and engulfment with CLB and OE; jointly performed P10 engulfment analysis with CLB. |
| **Nkemdilim Ndubuizu** | Jointly collected data in DEP+MS and CON mice for two-chamber social preference experiment with CB; jointly analyzed behavioral data for two-chamber social interaction experiments and performed neurophysiological data processing for all experiments presented in the paper with CB, SDM, and NMG; jointly performed histological confirmations for electrode implantation experiments with SDM, CLB, OE, TN, and CB. |
| **Austin Talbot** | Developed and implemented the CSFA-NMF machine learning analyses utilized for all neurophysiological analysis in the paper including the model discovery and projections of new neurophysiological data into the initial model space. |
| **Neil M. Gallagher** | Implemented granger features for data processing; jointly contributed to data processing with SDM, CB, and NN. |
| **Young Chan Jo** | Jointly collected behavior and tissue in ablation experiment with CLB. |
| **Timothy Nyangacha** | Jointly performed histological confirmations for all experiments with SDM, CLB, OE, NN, and CB. |
| **David E. Carlson** | Supervised development of all aspects of CSFA-NMF machine learning analyses and data processing methodology. |

| | |
|---|---|
| **Kafui Dzirasa** | Jointly conceived of experiments with CLB, SDM, CE and SDB; jointly analyzed neurophysiological data with AT; supervised all behavioral and neurophysiological experiments with SDM; edited the paper. |
| **Cagla Eroglu** | Jointly conceived of experiments with CLB, SDM, SDB and KD; supervised all behavioral and cellular and molecular experiments with CLB and SDB; wrote the paper jointly with CLB and SDB. |
| **Staci D. Bilbo** | Jointly conceived of experiments with CLB, SDM, KD and CE; supervised all behavioral and cellular and molecular experiments with CLB and CE; wrote the paper jointly with CLB and CE. |

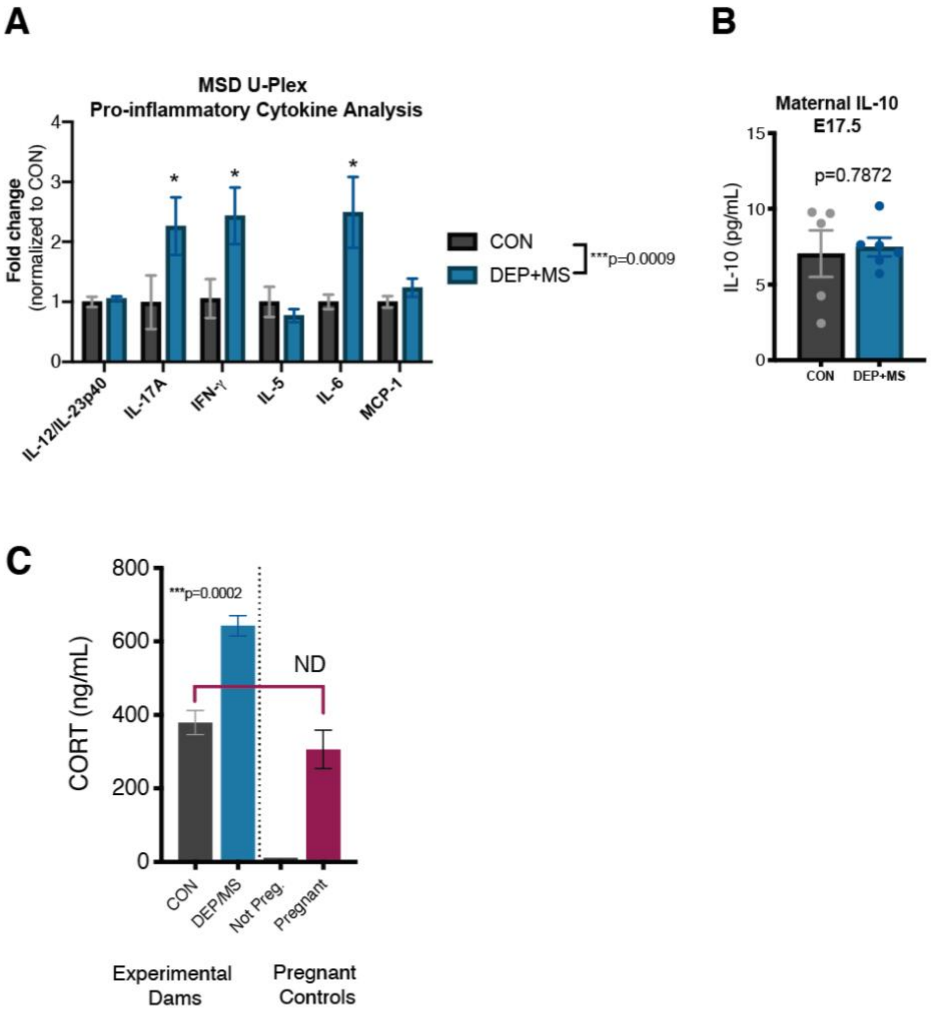

**Fig. S1. Pro-inflammatory cytokines and CORT measurements in pregnant dams and control animals. (A-B)** Serum concentrations of pro-inflammatory cytokines (A) and anti-inflammatory cytokine (B) in E17.5 dams 2 hours post last instillation (n=5-6 animals/condition), DEP+MS dams have elevated expression of pro-inflammatory cytokines (A), but no significant differences in IL-10. (**C**) Corticosterone measurements for experimental dams and non-treated non-pregnant and pregnant dams, CORT levels for CON are no higher than pregnant dams at baseline (n=4-6 mice/condition). Two-way ANOVA with Holm-Sidak's post hoc test (A), Unpaired t-test (B), One-way ANOVA with Holm-Sidak's post hoc test (C). $***P<0.001$, $*P < 0.05$; ND=not different. Means ± SEM.

A

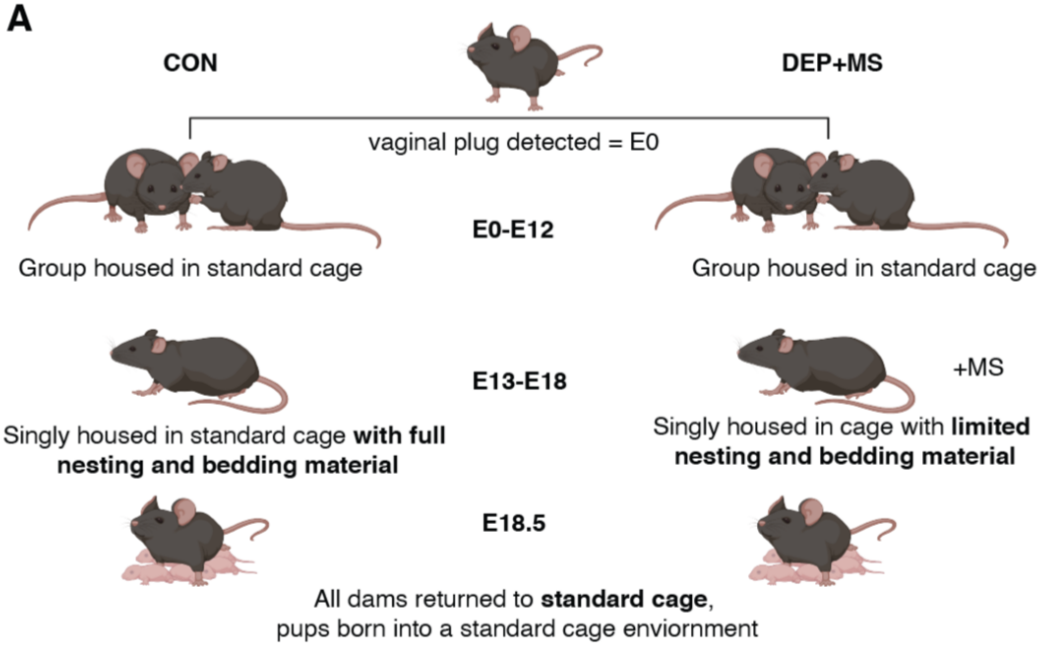

B

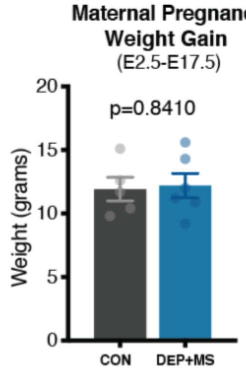

C

| | CON | DEP/MS |
|---|---|---|
| Total offspring | 31 | 33 |
| Male | 17 | 17 |
| Female | 14 | 16 |
| Percent Male | 54.84% | 51.52% |
| Average litter size | 7 | 8.25 |

D

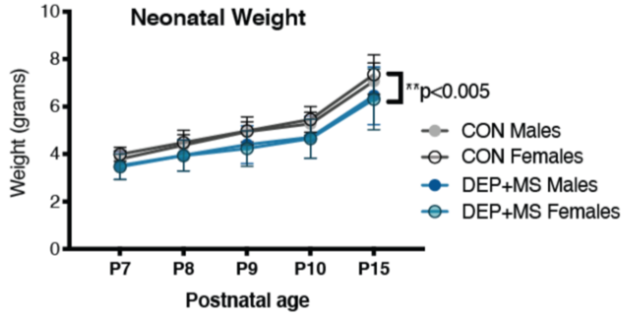

**Fig. S2 Maternal and neonatal outcomes**. (**A**) Females were time-mated and group house in standard cages from E0-E12, CON females (left) were singly housed in standard cages from E13-E18, while DEP+MS females (right) were singly-housed in nest restricted environment, all females returned to standard caging environment before the birth of pups. (**B**) There were no significant differences in maternal pregnancy weight gain from E2.5 to E17.5 (n=5-6 mice/condition). (**C**) No differences in litter size or litter sex composition. (**D**) DEP+MS offspring weighed significantly less than CON pups (n=14-17 mice/condition/sex). Unpaired t-test (B), Two-way ANOVA, condition x age, with Sidak's multiple comparison post hoc test (D). $^{**}P<0.01$; Means ± SEM.

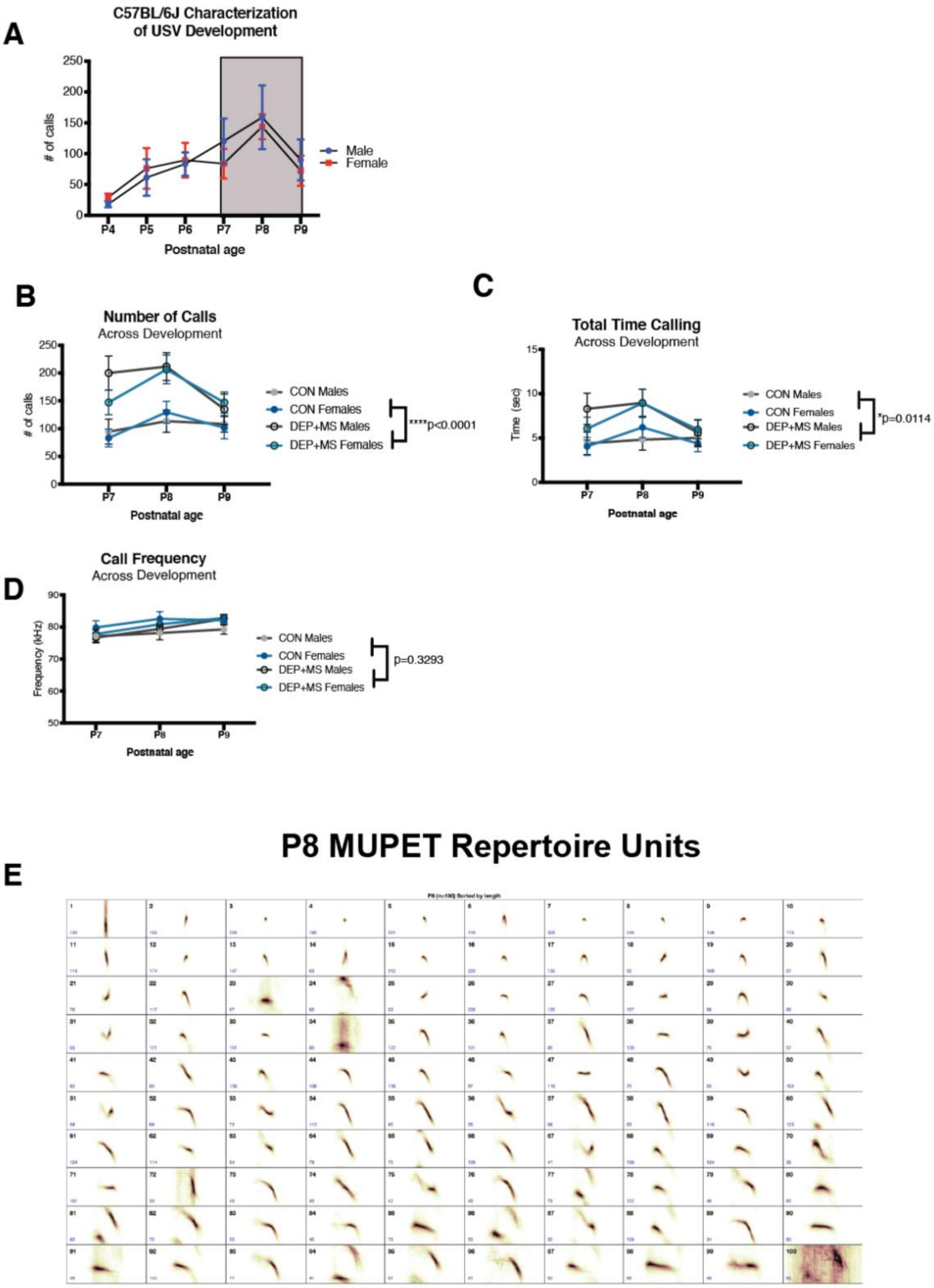
A
C57BL/6J Characterization
of USV Development
### of calls
Postnatal age
P4
P5
P6
P7
P8
P9
Male
Female
B
Number of Calls
Across Development
### of calls
Postnatal age
CON Males
CON Females
DEP+MS Males
DEP+MS Females
****p<0.0001
C
Total Time Calling
Across Development
Time (sec)
Postnatal age
CON Males
CON Females
DEP+MS Males
DEP+MS Females
*p=0.0114
D
Call Frequency
Across Development
Frequency (kHz)
Postnatal age
CON Males
CON Females
DEP+MS Males
DEP+MS Females
p=0.3293
E
P8 MUPET Repertoire Units

**Fig. S3 USV development in untreated C57BL/6J animals and experimental pups.** (**A**) Characterization of USV development in WT C57BL/6J mice (n=4-10 mice/group), no differences in number of USV between males and females across development, number of calls peaks at P8. (**B-C**) Number of calls and total time calling is significantly increased in DEP+MS offspring, but does not differ in developmental pattern (P7-P9). (**C**) no significant differences in frequency of calls (for B-D: n=14-17 mice/condition/sex). (**E**). P8 USV MUPET repertoire unit (RU) clustering, organized from shortest to longest, compiled from all CON and DEP+MS USV files. Two-way ANOVA, sex x age, with Holm Sidak's post hoc tests (A), Mixed effects model, condition x age, with Holm Sidak's post hoc tests, condition x age (B, C, D). ***$P$<0.001, **$P$ < 0.01; *$P$ < 0.05; Means ± SEM.

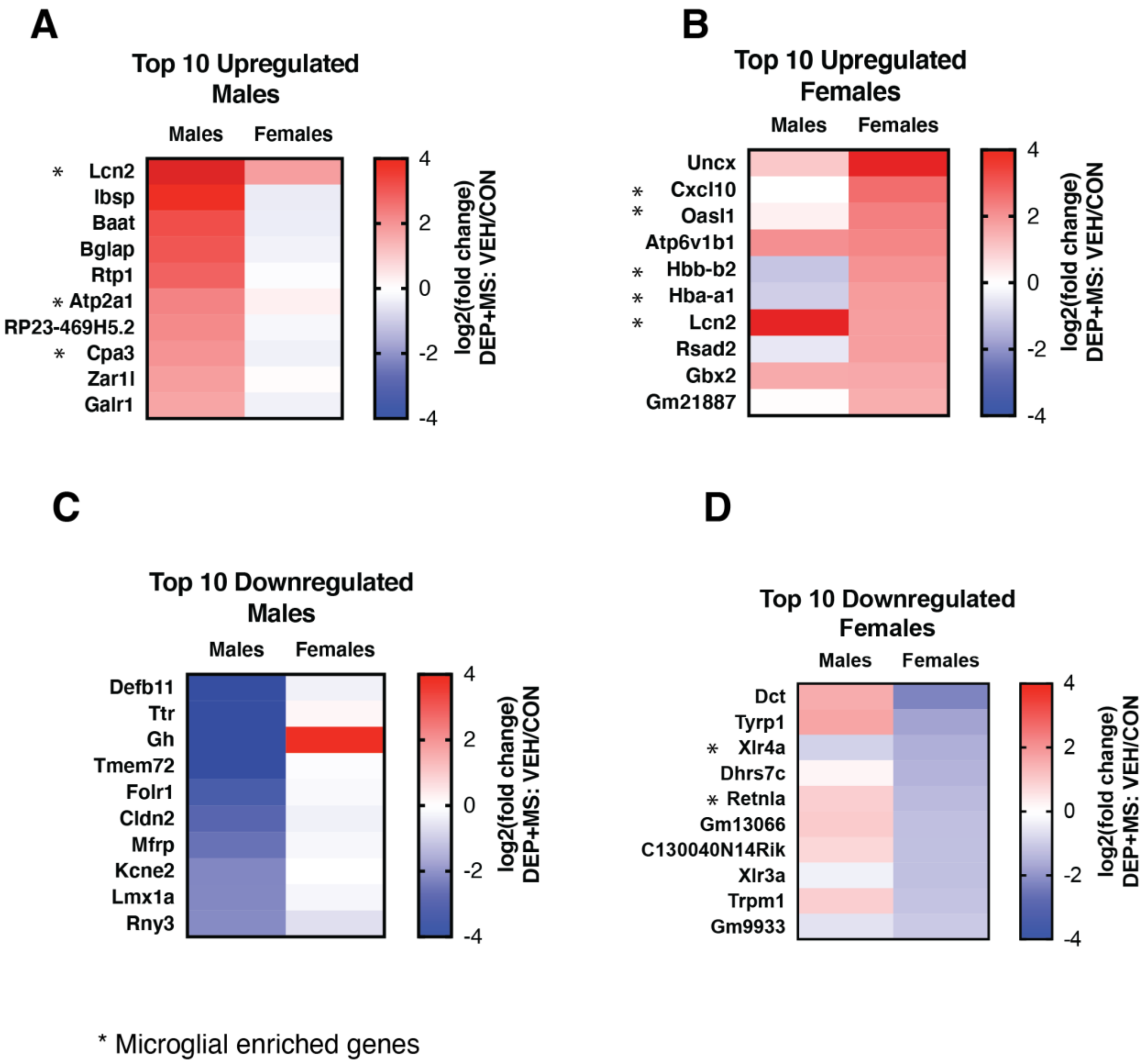

**Fig. S4. Microglia genes are enriched in differentially expressed genes.** (**A-B**) The top 10 differentially expressed genes that are upregulated in males (left) vs females (right), pattern of expression is sexually dimorphic. Asterisk denotes gene enrichment in microglia cells compared to other cells in the brain (compared using: brainrnaseq.org). (**C-D**) The top 10 differentially expressed genes that are downregulated in males (left) vs females (right), downregulated genes are also sexually dimorphic.

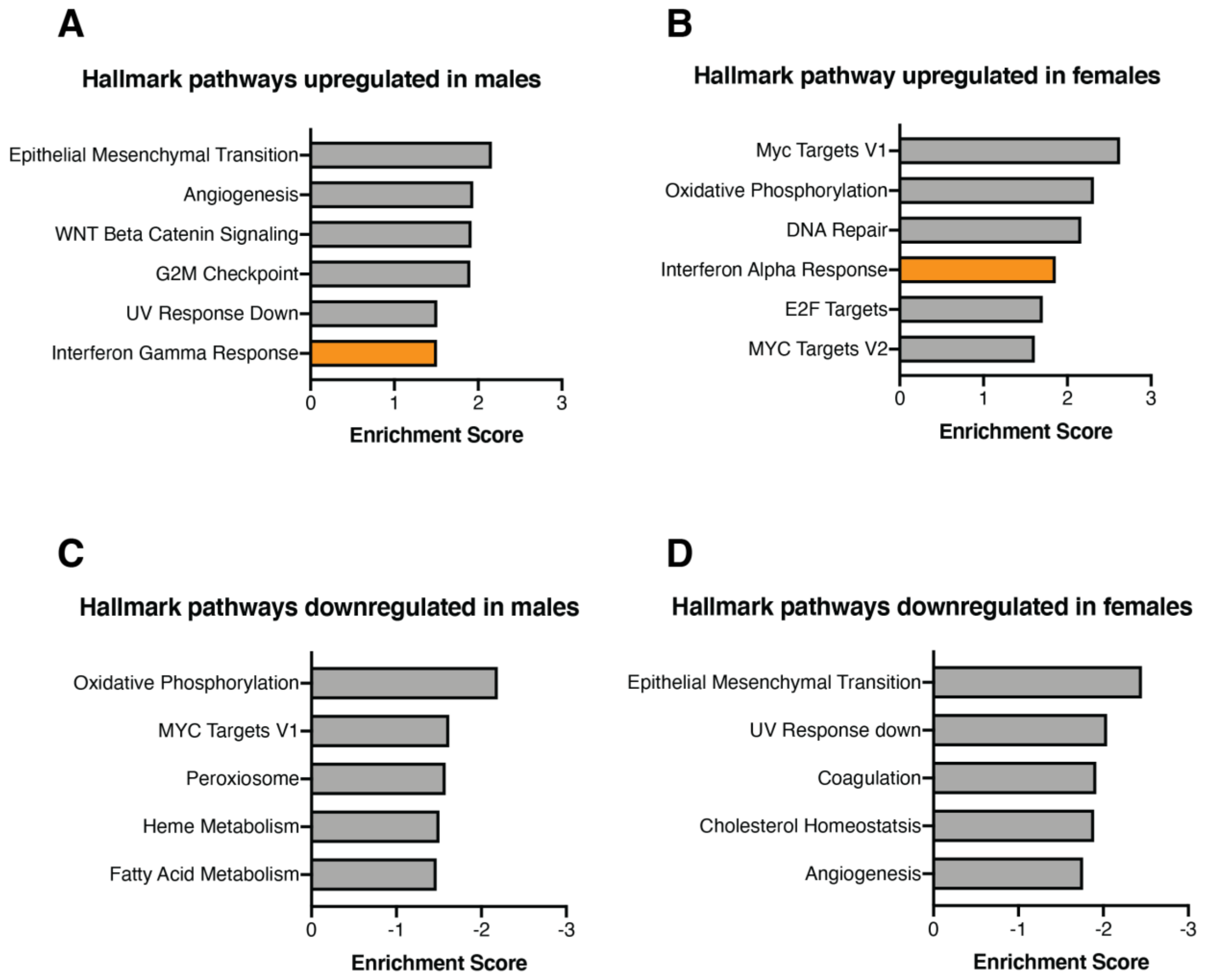

**Fig. S5 Immune pathways are upregulated in male and female DEP+MS offspring.** (**A-B**) Gene set enrichment analyses of upregulated hallmark pathways in males and females, males upregulated interferon gamma response, whereas females upregulated interferon alpha response. (**C-D**) Gene set enrichment analyses of downregulated hallmark pathways in males and females. Normalized enrichment scores of Molecular Signatures Database Hallmark from Gene Sets from Gene Set Enrichment Analysis (A-D), FDR q-value < 0.05.

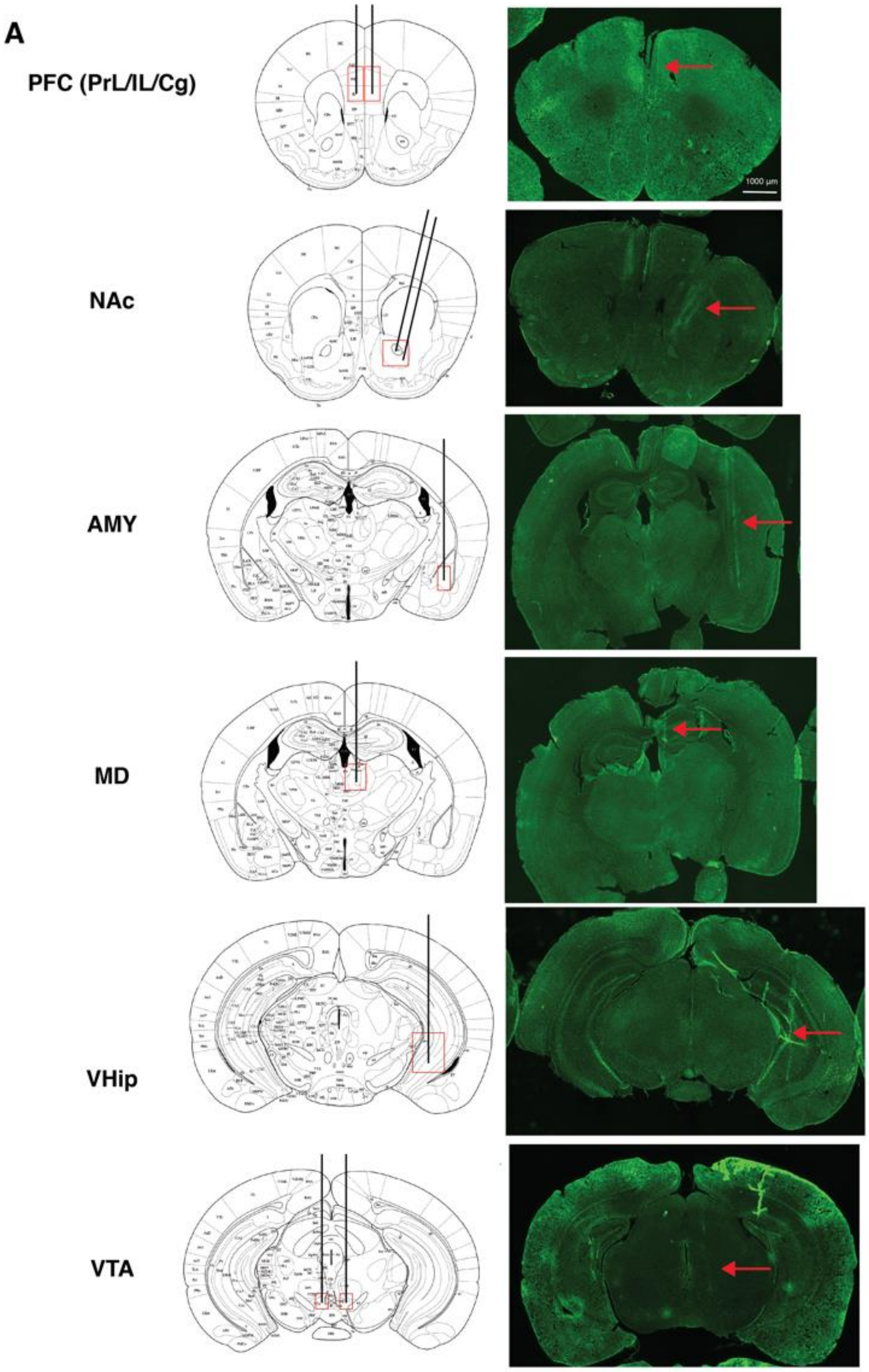

**Fig. S6 Histological confirmation of electrode placement.** (**A**) (Left) Electrode bundles (black lines) were centered within the red boxes on brain atlas image, electrodes were implanted bilateral for PFC and VTA. (Right) Exhaustive histology was performed on implanted brains to verify electrode placement, representative images of track verification on brains labeled with Nissl.

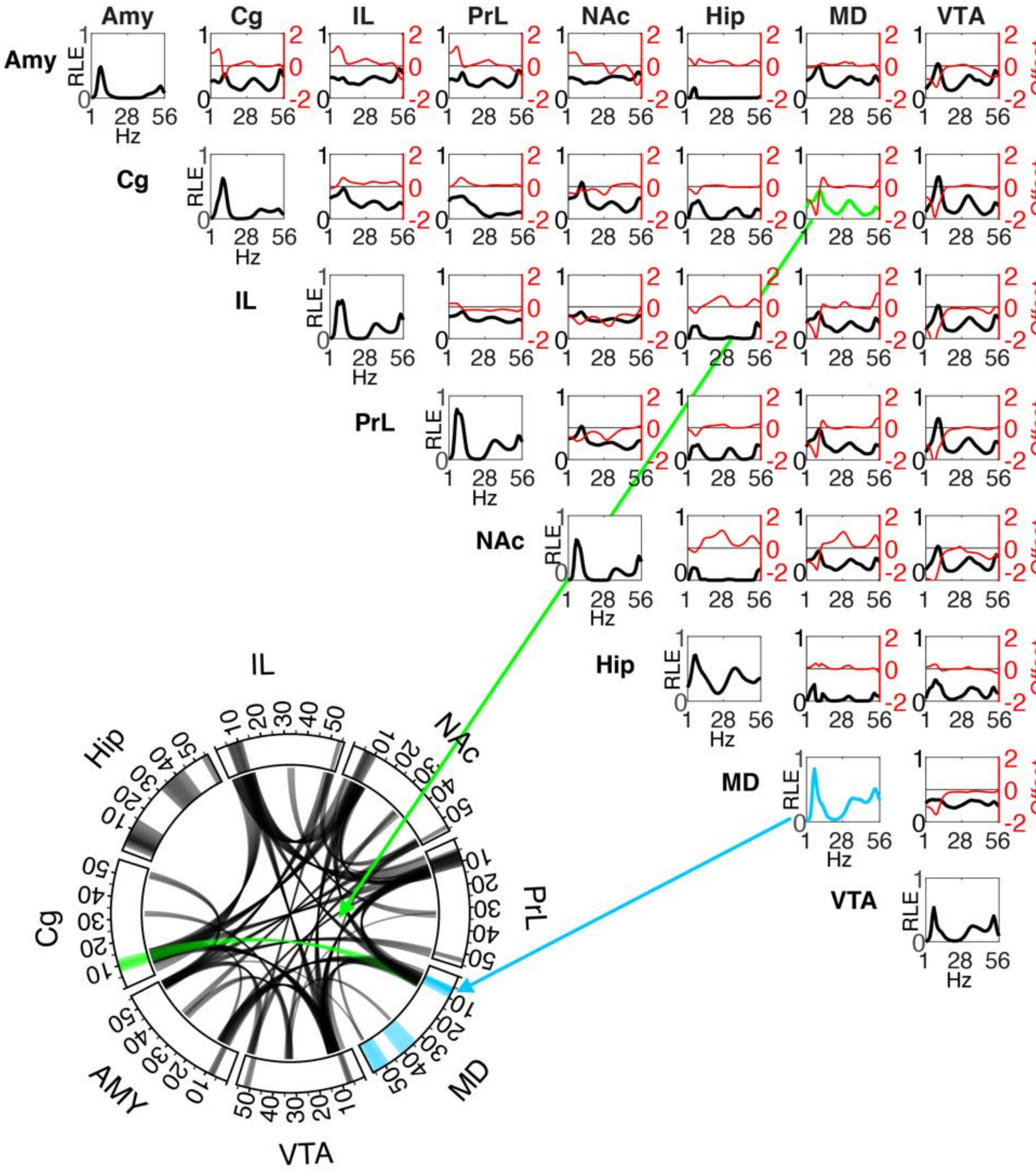
Amy
Cg
IL
PrL
NAc
Hip
MD
VTA
RLE
Offset
Hz
AMY

**Fig. S7 Power and synchrony measures that define the Social Electome Network adapted from Mague et al. 2020**. Brain areas are shown the top and the left identifying power and synchrony density function for Electome Factor 1. Amplitude values (shown in black) reflect the relative LFP spectral energy (RLE) observed at each frequency, where the Electome Factor is normalized to the total energy observed across the 6 networks. The offset between the two non-normalized granger synchrony fucntions for each brain area pair (A → B and B → A) are also shown in red (i.e. directionality; axis scale to the right). Positive spectral offsets correspond to frequencies at which the area listed along the top leads the area listed on the left. Negative spectral offsets correspond to the frequencies at which the area listed on the left leads the area listed on the top. The circular plot depicts the frequencies for power (outer rim) and synchrony (curved lines connecting two regions) above an amplitude threshold of 0.33. As a representative example, the power measures for the thalamus are highlighted in cyan in both the circular and correlation plots; synchrony between (PrL) and Medial Dorsal Thalamus (MD) are highlighted in green.

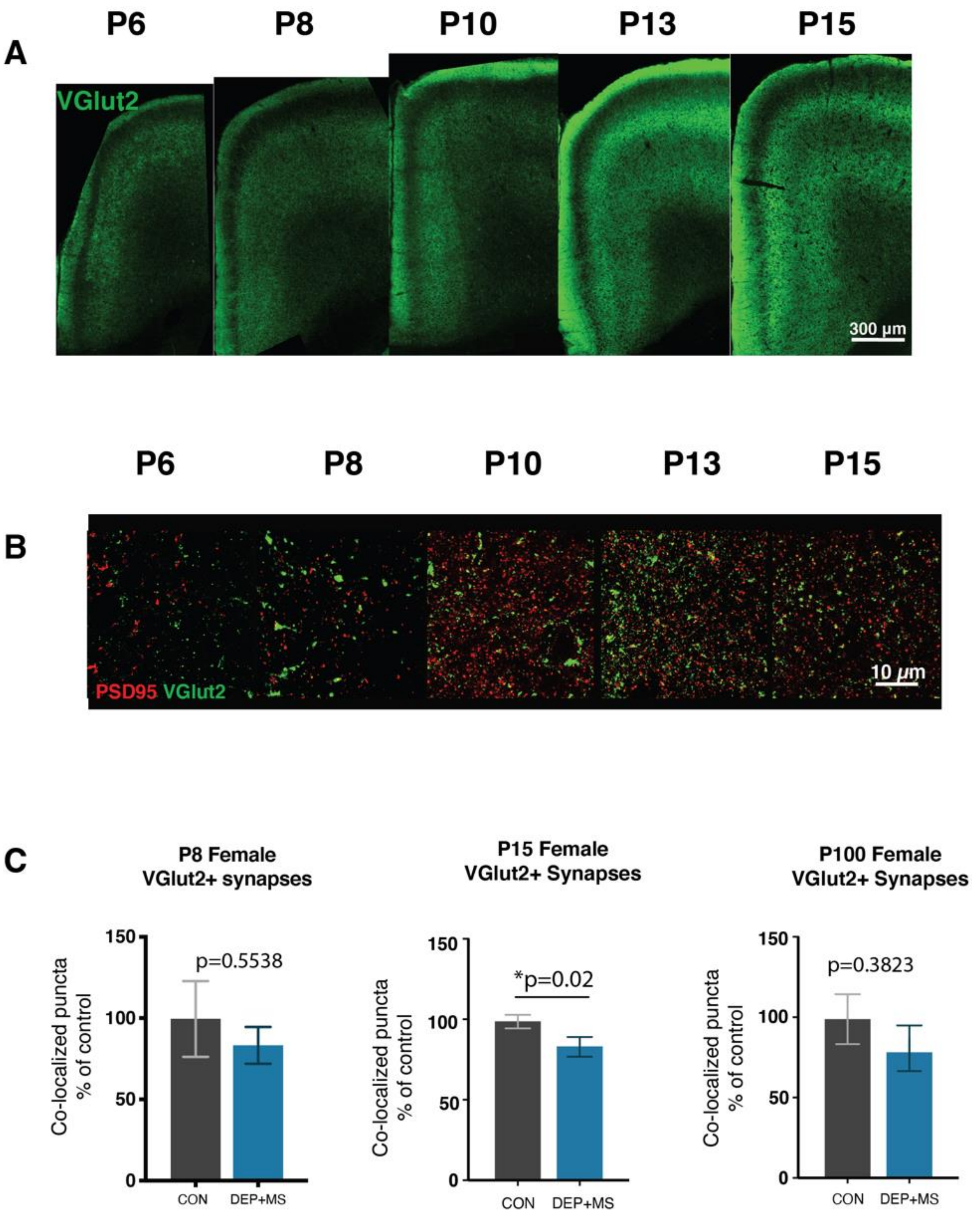
A
P6
P8
P10
P13
P15
VGlut2
300 µm
B
P6
P8
P10
P13
P15
PSD95 VGlut2
10 µm
C
P8 Female
VGlut2+ synapses
p=0.5538
Co-localized puncta
% of control
150
100
50
0
CON
DEP+MS
P15 Female
VGlut2+ Synapses
*p=0.02
Co-localized puncta
% of control
150
100
50
0
CON
DEP+MS
P100 Female
VGlut2+ Synapses
p=0.3823
Co-localized puncta
% of control
150
100
50
0
CON
DEP+MS

**Fig. S8 Thalamocortical synapses are increasing and become organized during the first and second postnatal week of life.** (**A**) Representative z-stack tile scans of VGlut2+ inputs in the Anterior Cingulate Cortex of untreated C57BL6/J mice from P6-P15. (**B**) Representative images of thalamocortical synapses labeled with VGlut2 and PSD95 in layer 1 of the ACC from untreated C57BL6/J mice P6-P15. (**C**) Quantification of TC synapse density in CON and DEP+MS females, TC synapse density is only transiently diminished at P15 in female DEP+MS offspring (n=3 animals/condition; 3 replicates/animal). Nested t-test (C). *$P < 0.05$;

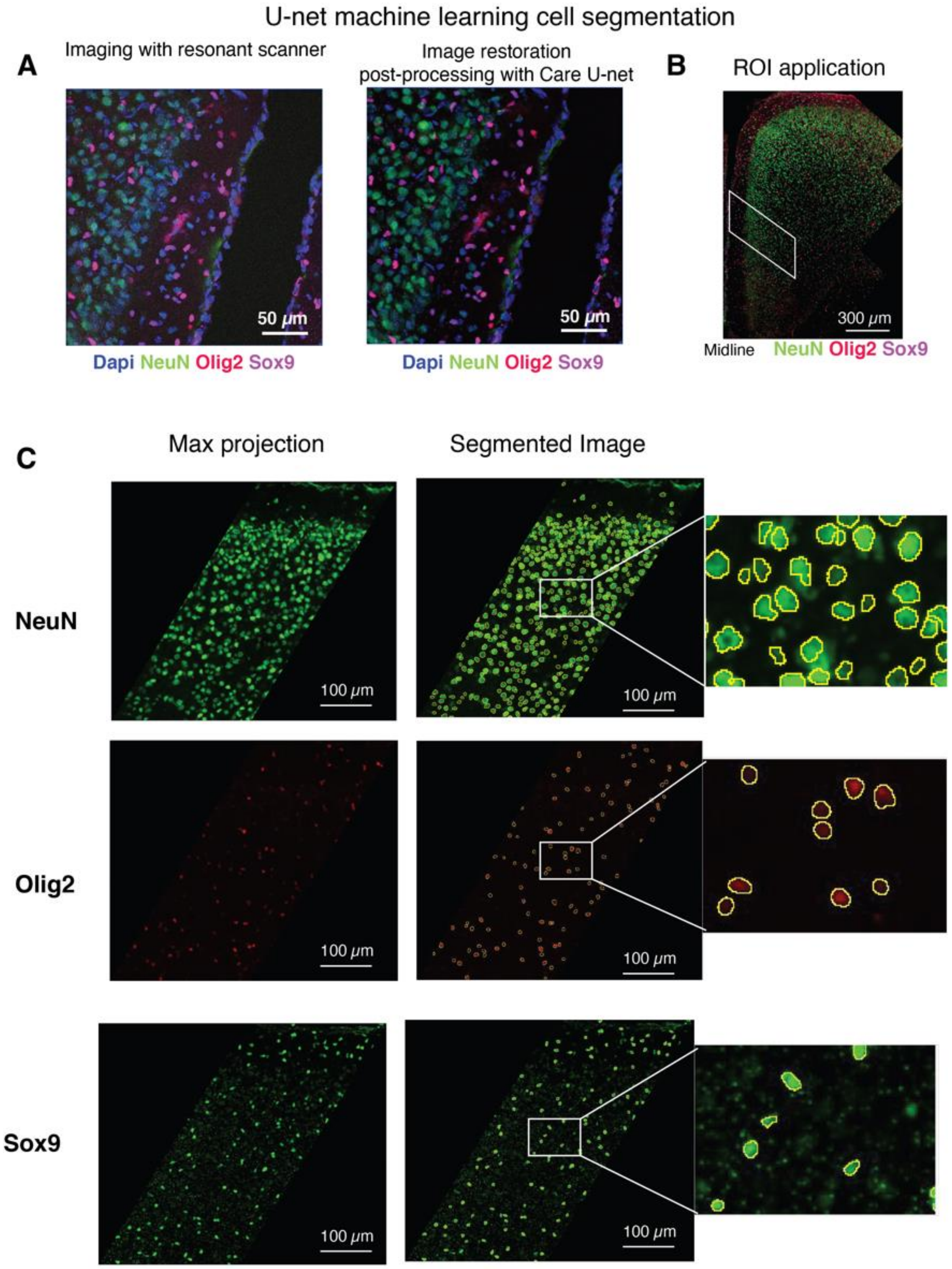
U-net machine learning cell segmentation
A
Imaging with resonant scanner
Image restoration
post-processing with Care U-net
50 μm
Dapi NeuN Olig2 Sox9
B
ROI application
300 μm
Midline
NeuN Olig2 Sox9
C
Max projection
Segmented Image
NeuN
Olig2
Sox9
100 μm

**Fig. S9. U-net image restoration and cell segmentation.** (**A**) To maximize acquisition speed, content-aware image restoration (CARE) u-net was used to restore lower resolution images acquired with resonant scanner. IHC was used to label neurons (NeuN), oligodendrocytes (Olig2) and astrocytes (Sox9). (**B**) Tile scan z-stack images of ACC were acquired, restored and a standard ROI was placed for cell quantification. (**C**) Segmentation models were trained for each marker, (left) original image of max projected restored tile scans, (right) segmentation model performance, yellow outlines generated by u-net models, model accuracy was verified for each image, and any errors were manually corrected.

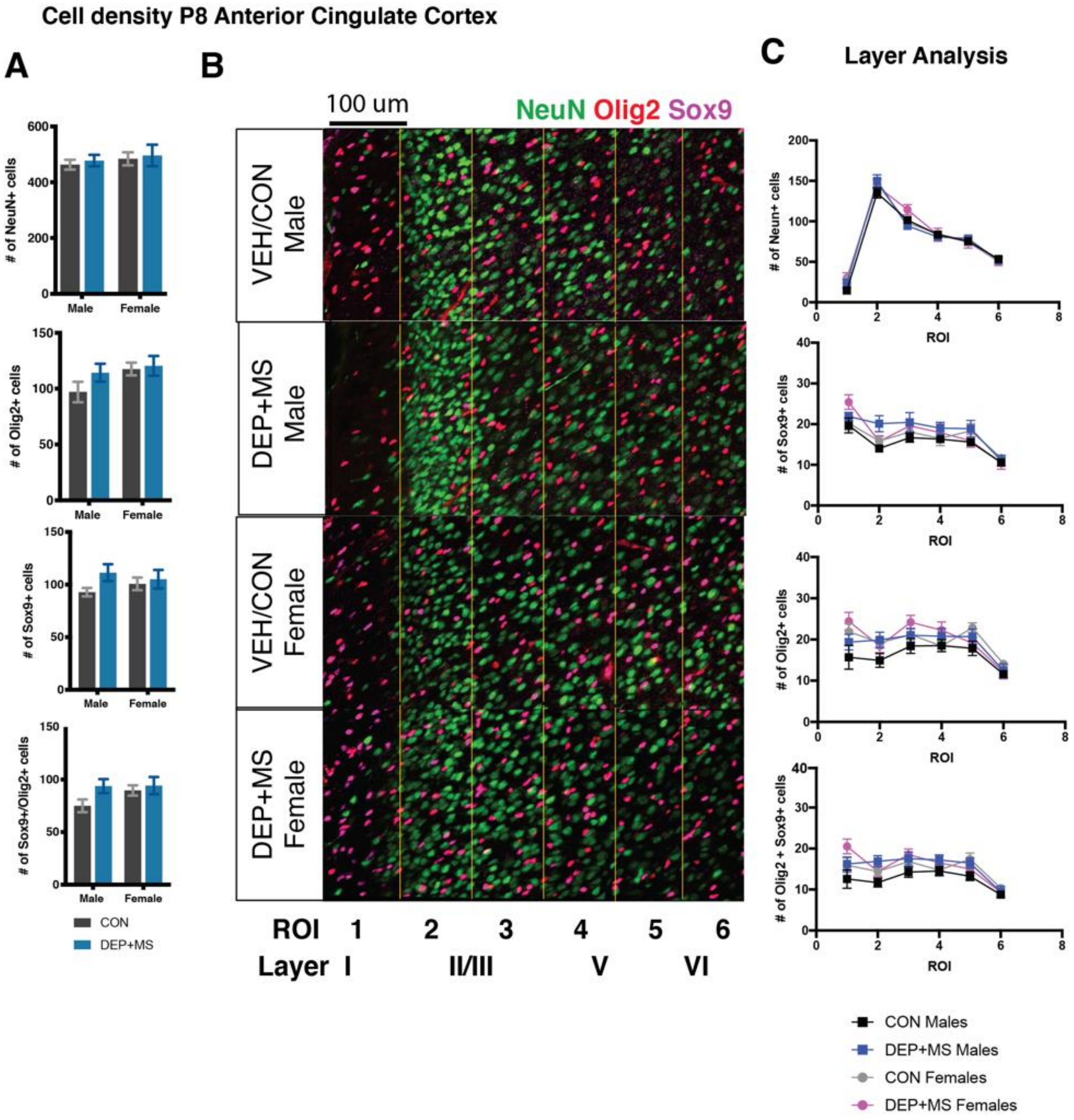

**Fig. S10. Density of neurons, oligodendrocytes and astrocytes unchanged in P8 ACC.** (**A**) Quantification of neurons (NeuN+), oligodendrocytes (Olig2+), astrocytes (Sox9+) and cells double positive for Olig2 and Sox9 using u-net (n=3 mice/condition/sex, n=3 biological replicates/mice), no significant differences in density of neurons, oligodendrocytes or astrocytes at P8. (**B**) Representative images of the ACC, a standard bin size of 160 microns was applied across image to determine differences in distribution. ROI's correspond roughly to cortical layers labeled below. (**C**) No significant differences in the distribution of cells across binned ROIs. Two-way ANOVA, condition x sex (A), ANCOVA (C). Means ± SEM.

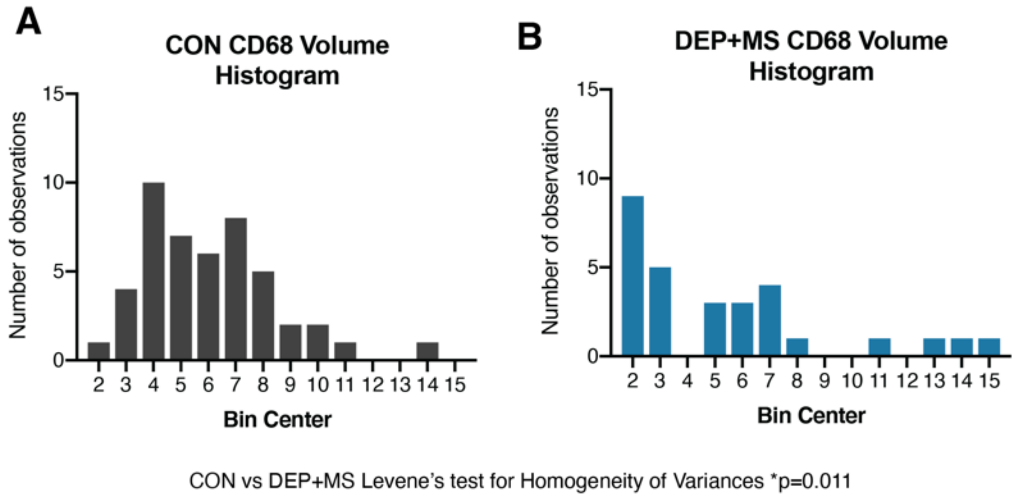

**Fig. S11 Distribution of the volume of CD68 is altered in DEP+MS male microglia. (A**) histogram of CD68 content within P10 microglia from CON males. (**B**) Histogram of CD68 content within P10 microglia from DEP+MS males. Levene's test for homogeneity of variances (CON vs DEP+MS).

A

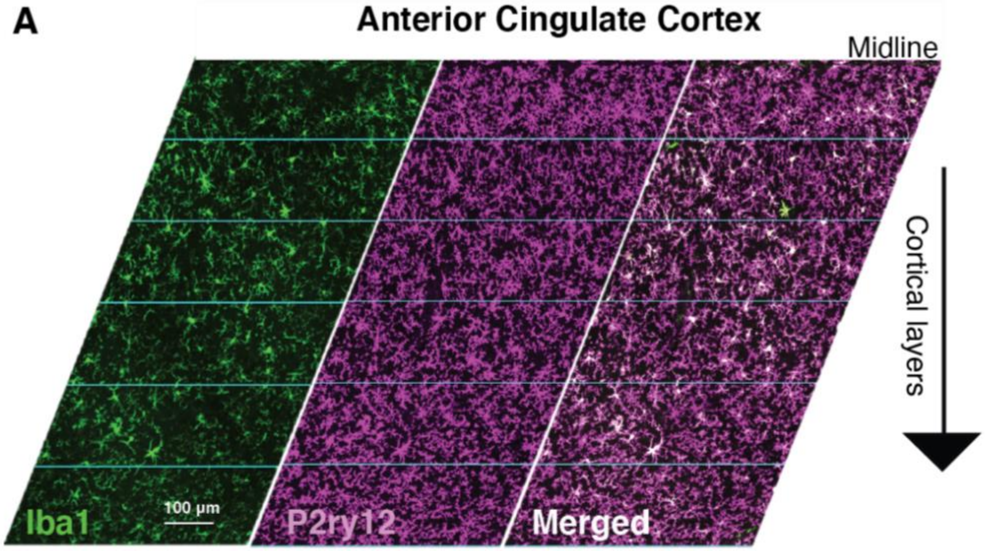

B

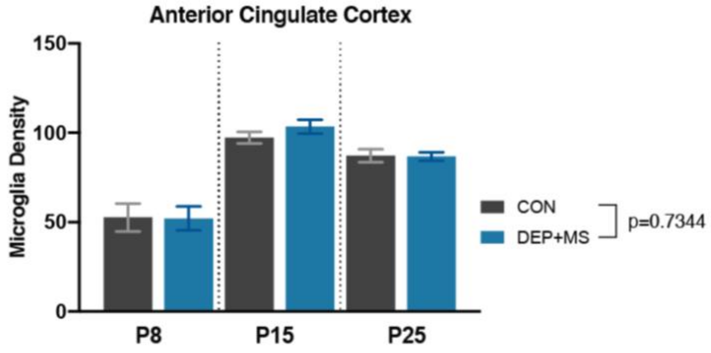

**Fig. S12 Microglial density unchanged across development in the ACC of DEP+MS offspring. (A)** Microglia were labeled with Iba1 and P2ry12, tile scan images were acquired, and an ROI and pseudo layering was applied to the ACC for microglia density quantification. (**B**) Microglia cell density varies by age, but there are no significant group differences in microglia density between CON and DEP+MS males at P8, P15 or P25 (n=3 mice/condition/age). Two-way ANOVA, condition x age (B). Means ± SEM.

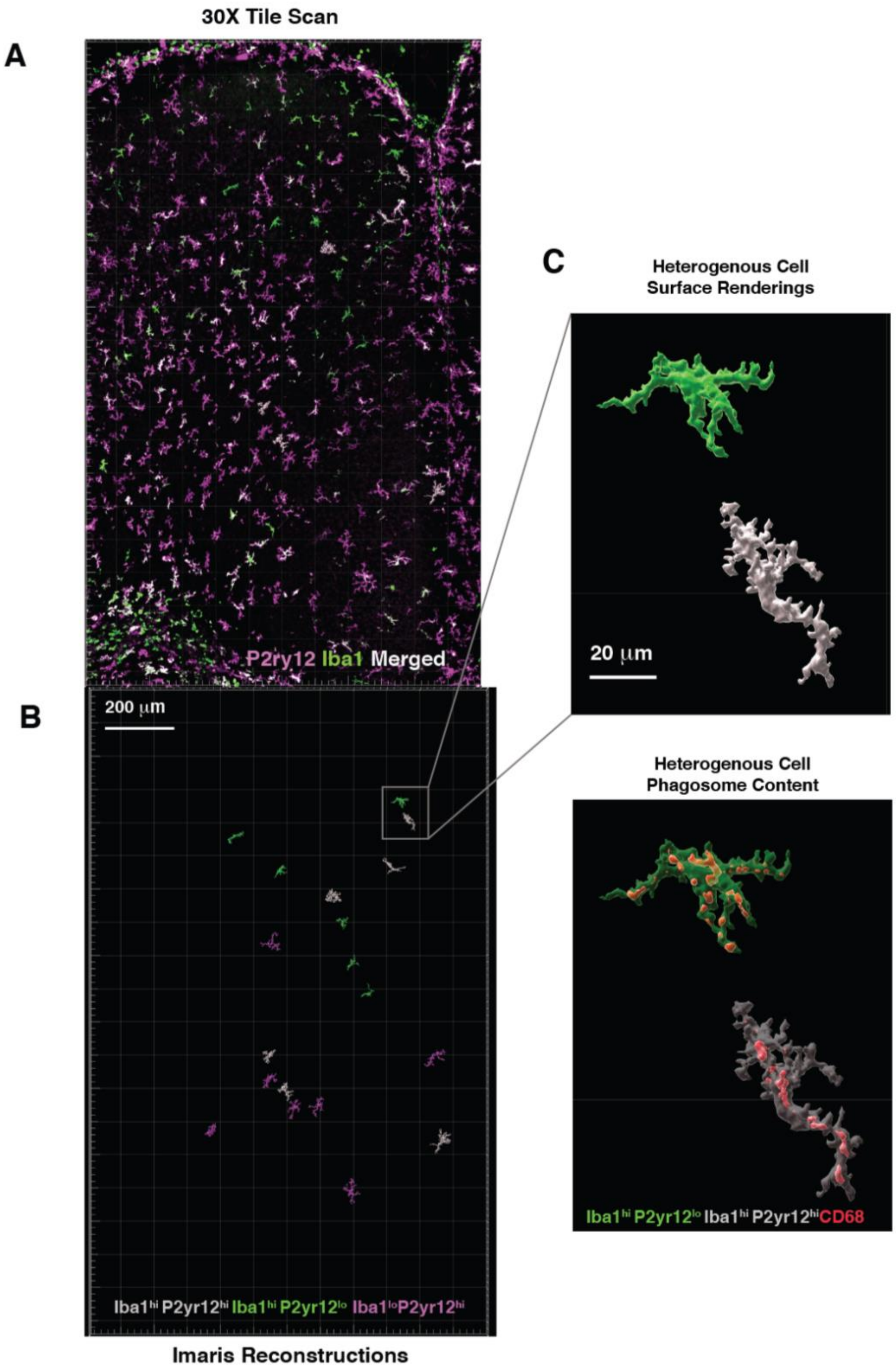
30X Tile Scan
A
P2ry12 Iba1 Merged
B
200 μm
Iba1hi P2yr12hi Iba1hi P2yr12lo Iba1loP2yr12hi
Imaris Reconstructions
C
Heterogenous Cell
Surface Renderings
20 μm
Heterogenous Cell
Phagosome Content
Iba1hi P2yr12lo Iba1hi P2yr12hiCD68

**Fig. S13 Imaris quantification of CD68 within microglia surface.** (**A**) Tile scan z-stack images of the ACC labeled for microglia (Iba1, P2ry12) and CD68 were acquired with a 30x objective. (**B**) Microglia were identified by differential expression of surface markers ($Iba1^{hi}P2ry12^{hi}$ (gray), $Iba1^{hi}P2ry12^{lo}$(green), $Iba1^{lo}P2ry12^{hi}$(magenta)) and 3-D surface renderings were generated using Imaris software. (**C**) CD68 surfaces within microglia were rendered with Imaris (n=3 mice/condition, 5-8 cells/subtype/mouse for a total ~20 cells/mouse, n=120 cells total).

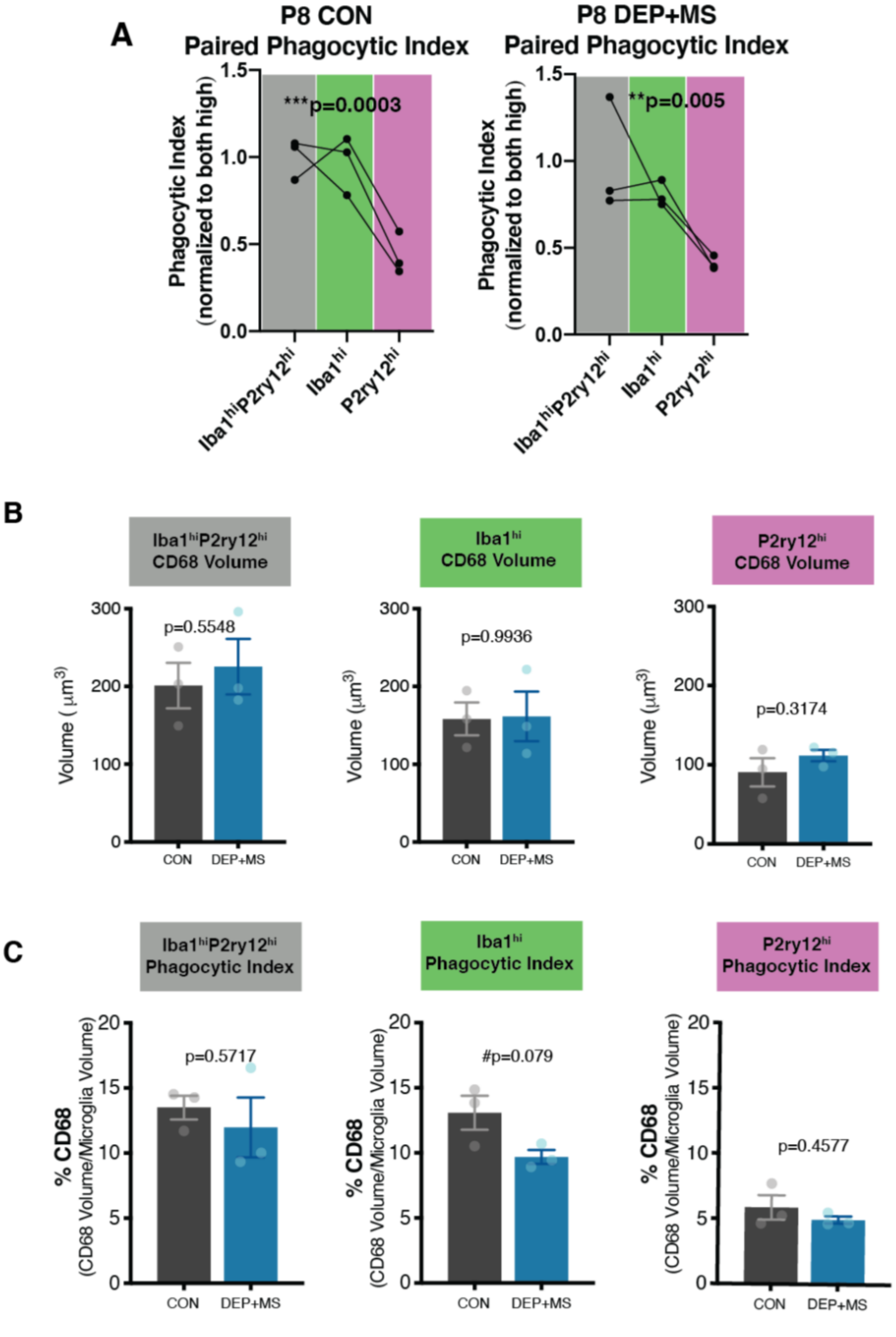
A
P8 CON
Paired Phagocytic Index
***p=0.0003
Phagocytic Index (normalized to both high)
1.5
1.0
0.5
0.0
Iba1hiP2ry12hi
Iba1hi
P2ry12hi
P8 DEP+MS
Paired Phagocytic Index
**p=0.005
B
Iba1hiP2ry12hi CD68 Volume
Iba1hi CD68 Volume
P2ry12hi CD68 Volume
Volume (μm3)
300
200
100
0
p=0.5548
p=0.9936
p=0.3174
CON
DEP+MS
C
Iba1hiP2ry12hi Phagocytic Index
Iba1hi Phagocytic Index
P2ry12hi Phagocytic Index
% CD68 (CD68 Volume/Microglia Volume)
20
15
10
5
0
p=0.5717
#p=0.079
p=0.4577

**Fig. S14 Functional differences in microglia phagocytic capacity present in both CON and DEP+MS microglia. (A)** Decreased phagocytic capacity of $Iba1^{lo}P2ry12^{hi}$ cells present in both CON and DEP+MS male microglia at P8 (n=20 cells/subtype/condition). **(B)** CD68 volume within each microglial subtype does not differ by condition (n=18-21 cells/subtype/condition, n=3 animals/condition). (**C**) Phagocytic index by microglial subtype is not significant different by condition, although there is a trend towards reduced phagocytic index in DEP+MS microglia from the $Iba1^{hi}$ group (n=18-21cells/subtype/condition). RM one-way ANOVA with Holm-Sidak's post hoc (A), Nested t-test (B, C). ***$P$<0.001, **$P$ < 0.01; *$P$ < 0.05; # P<0.10. Means ± SEM.

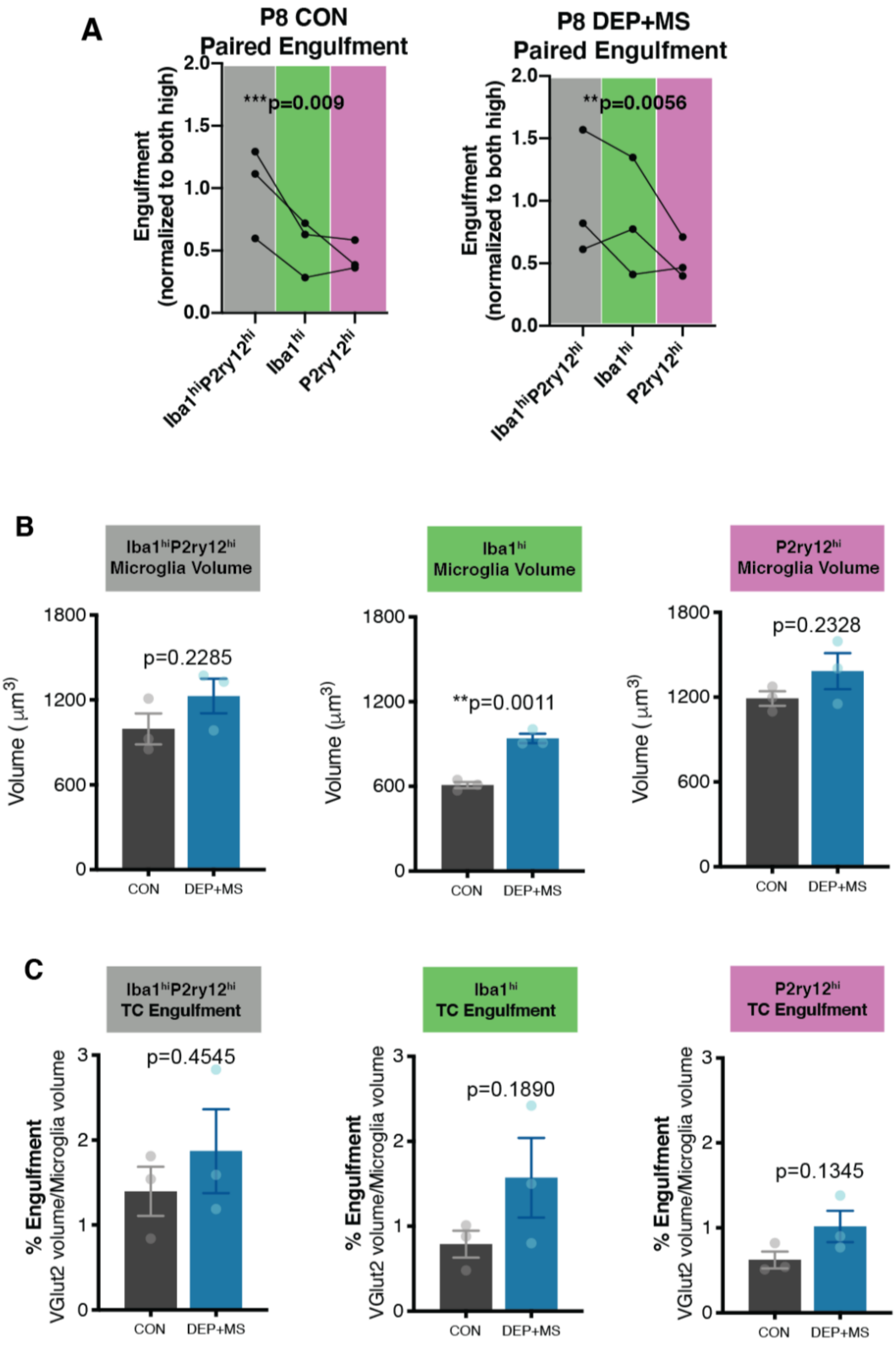
A
P8 CON
Paired Engulfment
***p=0.009
Engulfment
(normalized to both high)
Iba1hiP2ry12hi
Iba1hi
P2ry12hi
P8 DEP+MS
Paired Engulfment
**p=0.0056
B
Iba1hiP2ry12hi
Microglia Volume
Iba1hi
Microglia Volume
P2ry12hi
Microglia Volume
p=0.2285
**p=0.0011
p=0.2328
Volume (μm3)
CON
DEP+MS
C
Iba1hiP2ry12hi
TC Engulfment
Iba1hi
TC Engulfment
P2ry12hi
TC Engulfment
p=0.4545
p=0.1890
p=0.1345
% Engulfment
VGlut2 volume/Microglia volume

**Fig. S15 Functional differences in microglia engulfment of VGlut2 present in both CON and DEP+MS microglia. (A)** $Iba1^{lo}P2ry12^{hi}$ cells have significantly reduced VGlut2 engulfment in both CON and DEP+MS P8 microglia (n=20-40 cells/subtype/condition, n=3 animals/group). **(B)** Microglia volume of $Iba1^{hi}$ cells is significantly increased in DEP+MS male microglia, no significant volume changes in $Iba1^{hi}P2ry12^{hi}$ or $P2ry12^{hi}$ cells by condition (n=20-40 cells/subtype/condition, n=3 animals/group). (**C**) Engulfment of VGlut2 inputs within microglia subtypes does not differ significantly between prenatal treatments (n=20-40 cells/subtype/condition, n=3 animals/group). RM one-way ANOVA with Holm-Sidak's post hoc test (A), Nested t-test (B, C). $^{**}P < 0.01$; $^{*}P < 0.05$. Means ± SEM.

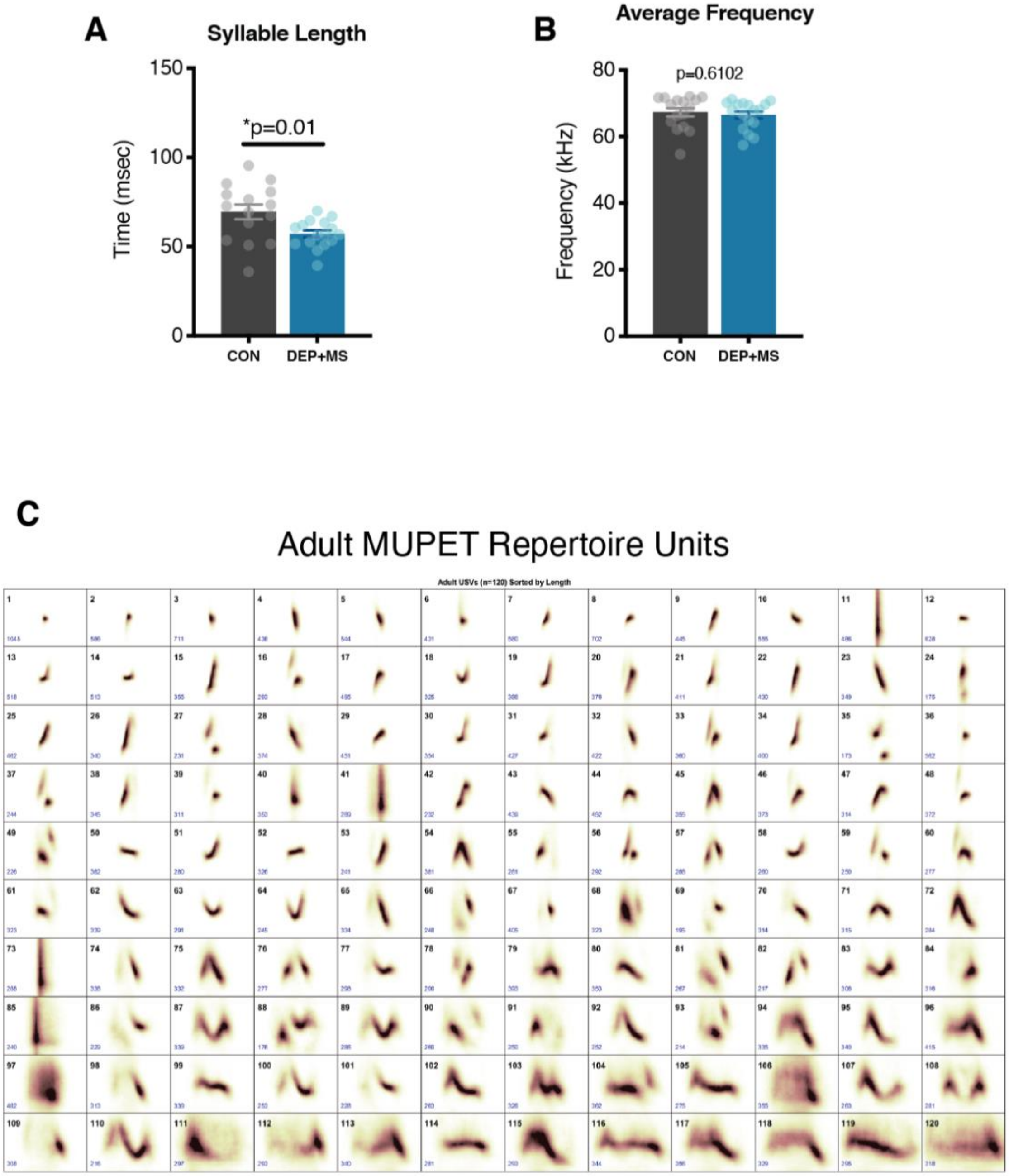

**Fig. S16 Adult USVs in males prenatally exposed to CON or DEP+MS. (A)** Length of USV syllable is significantly decreased in adult DEP+MS mice (n=15-17 mice/condition). **(B)** No significant differences in the frequency of USV emitted by adult males (n=15-17 mice/condition). (**C**) Adult USV MUPET repertoire unit (RU) clustering, organized from shortest to longest, compiled from all CON and DEP+MS USV files. Unpaired t-test (A-B). *$P$ < 0.05. Means ± SEM.

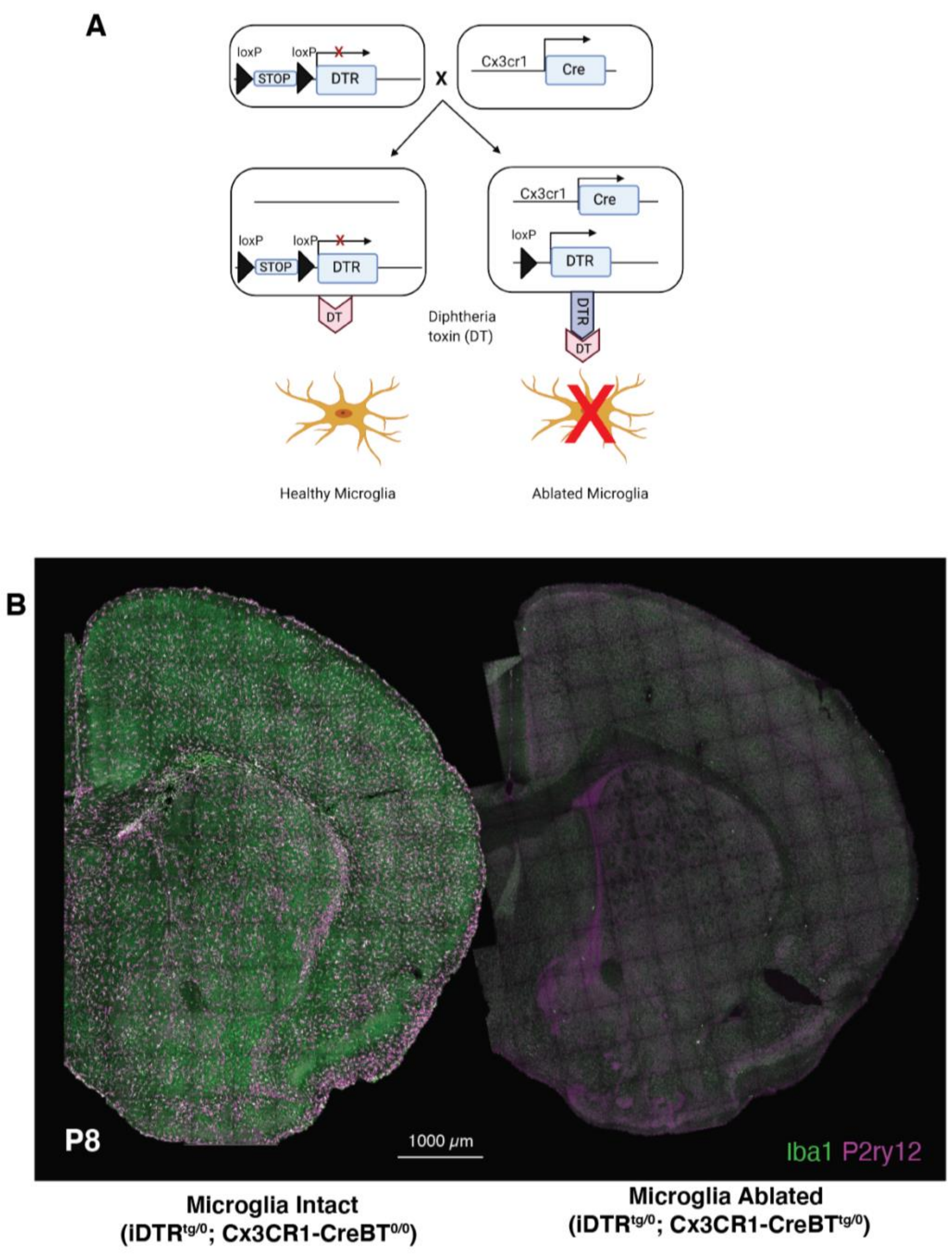

**Fig. S17 Genetic strategy to deplete microglia. (A)** Genetic strategy to render microglia cells susceptible to diphtheria toxin (DT). Crossing the iDTR strain to Cx3cr1-BTCre$^{tg/0}$ removes STOP cassette inducing expression of diphtheria toxin receptor (DTR, right). Microglia cells not expressing DTR remain intact when exposed to DT (left), whereas microglia expressing DTR undergo apoptosis when exposed to DT (right). **(B)** i.p. injection of DT exposure at P5 and P7, rapidly eliminates microglia by P8.

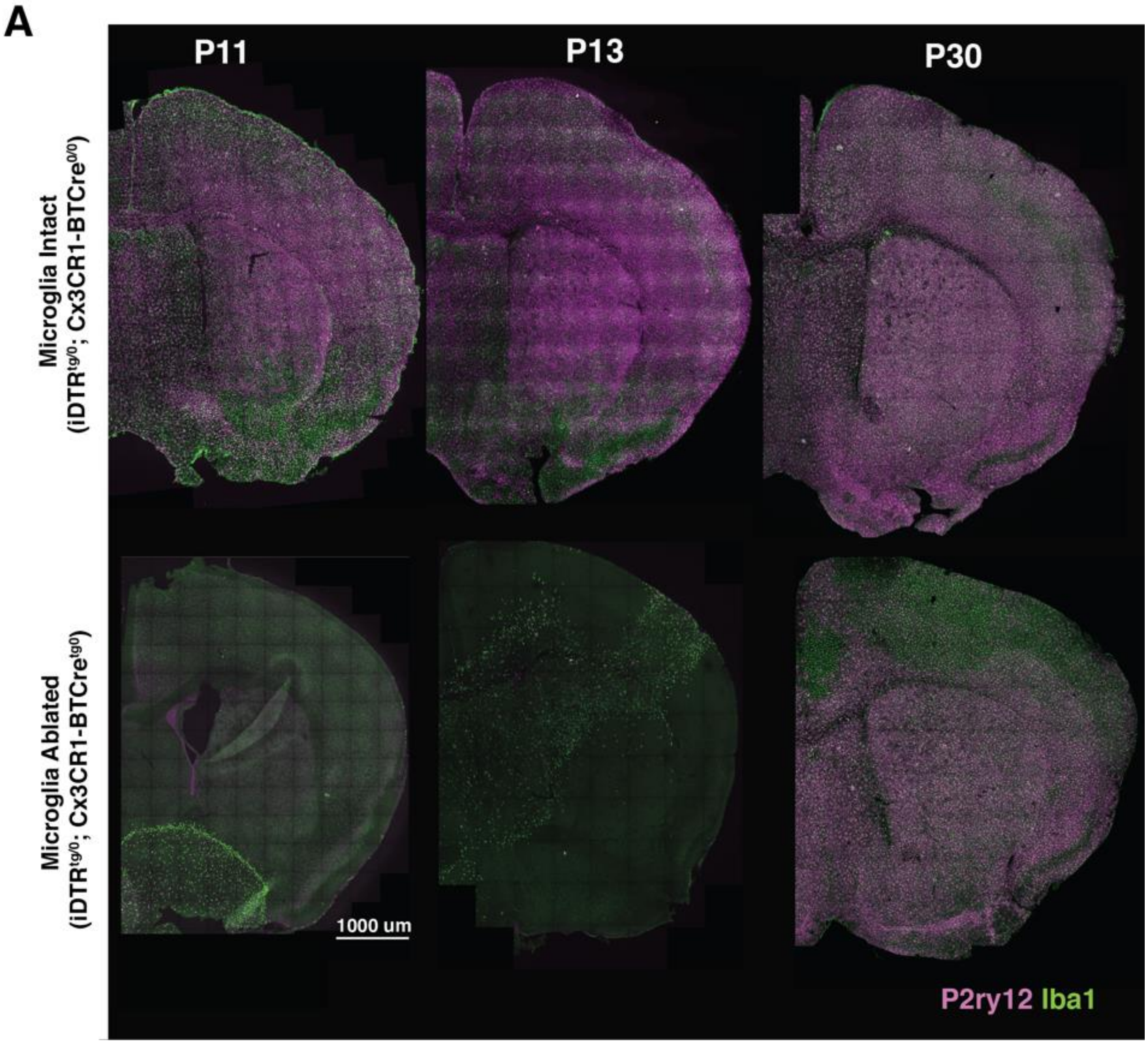

**Fig. S18. Microglia repopulate the brain by P30 but remain heterogenous.** (**A**) Coronal brain sections from mice with either intact or ablated microglia at P11, P13 and P30 (ablation occurred at P5-P7). At P11 and P13, $Iba1^{hi}$ microglia repopulate the brain. By P30, microglia repopulation is largely complete; however, cortical microglia still do not express high levels of homeostatic marker P2ry12 (bottom right).
